# CellNeighborEX: Deciphering Neighbor-Dependent Gene Expression from Spatial Transcriptomics Data

**DOI:** 10.1101/2022.02.16.480673

**Authors:** Hyobin Kim, Cecilia Lövkvist, António M. Palma, Patrick Martin, Junil Kim, Amit Kumar, Maria Leonor Peixoto, Esha Madan, Rajan Gogna, Kyoung Jae Won

## Abstract

Cells have evolved communication methods to sense their microenvironments and send biological signals. In addition to the communication using ligands and receptors, cells use diverse channels including gap junctions to communicate with their immediate neighbors. Current approaches, however, cannot effectively capture the influence of various microenvironments. Here, we propose a novel approach that identifies *cell neighbor*-dependent gene *ex*pression (CellNeighborEX). After categorizing cells based on their microenvironment from spatial transcriptomics (ST) data, CellNeighborEX identifies diverse gene sets associated with partnering cell types, providing further insight. To categorize cells along with their environment, CellNeighborEX uses direct cell location or the mixture of transcriptome from multiple cells depending on the ST technology. We show that cells express different gene sets depending on the neighboring cell types in various tissues including mouse embryos, brain, and liver cancer. These genes were associated with development (in embryos) or metastases (liver cancer). We further validate that gene expression can be induced by neighboring partners. The neighbor-dependent gene expression suggests new potential genes involved in cell-cell interactions beyond what ligand-receptor co-expression can discover.

## Introduction

Cells communicate with their microenvironment in various ways including the release of soluble molecules and direct cell contact (Yang *et al*, 2021), actively changing their transcriptomes in response to external signals (Cable *et al*, 2021; Fischer *et al*, 2023). To gain insights into critical biological processes such as diseases and development, it is essential to understand the various ways of cell-cell communication. Experimental approaches to studying cell-cell communication usually require an elaborate and intricate setup (Nishida-Aoki & Gujral, 2019).

Genome-scale study on cell-cell interactions has been recently possible by using ligand-receptor co-expression on single cell RNA-sequencing (scRNA-seq) (Browaeys *et al*, 2020; Efremova *et al*, 2020) and spatial transcriptomics (ST) data (Garcia-Alonso *et al*, 2021; Li *et al*, 2021; Pham *et al*, 2020; Shao *et al*, 2022). The use of ligand-receptor co-expression enabled inferring interacting cell type pairs and identifying intercellular signaling pathways without relying on a complicated experimental setup. However, it cannot elucidate the gene expression of individual cells changed by direct cell contact.

A growing body of studies on cell communication have demonstrated that cells are influenced by their microenvironment and neighboring cells (Barone *et al*, 2017; Hannezo & Heisenberg, 2019). Grafting experiments in developing embryos manifested that direct cell contact can induce signals for the development of a specific tissue type (Solini *et al*, 2017; Spemann & Mangold, 2003). More recently, RNA sequencing of physically interacting cells (PIC-seq) has revealed that cells express genes depending on neighboring cell types during mouse development (Kim *et al*, 2023). This study suggests that cells have distinct expression profiles through direct cell contact independently from ligand-receptor-mediated communication.

Recent development in ST has opened potential ways to explore the role of the microenvironment. The spatial gene expression profile has made it possible to study the transcriptional activity of a cell together with that of the neighborhood within intact tissues. There are largely two types of ST data with their own advantages and limitations. Image-based approaches including MERFISH (Chen *et al*, 2015) and seqFISH (Eng *et al*, 2019; Lubeck *et al*, 2014) use fluorescence *in situ* hybridization (FISH) to visualize the RNA species of interest. While image-based ST approaches can quantify the RNAs at cellular resolution, the number of detectable RNA species is still limited. The next-generation-sequencing (NGS) based ST approaches such as Visium (Ståhl *et al*, 2016) and Slide-seq (Rodriques *et al*, 2019; Stickels *et al*, 2021; Zhao *et al*, 2022) leverage spatially barcoded beads. While NGS-based approaches can unbiasedly profile the transcriptome, a barcode can be linked to the mixture of transcriptome of multiple cells or cell portions depending on the position and the resolution of the barcoded spots, making it hard to detect gene expression changed by the cellular microenvironment.

Many computational tools have been developed to understand cell-cell interactions from ST data. CellphoneDB v.3.0 (Garcia-Alonso *et al*., 2021), MESSI (Li *et al*., 2021), SpaTalk (Shao *et al*., 2022), and stLearn (Pham *et al*., 2020) use the co-expression of ligand-receptor pairs to study cellular communication. However, ligand-receptor co-expression cannot completely capture cell-cell interactions due to direct contact. SVCA (Arnol *et al*, 2019) decomposes the sources of gene expression variation into intrinsic effects, environmental effects, and cell-cell interactions. It explains the relationship between gene expression and cell-cell interactions. However, SVCA does not have a function to detect gene expression change associated with cell contact, and their strategy has been only optimized for image-based ST data. MISTy (Tanevski *et al*, 2022) quantifies the contributions of different spatial contexts to the expression of markers of interest. The influence of immediate neighborhoods on the expression of markers can be investigated. However, MISTy requires to pre-select the list of marker genes to find potential interactions and it has not been designed to identify gene expression change related to cell contact in an unbiased way. DeepLinc (Li & Yang, 2022) reconstructs a cell interaction network from ST data. Regarding three nearest neighbors as direct contact, DeepLinc finds signature genes contributing to interactions between cell types and infers proximal interactions between them. However, it does not uncover specific relationships between the signature genes and interacting cell types. C-SIDE (Cable *et al*, 2022) examines up- and down-regulated genes depending on proximity to a certain cell type. Because the interaction between cell types is defined based on cell density rather than cell contact, C-SIDE does not work for studying cell contact-dependent gene expression. NCEM (Fischer *et al*., 2023) investigates transcriptomic change depending on local environments but it has not been designed to study the influence of cell contact on gene expression, particularly for NGS-based data. NCEM considers one barcoded-spot a single cell type even in low resolution Visium data, so it does not look into the influence of direct contact between multiple cell types within one spot. Although spatial context has been applied to study cell-cell interactions, transcriptomic change associated with cell contact has not been fully explored yet. It is still challenging to detect genes influenced by cell contact regardless of the data types.

Here, we propose a universal approach called CellNeighborEX to identify genes influenced by neighboring cells from ST data. CellNeighborEX dissects the transcriptome of cells with their immediate neighbors to categorize cells based on the neighboring cell types. For NGS-based ST data where exact cell locations are not available, CellNeighborEX actively uses the mixture of transcriptomes to identify immediate neighbors. CellNeighborEX was applied to various ST data from mouse embryos, hippocampus, and liver cancer to identify neighbor-dependent genes. These transcriptomic changes were confirmed in the spatial context. We showed that cells express specific genes depending on neighboring cell types. The neighbor-dependent gene expression suggested new potential genes involved in cell-cell interactions beyond what ligand-receptor co-expression can discover and gave clues on understanding complex biological processes.

## Results

### CellNeighborEX categorizes transcriptome to investigate the influence of neighboring cell types

CellNeighborEX defines immediate neighbors differently for image- and NGS-based ST data. In image-based ST data where exact cell locations are available, CellNeighborEX finds the nearest neighbors using Delaunay triangulation (Delaunay, 1934). The immediate neighbors can also be defined using other algorithms such as radial distance, or k-nearest neighbors (KNN)(Cover & Hart, 1967; Fix & Hodges, 1951) (Fig. 1 A). CellNeighborEX classifies cells into two groups based on the cell types of the nearest neighbors for each cell type. *Homotypic neighbors* consist of the same cell type. *Heterotypic neighbors* are composed of different cell types. CellNeighborEX investigates the influence of neighboring cell types by comparing the transcriptome of heterotypic neighbors with that of homotypic neighbors (log2(fold-change) >0.4, p-value < 0.01, FDR < 0.05). Parametric (*i*.*e*., Student’s t-test, Welch’s t-test) or non-parametric (*i*.*e*., Mann-Whitney U test) statistical tests are performed as differential expression (DE) analysis depending on the sample size of the groups (Methods).

**Fig. 1.**
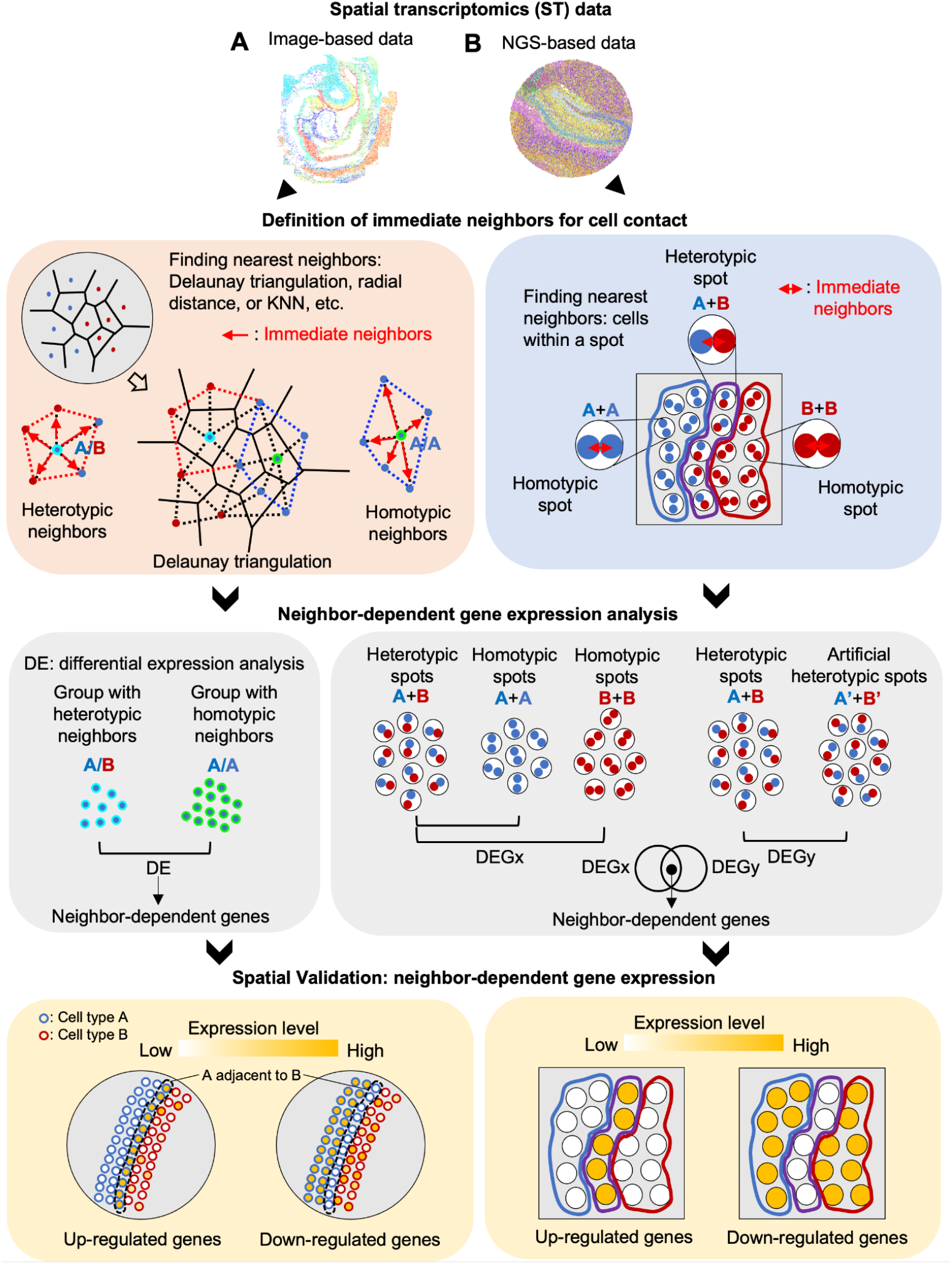
Workflow of CellNeighborEX. **A.** In image-based ST data, the immediate neighbors for cell contact are determined using algorithms such as Delaunay triangulation, radial distance, and KNN. Based on their cell types, homotypic and heterotypic neighbors are defined. CellNeighborEX detects genes influenced by neighbors by comparing the transcriptome from heterotypic neighbors with those from homotypic neighbors. **B**. In NGS-based ST data, there are homotypic spots (the same cell type in a bead) and heterotypic spots (multiple cell types in a bead). The heterotypic spots are regarded as evidence for cell contact. CellNeighborEX compares the heterotypic spots with the homotypic ones to detect neighbor-dependent genes. Additional statistical tests with a null model are applied for validation.

The locations of cells are not explicitly given for NGS-based ST approaches and a barcoded spot contains the mixed transcriptome from multiple cells. CellNeighborEX capitalizes on spots with multiple cell types (or heterotypic spots) as the evidence for cell contact (Fig. 1 B). This strategy is effective in studying neighbor-dependent gene expression when the diameter of a spot is near cellular resolution. For instance, Slide-seq, a ST approach with 10 μ*m* resolution, has *homotypic spots* (the same cell type in a bead) and *heterotypic spots* (two or more cell types in a bead), and the majority of the heterotypic spots are composed of two cell types (97% of spots are composed of one or two cell types) (Rodriques *et al*., 2019). We use deconvolution tool RCTD (Cable *et al*., 2021) to find heterotypic and homotypic spots. Next, the transcriptome of heterotypic spots is compared with the transcriptome of homotypic spots to identify neighbor-dependent genes (log2(fold-change) >0.4, p-value < 0.01). To confirm the statistical significance of the identified neighbor-dependent genes, CellNeighborEX generates a null model of artificial heterotypic spots (FDR < 0.01) (Methods, Fig. S1).

### CellNeighborEX identifies neighbor-dependent genes related with embryonic development from seqFISH data in a mouse embryo

We applied CellNeighborEX to seqFISH data from a mouse embryo. We used predefined cell type annotation provided with the seqFISH data (Lohoff *et al*, 2022). After defining immediate neighbors using Delaunay triangulation (Delaunay, 1934), CellNeighborEX categorized cells based on neighboring cell types. After the DE analysis, CellNeighborEX detected 354 up-regulated genes from 22 types of heterotypic neighbors and 429 down-regulated genes from 22 types of heterotypic neighbors (Table S1A, Table S1B).

For instance, we found 694 Gut tube cells surrounded by the same cell type (homotypic neighbors) and 33 Gut tube cells adjacent to Neural crest cells among other heterotypic neighbors. CellNeighborEX found that Gut tube cells express *Pitx1* when adjacent to Neural crest cells (p-value < 0.01 and FDR < 0.05) (Fig. 2A). To confirm this result, we visualized the expression of *Pitx1* for all Gut tube cells together with Neural crest cells. The Gut tube cells adjacent to Neural crest (Gut tube/Neural crest) are represented in red boundaries, while other Gut tube cells (Gut tube/Gut tube) are represented in blue boundaries. The Neural crest cells are shown in black boundaries. The rest of the cells are shown in gray (Fig 2B). The visualization demonstrated the clear expression of *Pitx1* in the Gut tube cells adjacent to Neural crest (Gut tube/Neural crest) but not in the Gut tube cells adjacent to other Gut tube cells (Gut tube/Gut tube) (Fig. 2B). The expression of *Pitx1* was previously observed in a patch of foregut endoderm that contacts the stomodeum (Lanctôt *et al*, 1997) with which neural crest contributes to facial development. Gene Ontology (GO) analysis for the 354 up-regulated genes were associated with biological terms such as “pattern specification process” and “embryonic organ morphogenesis” (Fig. 2C), suggesting that cell contact with other cell- or tissue-type may regulate the coordinated developmental processes.

**Fig. 2.**
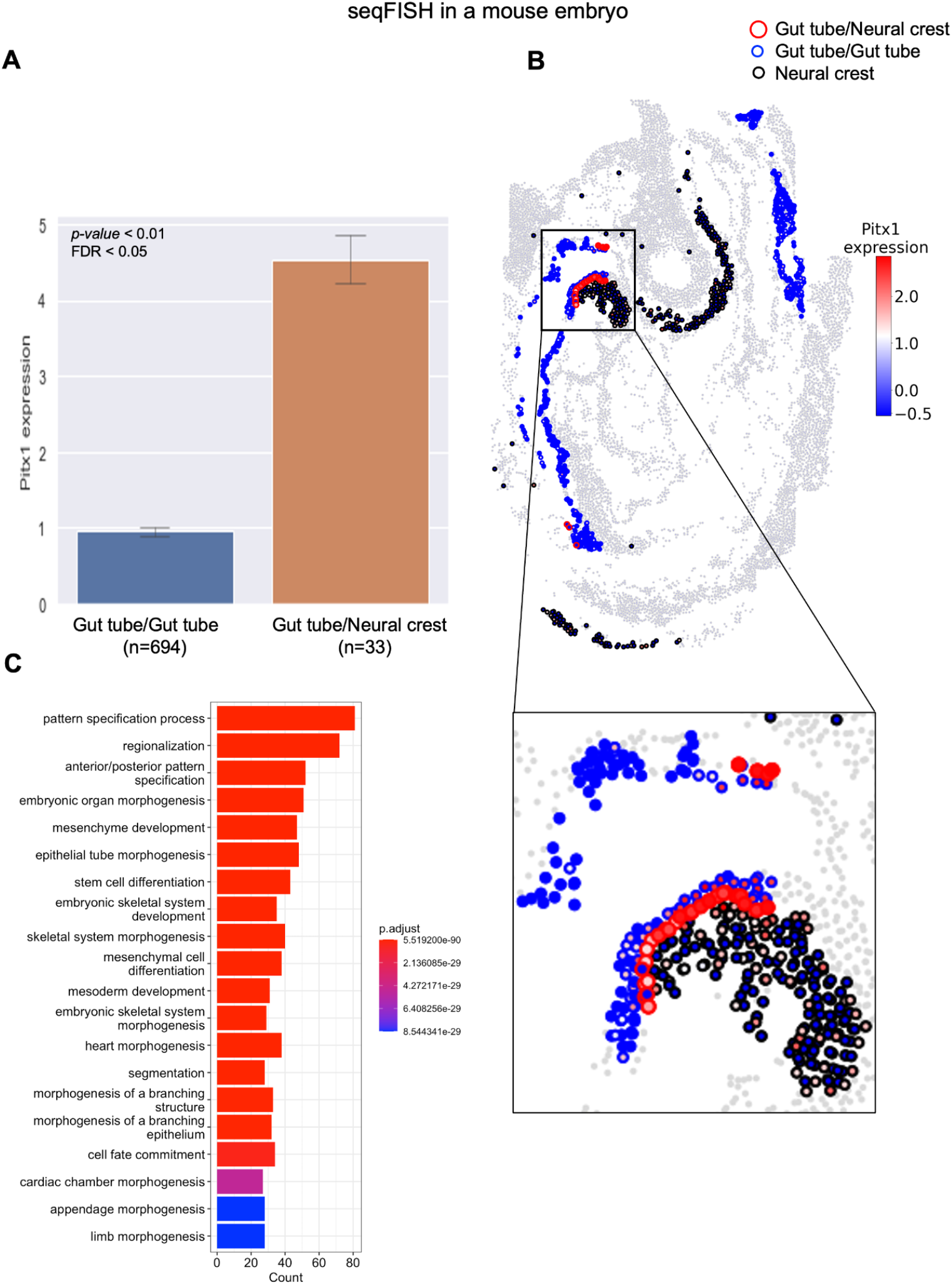
Neighbor dependent-genes identified by CellNeighborEX from the mouse embryo seqFISH data. **A.** *Pitx1* is more highly expressed in Gut tube cells proximal to Neural crest cells (Gut tube/Neural crest) than Gut tube cells proximal to other Gut tube cells (Gut tube/Gut tube). **B**. The spatial visualization displays that Gut tube cells adjacent to Neural crest more highly express *Pitx1*. **C**. For genes up-regulated by cell contact in the mouse embryo seqFISH data, GO analysis shows that the GO terms are associated with embryonic development.

### CellNeighborEX detects neighbor-dependent genes from Slide-seq data in a mouse embryo

We analyzed Slide-seq V2 data from a mouse embryo. For this NGS-based data, CellNeighborEX uses heterotypic beads to define immediate neighbors. To find heterotypic spots, we applied the RCTD (Cable *et al*., 2021) deconvolution tool using single cell RNA-sequencing (scRNA-seq) for a mouse embryo (Cao *et al*, 2019) as a reference. As a result, RCTD identified 8,094 homotypic and 34,268 heterotypic spots in the mouse embryo. The decomposed cell types of the heterotypic spots were additionally validated by the expression of cell type markers and correlation analysis (Methods, Fig. S2). Comparing the gene expression levels of heterotypic spots with the expression of homotypic spots, CellNeighborEX detected neighbor-dependent genes. Finally, CellNeighborEX found 28 up-regulated genes from 9 heterotypic pairs, and 28 down-regulated genes from 11 heterotypic pairs in the embryo (Table S1C, Table S1D).

For example, CellNeighborEX identified 17 genes including *Cd24a*, which are highly expressed in the heterotypic spots of Endothelial and Lens cells compared with their respective homotypic spots (Fig. S3A). The heterotypic spots expressed marker genes for both Endothelial (*Pecam1* and *Egfl7*) and Lens cells (*Cryba1* and *Cryaa*). We confirmed in the spatial visualization that *Cd24a* is more highly expressed in the heterotypic spots of Endothelial+Lens cells (red boundaries) compared with the homotypic spots of Endothelial (blue) and Lens (black) cells (Fig. S3B).

To find which cell type the expression of *CD24a* comes from between Endothelial cells and Lens cells, we used a regression model in which gene expression is shown along the proportion of each cell type in the heterotypic spots (Methods). We found that the expression level of *CD24a* increased as the proportion of Endothelial cells (against Lens cells) increased (Fig. S4A). We further confirmed that *Cd24a* is expressed mainly from Endothelial cells when examining scRNA-seq data from a mouse embryo (Cao *et al*, 2019) (Fig. S3C).

For the 28 up-regulated genes, the GO analysis showed terms related to tissue development including “eye development” and “digestive system development” (Fig. S3D), further suggesting the role of cell contact in the developmental processes.

### CellNeighborEX detects genes influenced by tumor microenvironments (TME) from Slide-seq data in mouse liver cancer

For Slide-seq data in mouse liver metastases, RCTD used single-nucleus RNA-sequencing (snRNA-seq) data in mouse liver cancer (Zhao *et al*., 2022) as a reference. It identified 16,557 homotypic and 6,284 heterotypic spots. After validating the cell type annotation of the heterotypic spots (Methods, Fig. S2), CellNeighborEX detected 42 up-regulated genes from 10 heterotypic pairs and 3 down-regulated genes from 2 heterotypic pairs (Table S1E, Table S1F).

For instance, CellNeighborEX found that *F13a1* is highly expressed when Monocyte cells contact Tumor cells (Fig. S5A). Its spatial mapping displays the higher expression in the heterotypic spots than the homotypic ones (Fig. S5B). Using the regression model and snRNA-seq data, we checked that the expression of *F13a1* is derived from Monocyte cells (Fig. S4B, Fig. S5C). Previous global proteomic analysis on small extracellular vesicles identified that *F13a1* is associated with liver cancers (Dong *et al*, 2022). Besides, *F13a1* promotes lung squamous cancer (Porrello *et al*, 2018) and is a biomarker for colorectal cancers (Peltier *et al*, 2018).

For the 42 up-regulated genes, the GO analysis showed terms such as “collagen fibril organization” and “endothelial cell migration” (Fig. S5D). Specifically, *Col1a2, Col3a1, Ext1*, and *Tgfbr1* are associated with collagen fibril organization. *Col1a2* and *Col3a1* are more expressed when Vascular smooth muscle cells contact Tumor III cells (VSMC+Tumor III). *Ext1* and *Tgfbr1* are more expressed when Tumor III cells contact Hepatocyte II cells (Tumor III+Hepatocyte II). These four genes are all highly expressed when non-tumor and tumor cells interact with each other. Based on previous studies showing that collagen influences cancer cell behaviors such as metastasis, tumorigenesis, and proliferation (Xu *et al*, 2019), we can infer that the four genes might affect liver cancer by controlling the collagen arrangement among VSMC, Hepatocyte, and Tumor cells. A vitro study actually showed that the down-regulation of *Col1a2* suppressed hepatocellular cancer (Ji *et al*, 2010), which supports that neighbor-dependent genes are indeed associated with critical biological processes. It is notable that CellNeighborEX can detect genes influenced by TME in an unbiased way.

### Neighbor-dependent genes are discovered from Slide-seq data in mouse brain

We also analyzed Slide-seq V2 data in the mouse brain. Using scRNA-seq for mouse hippocampus (Saunders *et al*, 2018) as a reference, we ran RCTD (Cable *et al*., 2021). In total, 12,013 homotypic and 29,331 heterotypic spots were identified in the mouse hippocampus. We additionally validated the decomposed cell types of the heterotypic spots from the expression of cell type markers and correlation analysis (Methods, Fig. S2). CellNeighborEX found 155 up-regulated genes from 21 heterotypic pairs and 55 down-regulated genes from 8 heterotypic pairs for this dataset (Table S1G, Table S1H).

For example, CellNeighborEX detected 3 up-regulated genes in the heterotypic spots of Endothelial tip (EnT) and Astrocyte cells including *Fabp7* (Fig. 3A). Its spatial visualization shows that *Fabp7* is more expressed in the heterotypic spots than the respective homotypic ones (Fig. 3B). From the scRNA-seq data taken from hippocampus (Saunders *et al*., 2018), we examined the potential cell types expressing these genes (Fig. 3C). For the 155 up-regulated genes, the GO analysis showed terms such as “regulation of metal ion transport” and “dopamine secretion”, suggesting that cell contact may contribute to neuronal processes in the mouse brain (Fig. 3D).

**Fig. 3.**
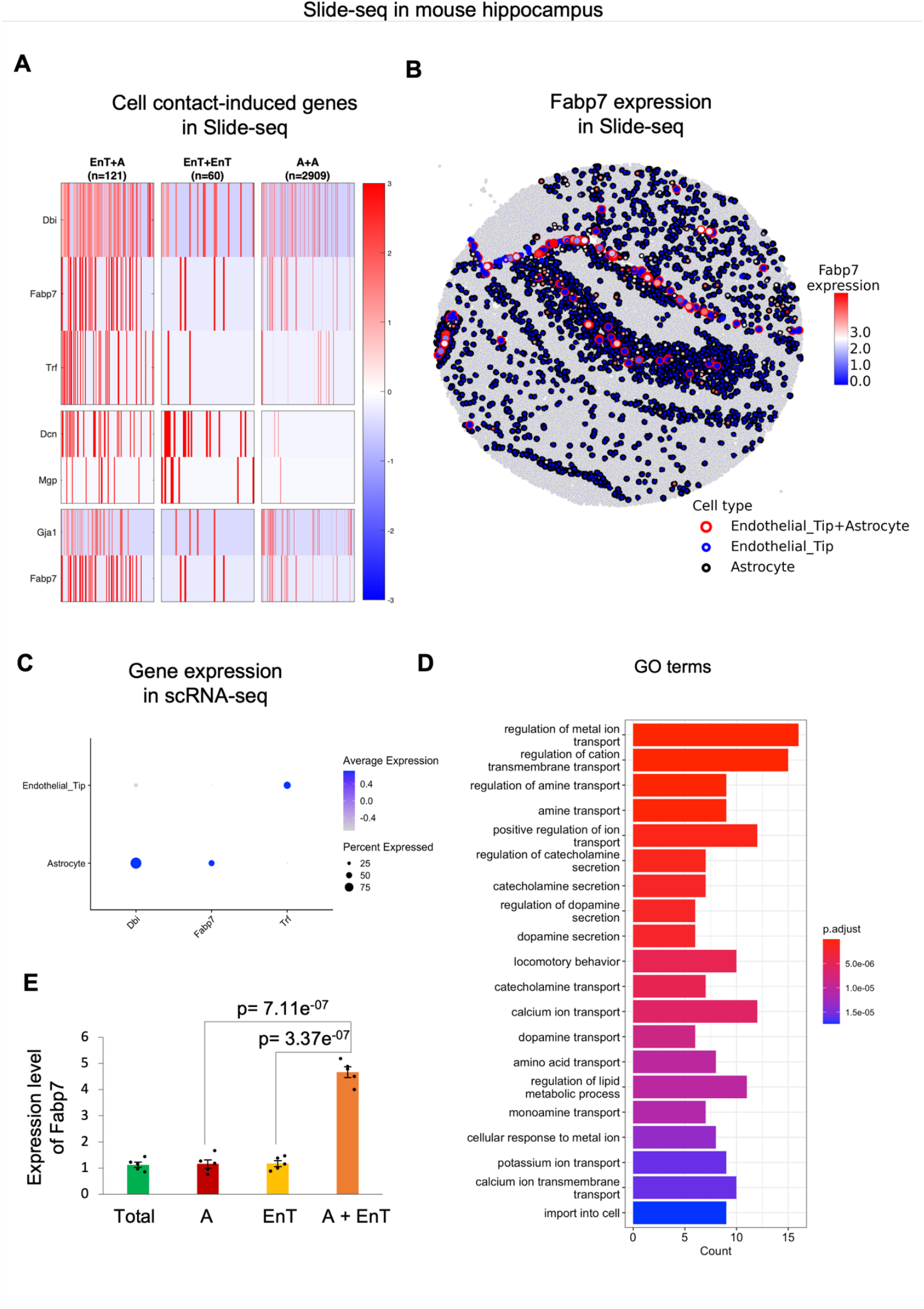
Neighbor dependent-genes identified by CellNeighborEX from the mouse hippocampus Slide-seq data. **A.** In the heatmaps, 3 genes including *Fabp7* are more highly expressed in the heterotypic spots of Endothelial tip and Astrocyte cells (EnT+A) than the respective homotypic spots (EnT, A). The heterotypic spots also express both EnT and Astrocyte markers. **B**. The spatial visualization shows the higher expression level of *Fabp7* in EnT+A. **C**. In the mouse hippocampus scRNA-seq data, *Fabp7* is mostly expressed in Astrocyte. **D**. In GO analysis, the GO terms are related to neuronal regulation. **E**. The analysis of qPCR-based mRNA expression of Astrocyte and EnT co-culture shows that *Fabp7* is up-regulated when compared to monocultures of Astrocyte (p-value = 7.11E-07) and EnT cells (p-value = 3.37E-07). The expression of *Fabp7* in the total mouse brain tissue represents the control (lane 1, green bar); we calculated the expression of *Fabp7* in monoculture and coculture relative to the expression of this gene in the total mouse brain tissue (N=5). The p-values shown in the bar plot were obtained by performing a two-tailed Student’s t-test.

To validate our findings, we designed an experiment to observe the expression of neighbor-dependent gene *Fabp7*. The expression of *Fabp7* was analyzed by qPCR-based mRNA expression analysis in monocultures of EnT and Astrocyte cells, and 48h co-culture of these two cell types in a 1:1 ratio (Fig. 3E). We observed that the expression of *Fabp7* was up-regulated in the co-culture of the two cell types, consistent with our prediction. Our results suggest that cell contact or proximal localization can induce the expression of specific genes.

### Neighbor-dependent genes are new potential genes involved in cell-cell interactions

Ligand-receptor co-expression has been used to study cell-cell interactions in ST data (Garcia-Alonso *et al*., 2021; Li *et al*., 2021; Pham *et al*., 2020; Shao *et al*., 2022). To see if the use of ligand-receptor pairs and the downstream genes mediated by them can recover our genes detected by CellNeighborEX, we ran NicheNet (Browaeys *et al*., 2020) on the seqFISH and Slide-seq datasets (Methods). We found 174, 1, 12, and 11 genes commonly detected between the two approaches, respectively (Fig. S6, Table S2). To see if the interactions are valid, we calculated minimum distances between the two interacting cell types using the spatial coordinates of the datasets. The estimated distances ranged from 60 to 1,600 μ*m* on average (Fig. S7). This suggests that the interactions identified by NicheNet may include many false predictions. NicheNet only finds frequently interacting cell types based on the averaged gene expression and it does not examine interactions between individual cells.

To demonstrate the usefulness of using the heterotypic spots in Slide-seq, we additionally ran NicheNet on the heterotypic spots. We set the heterotypic spots as a receiver as well as a sender (autocrine mode in NicheNet). We found that 2 genes (of 28 up-regulated genes from 9 heterotypic pairs) in the embryo, 15 genes (of 42 up-regulated genes from 10 heterotypic pairs) in the liver cancer, and additional 11 genes (of 155 up-regulated genes from 21 heterotypic pairs) in the hippocampus common to our neighbor-dependent genes (Table S3), suggesting that the use of heterotypic beads are useful in identifying genes related to cell communication.

### Neighbor-dependent genes demonstrates niche-specific expression

We further examined if neighbor-dependent gene expression suggests that cells express specific sets of genes depending on their niches. By running CellNeighborEX on the mouse embryo seqFISH data, we found that Gut tube cells highly express *Tbx1* when adjacent to Cranial mesoderm, *Pitx1* when adjacent to Neural crest, and *Foxf1* when adjacent to Splanchnic mesoderm in the mouse embryo (Fig. 4). To investigate niche-specific gene expression, we colored the boundary of Gut tube cells based on its neighboring cell types: red when proximal to Cranial mesoderm, green when to Neural crest, blue when to Splanchnic mesoderm, and orange when to another Gut tube. To easily distinguish gene expression change depending on neighboring cell types, we defined the neighboring cell type-specific genes using RGB color channels (Methods): *Tbx1* (red), *Pitx1* (green), and *Foxf1* (blue). Then, we represented the expression of these three genes using the combination of each color channel. Among the three genes, *Tbx1* (red inside the boundary) is dominantly expressed when Gut tube cells are next to Cranial mesoderm (Gut-tube/Cranial mesoderm, red boundary), *Pitx1* (green inside the boundary) is dominantly expressed when Gut tube cells are next to Neural crest (Gut-tube/Neural crest, green boundary), and *Foxf1* (blue inside the boundary) is dominantly expressed when Gut tube cells are next to Splanchnic mesoderm (Gut-tube/Splanchnic-mesoderm) (Fig. 4). These results indicate that Gut tube cells vary the expression levels of these genes depending on their neighboring cell types.

**Fig. 4.**
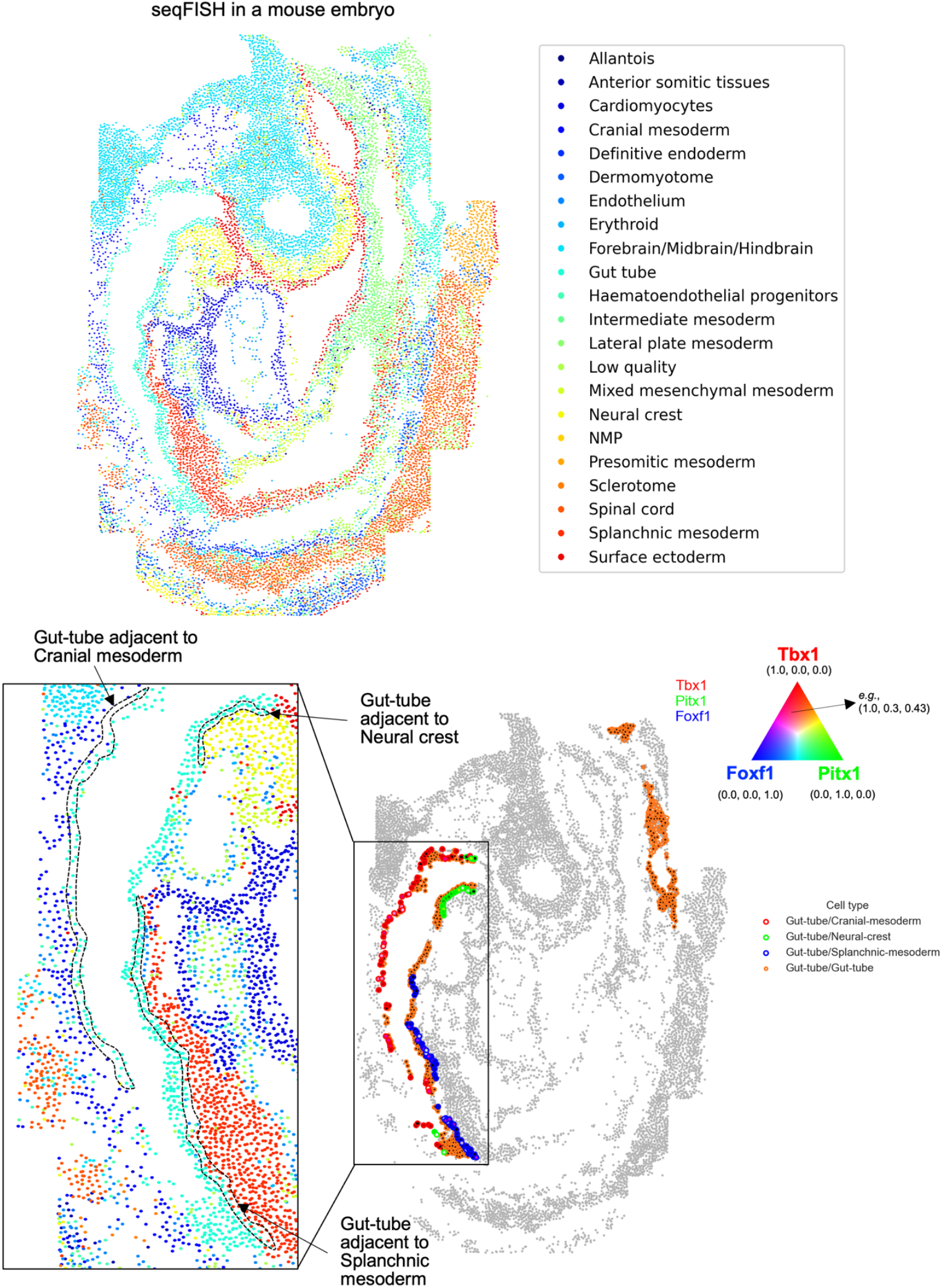
Niche-specific gene expression in seqFISH. The cells of the seqFISH data consist of 21 cell types except for Low quality (*i*.*e*., unidentified cells due to low quality). The spatial mapping with RGB channels simultaneously visualizes the expression of three neighbor-dependent genes. Gut tube cells express *Tbx1* (red) when adjacent to Cranial mesoderm, *Pitx1* (green) when adjacent to Neural crest, and *Foxf1* (blue) when adjacent to Splanchnic mesoderm.

In the Slide-seq data, we also observed similar niche-specific expression changes: in the embryo, *Hba-a1* (red) and *Cdkn1c* (green) are primarily expressed in Primitive erythroid lineage when contacting Definitive erythroid lineage (PEL+DEL) and Limb mesenchyme (PEL+LM), respectively (Fig. 5A). A regression study and investigation of scRNA-seq further confirmed that PEL cells mostly express these two genes in a neighboring cell type specific manner (Cao *et al*., 2019) (Fig. S4C, Fig. S4D, Fig. 5B). In the hippocampus, *Nnat* (red), *Gda* (green), and *Atp2b1* (blue) are mostly expressed in Entorhinal cells when contacting Choroid (Ento+Ch), Neuron.Slc17a6 (Ento+N), and Interneuron (Ento+In), respectively (Fig. 5C). We confirmed that the expression of the three genes was derived from Ento in the hippocampus scRNA-seq data (Saunders *et al*., 2018) (Fig. 5D). In the liver cancer, *Marco* (red), *Vti1a* (green), *F13a1* (blue) are largely expressed in Monocyte cells when contacting Hepatocyte I (Monocyte+Hepatocyte I), Hepatocyte II (Monocyte+Hepatocyte II), and Tumor III (Monocyte+Tumor III), respectively (Fig. 5E). From the regression models and snRNA-seq data in the liver cancer, we confirmed that Monocyte cells dominantly express the three genes from Monocyte (Fig. S4B, Fig. S4E, Fig. S4F, Fig. 5F). Our results indicate that cells communicate with the neighboring cells, actively responding to them.

**Fig. 5.**
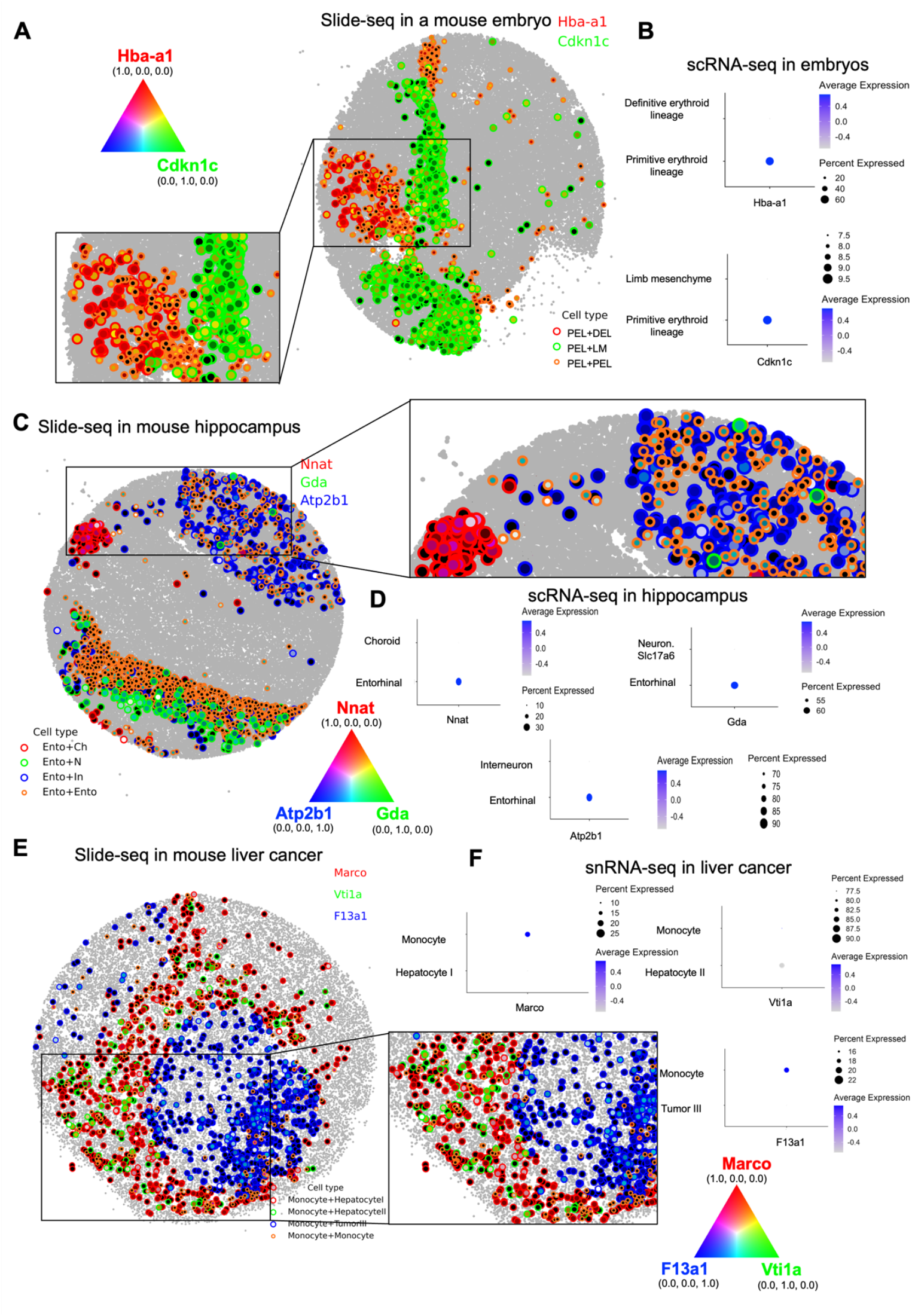
Niche-specific gene expression in Slide-seq. **A.** The spatial mapping with RGB channels displays the simultaneous expression of neighbor-dependent genes in the mouse embryo. Primitive erythroid lineage (PEL) cells dominantly express *Hba-a1* (red) when contacting to Definitive erythroid lineage (DEL), and *Cdkn1c* (green) when contacting Limb mesenchyme (LM). **B**. In the mouse embryo scRNA-seq data, *Hba-a1* and *Cdkn1c* are expressed from PEL. **C**. In the Spatial mapping for the mouse hippocampus, Entorhinal (Ento) cells express *Nnat* (red) when contacting Choroid (Ch), *Gda* (green) when contacting Neuron.Slc17a6 (N), and *Atp2b1* (blue) when contacting Interneuron (In). **D**. In the mouse hippocampus scRNA-seq data, *Nnat, Gda*, and *Atp2b1* are mostly expressed from Ento. **E**. In the spatial mapping for the mouse liver cancer, Monocyte cells dominantly express *Marco* (red) when contacting Hepatocyte I, *Vti1a* (green) when contacting Hepatocyte II, and *F13a1* (blue) when contacting Tumor III. **F**. In the mouse liver cancer snRNA-seq data, *Marco, Vti1a*, and *F13a1* are expressed from Monocyte cells.

### Niche specific-gene expression accounts for cellular heterogeneity

Cellular heterogeneity is caused by a number of reasons in different contexts. scRNA-seq has been useful in describing cell heterogeneity but could not explain the cause of heterogeneity. We tested if the niche specific genes obtained from ST study can account for cellular heterogeneity in scRNA-seq. We selected a cell type showing neighbor-dependent gene expression. Fig. S8A shows an example of neighboring cell type-dependent gene expression in the hippocampus Slide-seq V2 data. Endothelial tip cells dominantly express *Igfbp7* (red) when proximal to Choroid (EnT+Ch), *Trf* (green) when proximal to Astrocyte (EnT+A), and *Plp1* (blue) when proximal to Interneuron (EnT+In). In the hippocampus scRNA-seq data, it was confirmed that the three genes were mostly expressed from EnT (Fig. S8B).

To investigate the heterogeneity of EnT cells, we separately selected EnT cells in the scRNA-seq data. We performed clustering analysis using the expression values of the three genes and identified 4 clusters (Fig. S8C). The UMAP plot shows the expression of *Igfbp7, Trf*, and *Plp1* in the corresponding clusters. We further labeled them based on the neighboring cell type information (Fig. S8D). We also found an additional example for the heterogeneity of Interneuron cells in the hippocampus Slide-seq V2 data (Fig. S9). These findings suggest a possibility that cellular heterogeneity might be stemmed from neighboring cell type-dependent gene expression.

## Discussion

Cell communication is a fundamental process related to various functions such as cell growth, development, and diseases (Yang *et al*., 2021). Cell communication coordinates the functions of multicellular organisms (Radhakrishnan *et al*, 2010). Even with its importance, a systematic study of cell communication was not easy without a well-curated experimental setup until scRNA-seq is available. To study cell-cell interactions from scRNA-seq, co-expression of ligand-receptor pairs has been used (Browaeys *et al*., 2020; Efremova *et al*., 2020). However, it was not possible to study other types of cell communication such as direct contact due to the loss of spatial information in scRNA-seq.

RNA sequencing of physically interacting multi-cells or PIC-seq has provided the transcriptomic landscapes of cells contacting each other (Boisset *et al*, 2018; Giladi *et al*, 2020; Kim *et al*., 2023). In our previous study using PIC-seq from mouse embryos, we found that direct cell contact can induce the expression of specific gene sets depending on the neighboring cell types (Kim *et al*., 2023). This unbiased approach systematically studied cell contact-dependent expression during mouse development. However, PIC-seq can be biased to cell interactions more strongly bound to each other.

ST technologies have enabled spatial mapping of gene expression and have provided opportunities to look into cellular microenvironments. Accordingly, a number of computational methods that investigate cell-cell interactions in the spatial domain have been developed (Arnol *et al*., 2019; Cable *et al*., 2022; Fischer *et al*., 2023; Garcia-Alonso *et al*., 2021; Li *et al*., 2021; Li & Yang, 2022; Pham *et al*., 2020; Shao *et al*., 2022; Tanevski *et al*., 2022). However, cell communication through direct cell contact has not been thoroughly explored yet. Still, most methods focus on finding frequently interacting cell type pairs based on ligand-receptor c o-expression (Garcia-Alonso *et al*., 2021; Li *et al*., 2021; Pham *et al*., 2020; Shao *et al*., 2022). Here, we presented CellNeighborEX to analyze the influence of direct cell contact on the transcriptome of cells from ST data. CellNeighborEX was designed to work for both image- and NGS-based ST data by defining immediate neighbors differently (Fig. 1). For image-based ST data where exact cell locations are available, CellNeighborEX can use various algorithms including Delaunay triangulation, radial distance, and k-nearest neighbors (KNN) to find immediate neighbors. For NGS-based ST data where exact cell locations are not available, we used the heterotypic beads.

The use of heterotypic beads depends on the resolution of ST data. We have shown that CellNeighborEX successfully identified neighboring cell type-dependent genes from Slide-seq with 10 um resolution. However, it is not suitable to detect neighbor-dependent genes from ST data with low resolution such as Visium (Ståhl *et al*., 2016) as there could be more than 2 cell types. For the higher resolution ST data such as Seq-Scope (Cho *et al*, 2021), it is possible to use heterotypic beads by rescaling them to 10 um resolution.

CellNeighborEX identified neighbor-dependent genes from various ST datasets including mouse embryos (Fig. 2 and S3), liver cancer (Fig. S5), and mouse hippocampus (Fig. 3). Our results indicate that neighbor-dependent genes are found in most of the cell types and tissues, further expanding our observation in the developing mouse embryos (Kim *et al*., 2023).

Interestingly, we found that genes influenced by neighbors were associated with important cell functions. For instance, CellNeighborEX found neighbor-dependent genes are associated with embryonic development from both seqFISH (Fig. 2D) and Slide-Seq V2(Fig. S3D) data. Our results may suggest that cell contact triggers genes important for further development. CellNeighborEX also provided information about cell types and genes influenced by TME. For instance, CellNeighborEX found that *F13a1* is highly expressed when Monocyte cells contact Tumor cells. *F13a1* has been known to be associated with various cancers including liver cancer (Dong *et al*., 2022; Peltier *et al*., 2018; Porrello *et al*., 2018). In the GO analysis, we found that the neighbor-dependent genes are further associated with cancer metastases (Fig. S5D). Our results show that CellNeighborEX is a useful tool to study the influence of TME in an unbiased way. Besides, the experiment using a co-culture system clearly demonstrated the higher expression of neighbor-dependent gene (Fig. 3E). It is also of note that we saw a little overlap with the results obtained from ligand-receptor pairs (Fig. S6). These findings indicate that studying direct cell contact is important to understand cell-cell interactions more thoroughly.

We used heterotypic beads as evidence for cell contact in NGS-based ST data. The neighbor-dependent genes that we identified from Slide-seq data were more highly expressed than artificially generated null models (Fig. S1). From the regression model using the heterotypic beads, we predicted cell types expressing the neighbor-dependent genes (Fig. S4). Our strategy suggests new ways to utilize heterotypic beads in the high-resolution ST data such as Slide-Seq.

We observed that cells express specific sets of genes depending on their neighbors (Fig. 4, Fig. 5). This niche-specific gene expression partly explains the cause of cellular heterogeneity shown in scRNA-seq data (Fig. S8, Fig. S9). Also, this can suggest that we can re-annotate cells based on their neighboring cell types.

ST information has been used to understand cell communication. NCEM (Fischer *et al*., 2023) is a graph neural network model to investigate the influence of neighboring cells on gene expression. NCEM uses neighbors in an intermediate range while we focus on immediate neighbors, thereby having the lower complexity of neighboring cell types. That allows studying the influence of direct contact between two different cell types. From it, the gene expression changes can be explicitly validated in the spatial domain, suggesting niche-specific expression.

To sum up, CellNeighborEX is a new approach to explore transcriptomic changes caused by direct cell contact from ST data. Studying cell contact-dependent gene expression provides opportunities to understand cell-cell interactions between two adjacent cells from a new perspective. It enabled the identification of new genes potentially involved in intercellular communication beyond previous approaches that use ligand-receptor pairs. It also demonstrated gene expression varies depending on neighboring cell types, explaining cellular heterogeneity.

## Materials and Methods

### Data pre-processing

For seqFISH data in a mouse embryo (Lohoff *et al*., 2022), we used the gene expression data and annotated cell types pre-processed by Squidpy (Palla *et al*, 2022). For Slide-seq data in a mouse embryo (Stickels *et al*., 2021), hippocampus (Stickels *et al*., 2021), and liver cancer (Zhao *et al*., 2022), we obtained them from Puck_190926_03, Puck_200115_08, and mouse_liver_met_2_rna_201002_04, respectively. For the embryo and liver cancer, samples that have unique feature counts less than 200 were filtered out. For the hippocampus, all samples were used without the filtering because the samples with unique feature counts less than 200 take up a considerable percentage of the total samples (*i*.*e*., about 40%). The values of the count matrix per dataset were log-normalized and then the top 2000 variable genes as well as cell type markers (Table S4) were selected.

For liver cancer snRNA-seq data, we pre-processed paired dataset (Zhao *et al*., 2022) given with the Slide-seq data in the mouse liver cancer. The genes expressed in less than 3 nuclei were filtered out. The samples that have unique feature counts less than 200 and mitochondrial RNA larger than one percent were filtered out. Additionally, approximately 10 percent of doublets detected by DoubletFinder (McGinnis *et al*, 2019) were removed. After the clustering analysis, we annotated the cell clusters using the information of cell type markers accompanied with the snRNA-seq dataset. The analyses mentioned above were all performed with Seurat 3.2.2 (Satija *et al*., 2015).

### Cell type inference of Slide-seq spots

We used RCTD (Cable *et al*., 2021) to identify the cell types of the spots in Slide-seq. To run RCTD, we trained RCTD with scRNA-seq or snRNA-seq datasets with annotated cell types. For the embryo, we used a scRNA-seq dataset (Cao *et al*., 2019) at E12.5 equivalent to the developmental stage of Slide-seq embryo. It consists of 26,183 genes and 270,197 cells assigned into 37 cell types (Fig. S10A). For the hippocampus, we obtained a scRNA-seq dataset (Saunders *et al*., 2018) from DropViz. It is composed of 27,953 genes and 113,507 cells assigned into 17 cell types (Fig. S10B). For the liver cancer data, we used the paired snRNA-seq dataset (Zhao *et al*., 2022) given with the Slide-seq data in mouse liver cancer. The pre-processed snRNA-seq dataset consists of 24,098 genes and 11,683 nuclei assigned into 14 cell types (Fig. S10C). Training RCTD, we predicted the cell types of spots in Slide-seq. The simulation was performed under doublet mode (Cable *et al*., 2021) that constraints each spot to contain up to two cell types, which is recommended for data with fine resolution such as Slide-seq. From RCTD, we identified the cell type of the spots and further estimated the cell type proportions for each spot.

The results on the inference of cell types were additionally validated from correlation analysis. We examined the correlation between the ture heterotypic spots annotated by RCTD and artificial heterotypic spots. Specifically, we generated artificial heterotypic spots by combining two homotypic spots based on the cell type proportions of heterotypic spots obtained from RCTD (Fig. S1).

With repeated random sampling, we created 100 artificial heterotypic spots for each heterotypic spot. Next, we calculated Pearson’s correlation coefficients based on the gene expression values between the true heterotypic spots and many combinatorial types of artificial heterotypic spots (Fig. S2). If the cell type of true heterotypic spots is consistent with that of artificial ones with the largest Pearson’s coefficient, we regarded the cell type of the true heterotypic spots as validated. On top of the correlation analysis, the inferred cell types were validated by cell type markers (Table S4) accompanied with the scRNA-seq (Cao *et al*., 2019; Saunders *et al*., 2018) and snRNA-seq (Zhao *et al*., 2022) datasets, respectively.

### Neighbor-dependent gene expression analysis

We studied neighbor-dependent gene expression by comparing heterotypic groups (heterotypic neighbors for seqFISH, heterotypic spots for Slide-seq) with homotypic groups (homotypic neighbors for seqFISH, homotypic spots for Slide-seq). We identified genes up- and down-regulated by direct cell contact. We carried out rigorous statistical analysis between the two groups. We determined whether to use parametric or non-parametric two-sided tests depending on the sample size of groups. When both samples were larger than sample size 30, we chose parametric tests under the normality assumption. To be specific, we conducted the Student’s t-test for equal variances and the Welch’s t-test for unequal variances, where a two-sample F-test was used to test whether the variances are equal or not.

For the Student’s t-test, the *t* statistic is calculated as follows:

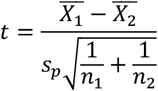

where 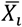 is the mean of expression values in group *i*(*i*=1,2). In the equations, group 1 and group 2 represent heterotypic group and homotypic group, respectively. *n*_*i*_ is the sample size of group *i. s*_*p*_ is the pooled standard deviation of the two groups:

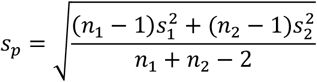

where *s*_*i*_ is the standard deviation of group *i*.

For the Welch’s t test, the *t* statistic is computed as follows:

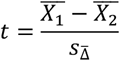

where 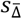is the

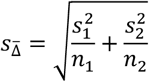

Meanwhile, when the sample size of at least one sample was smaller than 30, we performed the Mann-Whitney U test as a non-parametric test. The *U* statistic is calculated as follows:

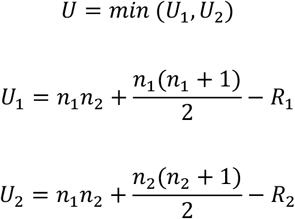

where *R*_1_is the sum of the ranks for the heterotypic group, and *R*_2_is the sum of the ranks for the homotypic group.

In the DE analysis, the log2(fold-change) >0.4 and p-value < 0.01 were used as criteria for differential expression. For the seqFISH data, FDR < 0.05 was added as an additional criterion.

### Verification of cell-cell interactions in the heterotypic spots of Slide-seq data

We developed a null model to verify that individual heterotypic spots represent two different cell types interacting with each other. Our null model refers to artificial heterotypic spots (Fig. S1). In contrast with the true heterotypic spots, the artificial heterotypic spots indicate two different cell types just combined without cell-cell interactions. We compared the true heterotypic spots with the artificial heterotypic spots to confirm the statistical significance of the neighbor-dependent genes. The significant neighbor-dependent genes mean that their expression resulted from interacting two cell types. The same statistical tests as the neighbor-dependent gene expression analysis were applied. The log2(fold-change) >0.4, p-value < 0.01, and FDR < 0.01 were used as criteria for differential expression.

### Finding cell types expressing neighbor-dependent genes in the heterotypic spots of Slide-seq

Heterotypic spots in Slide-seq represent interacting two different cell types. For the genes up-regulated by cell contact, it is challenging to find which cell type the expression of the neighbor-dependent genes come from between the two cell types. We created linear regression models to find the origin of the expression. For instance, we suppose that *g* is a neighbor-dependent gene found from the heterotypic spots of *A + B*. For *n* data pairs { (*x*_*i*_, *y*_*i*_), *i*=1,2, …, *n}*, the regression model is as follows:

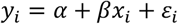

where *n* is the number of *A + B*heterotypic spots expressing *g. x*_*i*_ is the proportion of *A*, 1 − *x*_*i*_ is the proportion of *B*, and *y*_*i*_ is the expression value of *g* in heterotypic spot *A+B*_*i*_. We use the ordinary least squares method to find the intercept 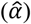 and slope (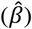) illustrating the best fit line to the *n* data pairs. Estimated 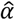 and 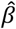 are computed as follows:

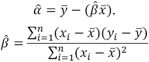

where 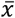 and 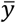 are the mean of the *x*_*i*_ and *y*_*i*_, respectively. If slope 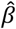 is positive, it indicates that the expression value of *g* increases as the proportion of cell type *A* grows. From this, we can infer that the expression of *g* comes from cell type *A*. If 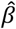 is negative, the expression of *g* decreases as the proportion of cell type *A* becomes larger. It means that the expression of *g* increases as the proportion of cell type *B* grows. That is, the expression of *g* comes from cell type *B*.

We applied the regression models to the neighbor-dependent genes obtained from Slide-seq when the number of the heterotypic spots is large enough. We additionally used scRNA-seq or snRNA-seq data (Fig. S11) and found that the cell types inferred from the single cell or single nucleus data are considerably consistent with the cell types predicted from the statistically significant regression models (*i*.*e*., p-value < 0.05) (Fig. S4, Table S5).

### Spatial visualization with RGB color channels

We used RGB coordinates (r, g, b) composed of values between 0 and 1 to observe how gene expression varies depending on neighboring cell types. We first selected one cell type and then collected multiple heterotypic groups where the selected cell type is included. For example, if the cell type of interest is *A*, heterotypic groups *A+B, A+C*, and *A+D* are collected. If *gene* γ, *gene* δ, and gene *ρ* are neighbor-dependent genes found from the heterotypic spots of *A+B, A+C*, and *A+D* respectively, red, green, and blue channels are assigned to the expression values of the three neighbor-dependent genes: *gene* γ is red (R), *gene δ* is green (G), and gene *ρ* is blue (B). To normalize the expression values between 0 and 1, we divided them by the maximum expression value in each heterotypic group. In case that there are two genes, the value of the blue channel is fixed as zero. The expression of the three neighbor-dependent genes is simultaneously visualized by the RGB channels.

### Identification of ligands, receptors, and downstream targets

We used NicheNet (Browaeys *et al*., 2020) to compare our neighbor-dependent genes with already known ligands, receptors, and their target genes. We ran NicheNet on the seqFISH and Slide-seq data to see if NicheNet can detect our neighbor-dependent genes. For the seqFISH, we investigated cells corresponding to the heterotypic neighbors (22 types of heterotypic neighbors in the embryo) where our neighbor-dependent genes were identified. We set the centered cell type as a sender and the neighboring cell type as a receiver, vice versa. In the case of Slide-seq, we used homotypic spots corresponding to the heterotypic pairs (9 heterotypic pairs in the embryo, 10 heterotypic pairs in the liver cancer, and 21 heterotypic pairs in the hippocampus). We set the respective homotypic spots as a sender and a receiver by turns.

To validate if the heterotypic spots in Slide-seq are useful to study cell-cell interactions, we examined the heterotypic spots. We set the heterotypic spots as a receiver as well as a sender (autocrine mode in NicheNet). Highly expressed ligand-receptor-target genes detected by NicheNet mean being expressed in at least 10% of cells in one cluster.

### Experimental validation for cell contact-specific gene expression

We isolated total RNA from the individual cultures and cocultures of Endothelial tip and Astrocyte cells. We used the PureLink RNA Mini kit, as per the manufacturer’s instruction, and eluted total RNA in 50 μL RNase/DNase-free H2O. Then, we reverse-transcribed to cDNA 10 ng of total RNA using Superscript Vilo cDNA synthesis kit. Finally, we carried out real-time PCR (qPCR) in QuantStudioTM 5 (Applied Biosystems) using PowerUp SYBR Green master mix (Thermo Fisher Scientific) and the following reaction conditions. The initial denaturation step was performed at 95 °C for 2 minutes, followed by 40 cycles of 95 °C for 15 seconds and 60 °C for 60 seconds. We used the comparative CT method (ΔΔCt) to quantify relative gene expression, normalizing the expression of our target genes with the housekeeping gene Gapdh. All samples were run by using the following primers: Gapdh: 5′-CATCACTGCCACCCAGAAGACTG-3′(F) and 5′-ATGCCAGTGAGCTTCCCGTTCAG-3′ (R); and *Fabp7*: 5′-TGGGAAACGTGACCAAACCA-3′ (F) and 5′-AGCTTGTCTCCATCCAACCG-3′ (R).

### Analyzed publicly available datasets

- seqFISH data in a mouse embryo: Squidpy (https://squidpy.readthedocs.io/en/stable/auto_tutorials/tutorial_seqfish.html).
- Slide-seq V2 data in a mouse embryo and hippocampus: Single Cell Portal (https://singlecell.broadinstitute.org/single_cell/study/SCP815/sensitive-spatial-genome-wide-expression-profiling-at-cellular-resolution#study-summary).
- Slide-seq and snRNA-seq data in mouse liver cancer: Single Cell Portal (https://singlecell.broadinstitute.org/single_cell/study/SCP1278/spatial-genomics-enables-multi-modal-study-of-clonal-heterogeneity-in-tissues).
- scRNA-seq data in a mouse embryo: Gene Expression Omnibus GSE119945 (https://www.ncbi.nlm.nih.gov/geo/query/acc.cgi?acc=GSE119945).
- scRNA-seq data in mouse hippocampus: DropViz (http://dropviz.org/).

## Code availability

Codes for our neighbor-dependent gene expression analysis are available at https://github.com/hkim240/CellContact.

## Acknowledgements

We appreciate Dr. Patrick Martin for critical reading of the manuscript. We appreciate the institutional support from Cedars-Sinai Medical Center.

