## supplementary_materials for "CellNeighborEX: Deciphering Neighbor-Dependent Gene Expression from Spatial Transcriptomics Data"

**This PDF file includes:**

Supplementary Figures 1 to 11

Supplementary Tables 1 to 5

#### Supplementary Figures

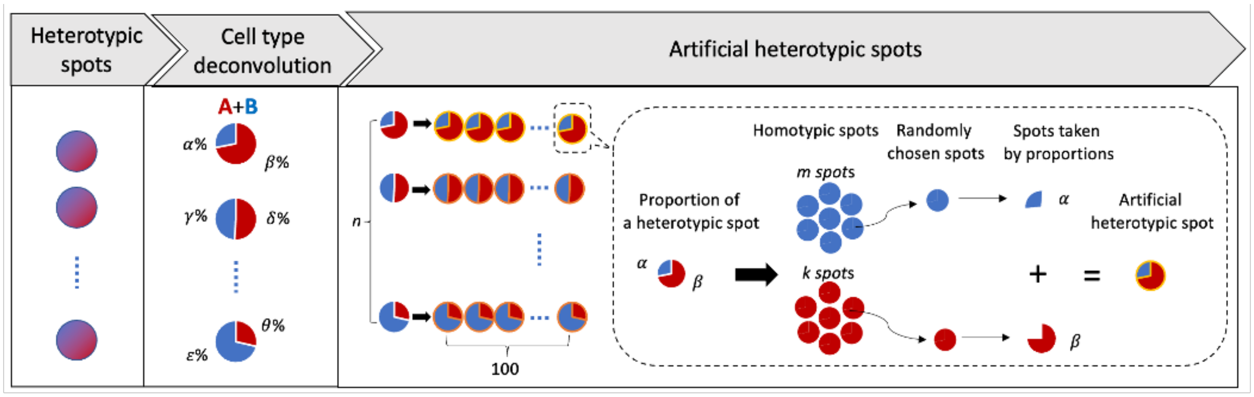

**Fig. S1. Generation of artificial heterotypic spots for statistical tests.** Artificial heterotypic spots are produced by combining two different cell types of homotypic spots. For the two cell types, a homotypic spot is randomly chosen respectively and then their transcriptomes are mixed according to the cell type proportions of the heterotypic spots. From repeated random sampling, 100 artificial heterotypic spots are created for each heterotypic spot.

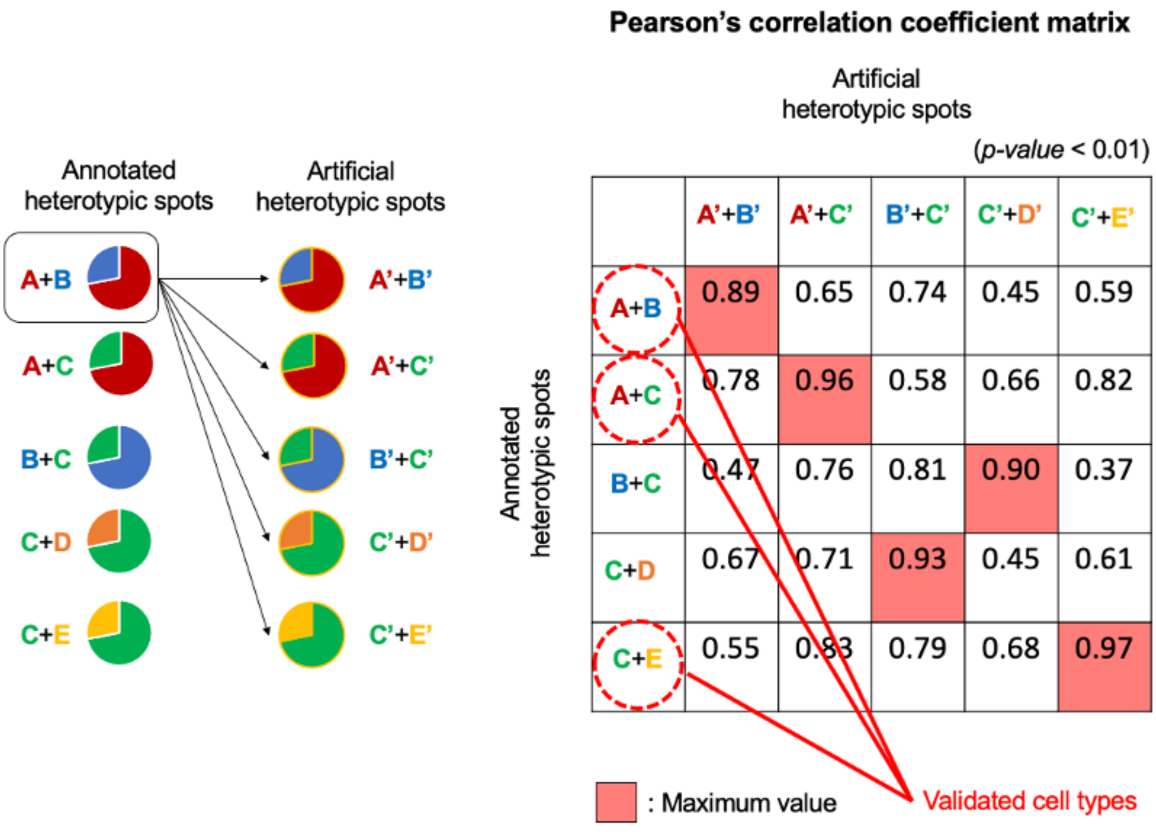

**Fig. S2. Validation of cell types of heterotypic spots.** Correlation analysis is performed to validate if the cell types of the heterotypic spots annotated by RCTD are correct. The Pearson's correlation coefficients are calculated based on the gene expression values between true heterotypic spots and many combinatorial types of artificial heterotypic spots. In the case that the cell type of true heterotypic spots matches that of artificial heterotypic spots with the largest Pearson's coefficient, the cell type annotation is regarded as validated.

Slide-seq in a mouse embryo

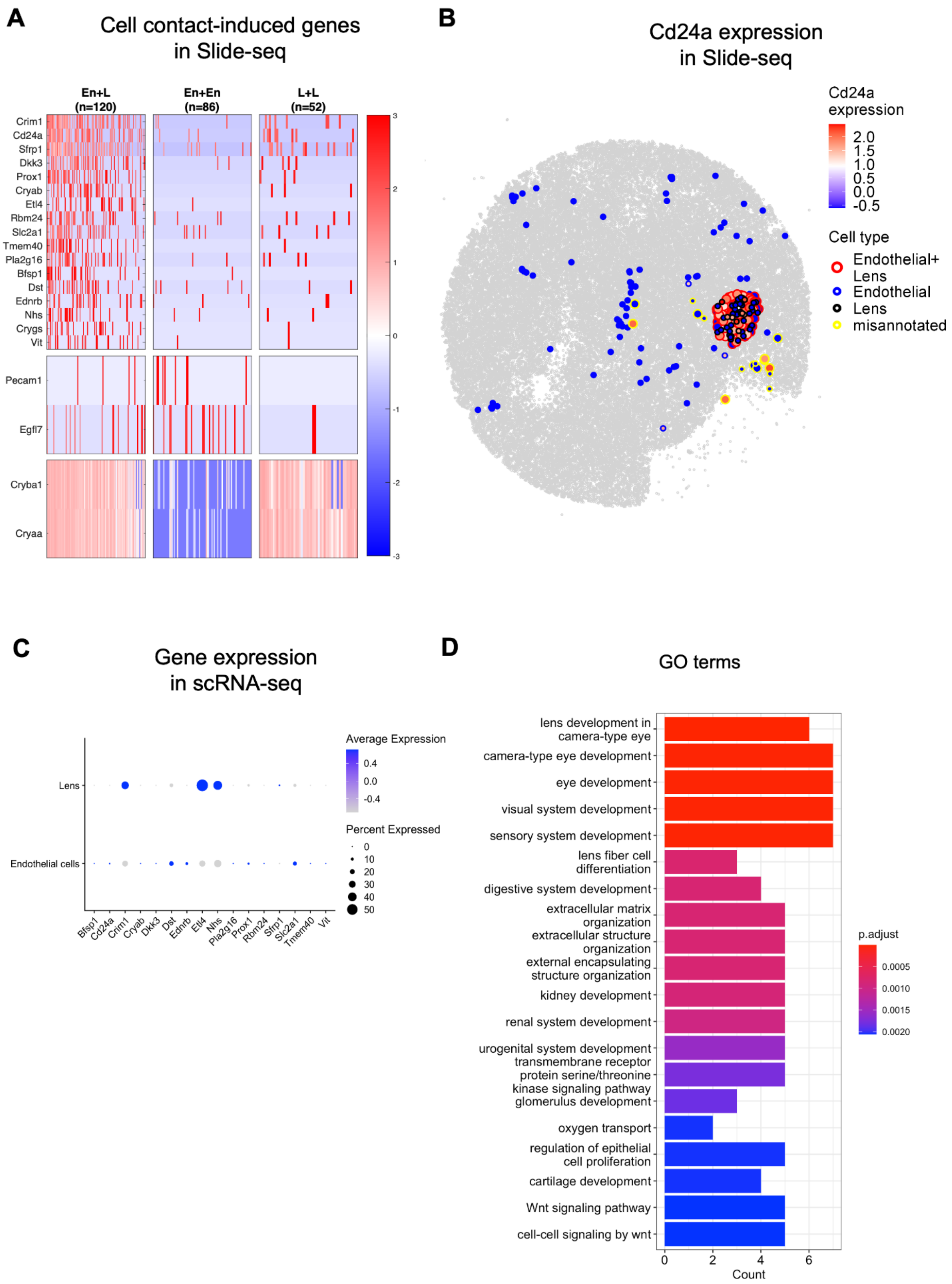

**Fig. S3. Transcriptomic change due to direct cell contact in the mouse embryo Slide-seq data.** **A.** Neighbor-dependent gene expression in the embryo. In the heatmaps, 17 genes including *Cd24a* are highly expressed in the heterotypic spots of Endothelial and Lens cells (En+L) than the respective homotypic spots (En, L). The heterotypic spots also express both En and L markers. **B.** The spatial visualization shows the higher expression level of *Cd24a* in En+L. **C.** Expression of neighbor-dependent genes in the mouse embryo scRNA-seq data. It confirms that *Cd24a* is expressed in En. **D.** GO terms for the genes influenced by neighbors are associated with embryonic development.

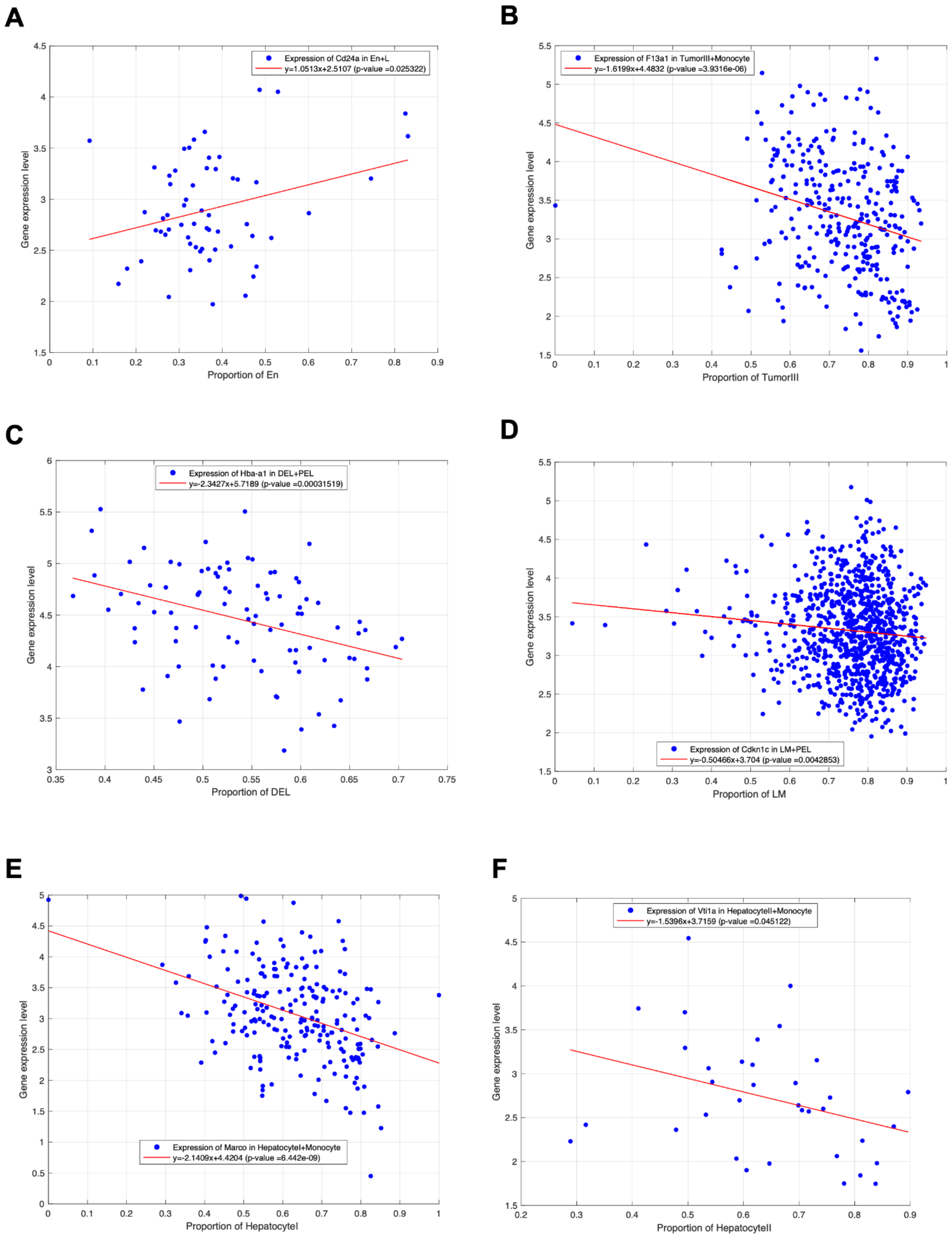
A
Expression of Cd24a in En+L
y=1.0513x+2.5107 (p-value =0.025322)
Gene expression level
Proportion of En
B
Expression of F13a1 in TumorIII+Monocyte
y=-1.6199x+4.4832 (p-value =3.9316e-06)
Gene expression level
Proportion of TumorIII
C
Expression of Hba-a1 in DEL+PEL
y=-2.3427x+5.7189 (p-value =0.00031519)
Gene expression level
Proportion of DEL
D
Expression of Cdkn1c in LM+PEL
y=-0.50466x+3.704 (p-value =0.0042853)
Gene expression level
Proportion of LM
E
Expression of Marco in HepatocyteI+Monocyte
y=-2.1409x+4.4204 (p-value =6.442e-09)
Gene expression level
Proportion of HepatocyteI
F
Expression of Vti1a in HepatocyteII+Monocyte
y=-1.5396x+3.7159 (p-value =0.045122)
Gene expression level
Proportion of HepatocyteII

**Fig. S4. Regression models to determine the origin of the expression of neighbor-dependent genes. A.** The expression of *Cd24a* in the heterotypic spots of Endothelial and Lens cells (En+L). X-axis indicates the ratio of Endothelial cells against Lens cells in the heterotypic spots. The positive relationships indicate that *Cd24a* is expressed in Endothelial cells **B.** The expression of Hba-a1 in the heterotypic spots of Definitive erythroid lineage and Primitive erythroid lineage cells (DEL+PEL). It suggests that *Hba-a1* is expressed in PEL. **C.** The expression levels of *Cdkn1c* in the heterotypic spots of Limb mesenchyme and Primitive erythroid lineage cells (LM+PEL). It suggests that the expression of *Cdkn1c* is expressed in PEL. **D.** The expression levels of *Marco* in the heterotypic spots of Hepatocyte I and Monocyte cells (Hepatocyte I+Monocyte). It suggests that the expression of *Marco* is expressed in Monocyte cells. **E.** The expression levels of *Vti1a* in the heterotypic spots of Hepatocyte II and Monocyte cells (Hepatocyte II+Monocyte). It suggests that the expression of *Vti1a* is expressed in Monocyte cells. **F.** The expression levels of *F13a1* expression in the heterotypic spots of Tumor III and Monocyte cells (Tumor III+Monocyte). It suggests that the expression of *F13a1* is expressed in Monocyte cells.

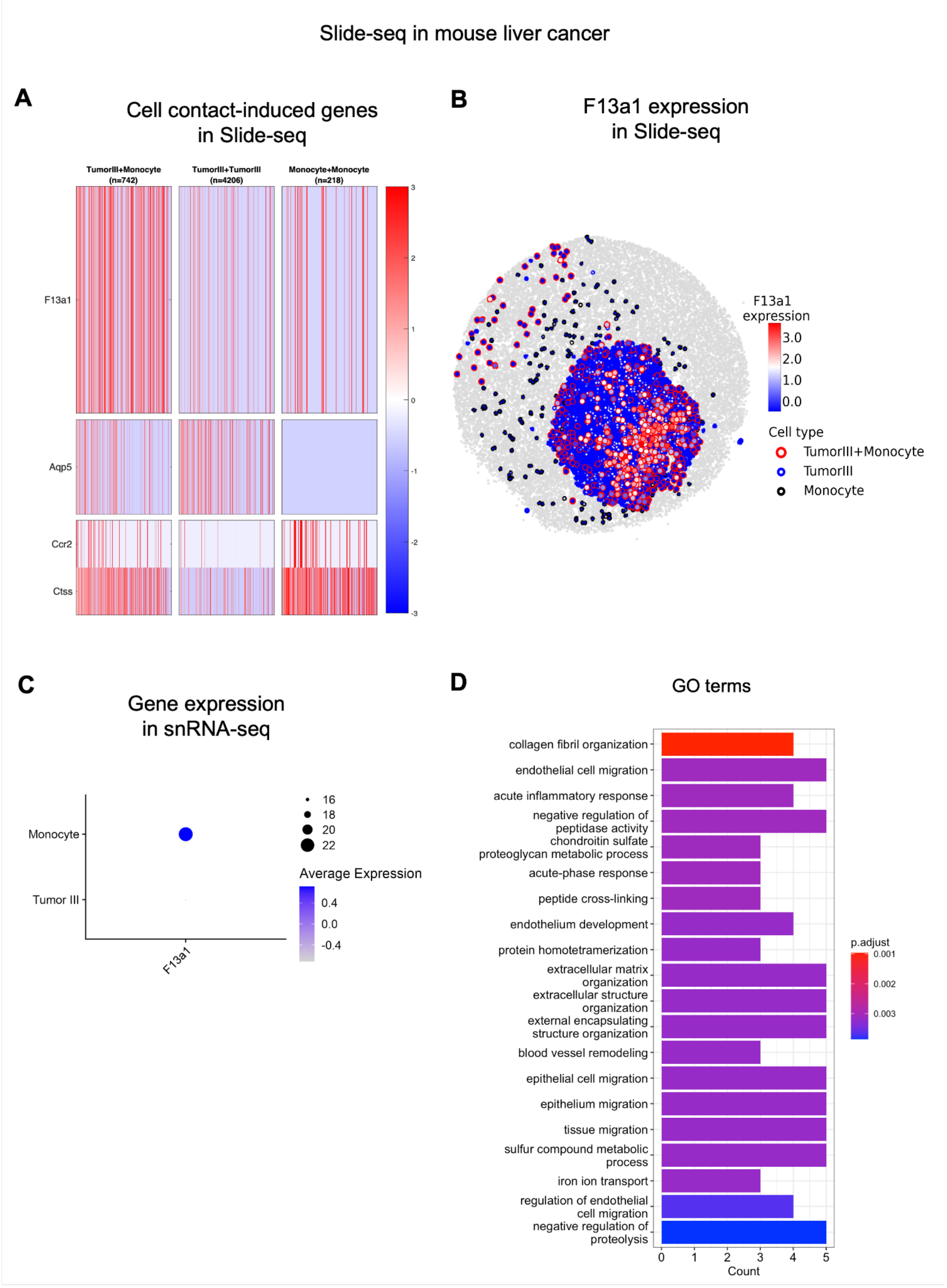
Slide-seq in mouse liver cancer
A
Cell contact-induced genes
in Slide-seq
TumorIII+Monocyte
(n=742)
TumorIII+TumorIII
(n=4206)
Monocyte+Monocyte
(n=218)
F13a1
Aqp5
Ccr2
Ctss
3
2
1
0
-1
-2
-3
B
F13a1 expression
in Slide-seq
F13a1
expression
3.0
2.0
1.0
0.0
Cell type
TumorIII+Monocyte
TumorIII
Monocyte
C
Gene expression
in snRNA-seq
Monocyte
Tumor III
F13a1
16
18
20
22
Average Expression
0.4
0.0
-0.4
D
GO terms
collagen fibril organization
endothelial cell migration
acute inflammatory response
negative regulation of peptidase activity
chondroitin sulfate proteoglycan metabolic process
acute-phase response
peptide cross-linking
endothelium development
protein homotetramerization
extracellular matrix organization
extracellular structure organization
external encapsulating structure organization
blood vessel remodeling
epithelial cell migration
epithelium migration
tissue migration
sulfur compound metabolic process
iron ion transport
regulation of endothelial cell migration
negative regulation of proteolysis
0
1
2
3
4
5
Count
p.adjust
0.001
0.002
0.003

**Fig. S5. CellNeighborEx identified genes influenced by TME. A.** Neighbor-dependent gene expression in liver cancer. In the heatmaps, *F13a1* is more highly expressed in the heterotypic spots of Tumor III and Monocyte cells (Tumor III+Monocyte) than the respective homotypic spots (Tumor III, Monocyte). The heterotypic spots also express both Tumor III and Monocyte markers. **B.** The spatial visualization shows the higher expression level of *F13a1* in Tumor III+Monocyte. **C.** Expression of neighbor-dependent genes in the mouse liver cancer snRNA-seq data. It confirms that *F13a1* is mostly expressed in Monocyte cells. **D.** GO analysis in the mouse liver cancer data. The terms are associated with tumor metastasis.

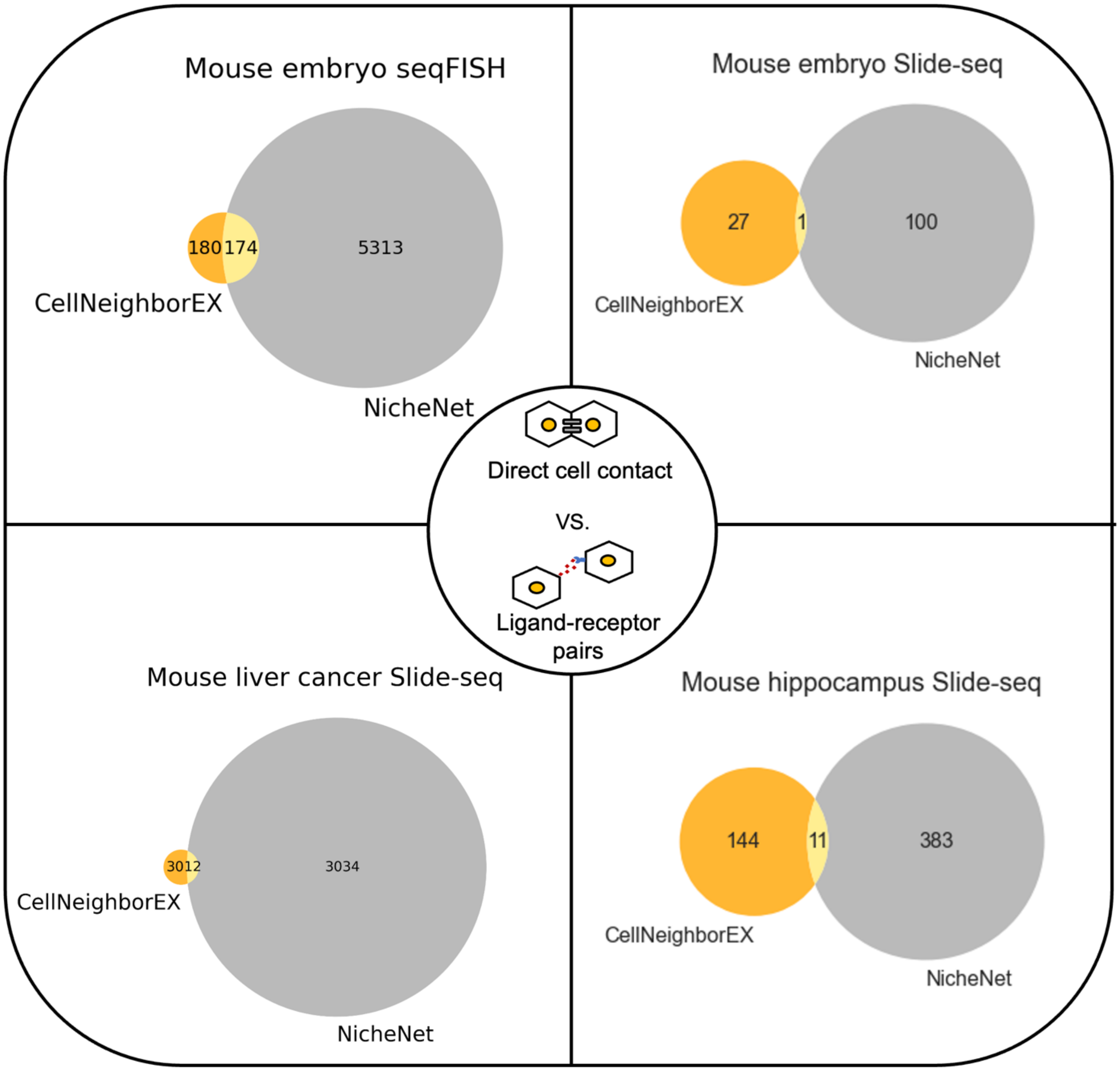

**Fig. S6. Comparison of neighbor-dependent genes to ligand-receptor pairs & downstream targets.** In Venn diagrams, "CellNeighborEX" represents genes up-regulated by cell contact while "NicheNet" indicates ligand-receptor pairs and their downstream target genes.

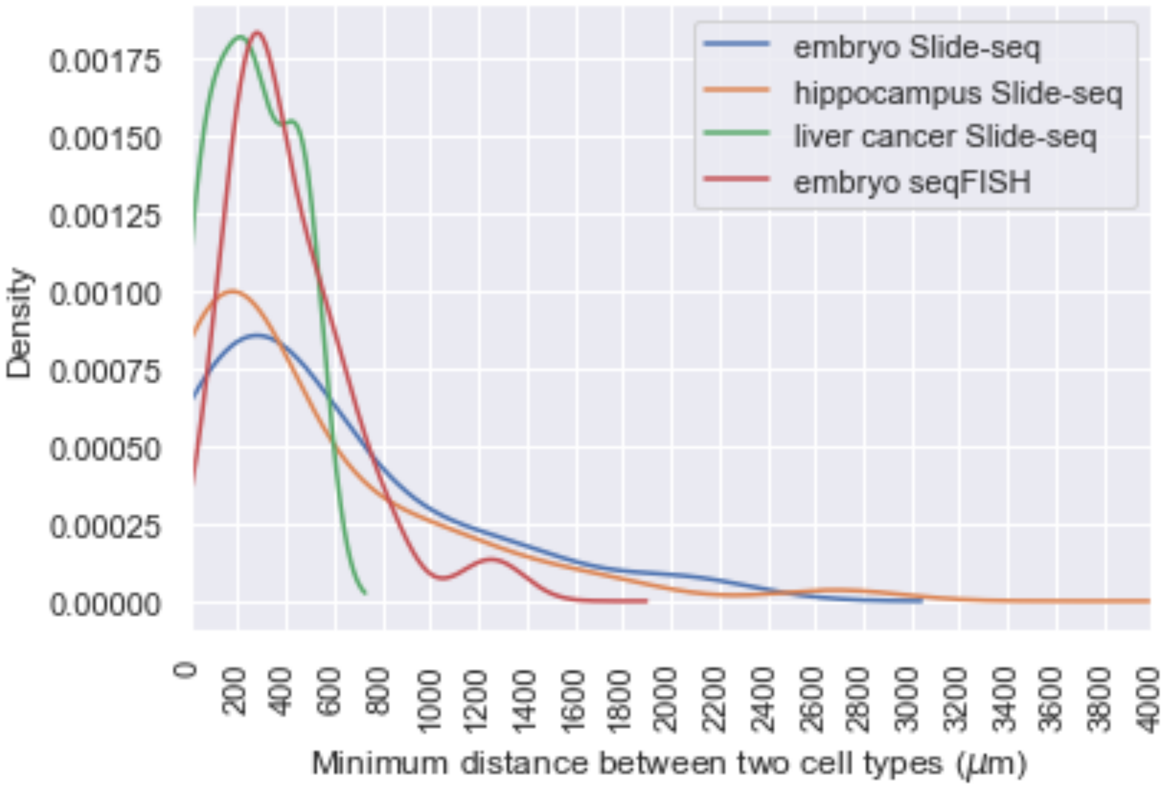

**Fig. S7. Distributions of minimum distances between two interacting cell types detected by NicheNet.** The minimum distance is defined as a distance from a sender cell to the nearest receiver cell between two interacting cell types identified by NicheNet. For each sender cell, the minimum distance is calculated and then the distance values are averaged. The distribution of each dataset is obtained from the averaged minimum distances for the entire cell type pairs of a sender and a receiver. The distributions are estimated by kernel density estimation.

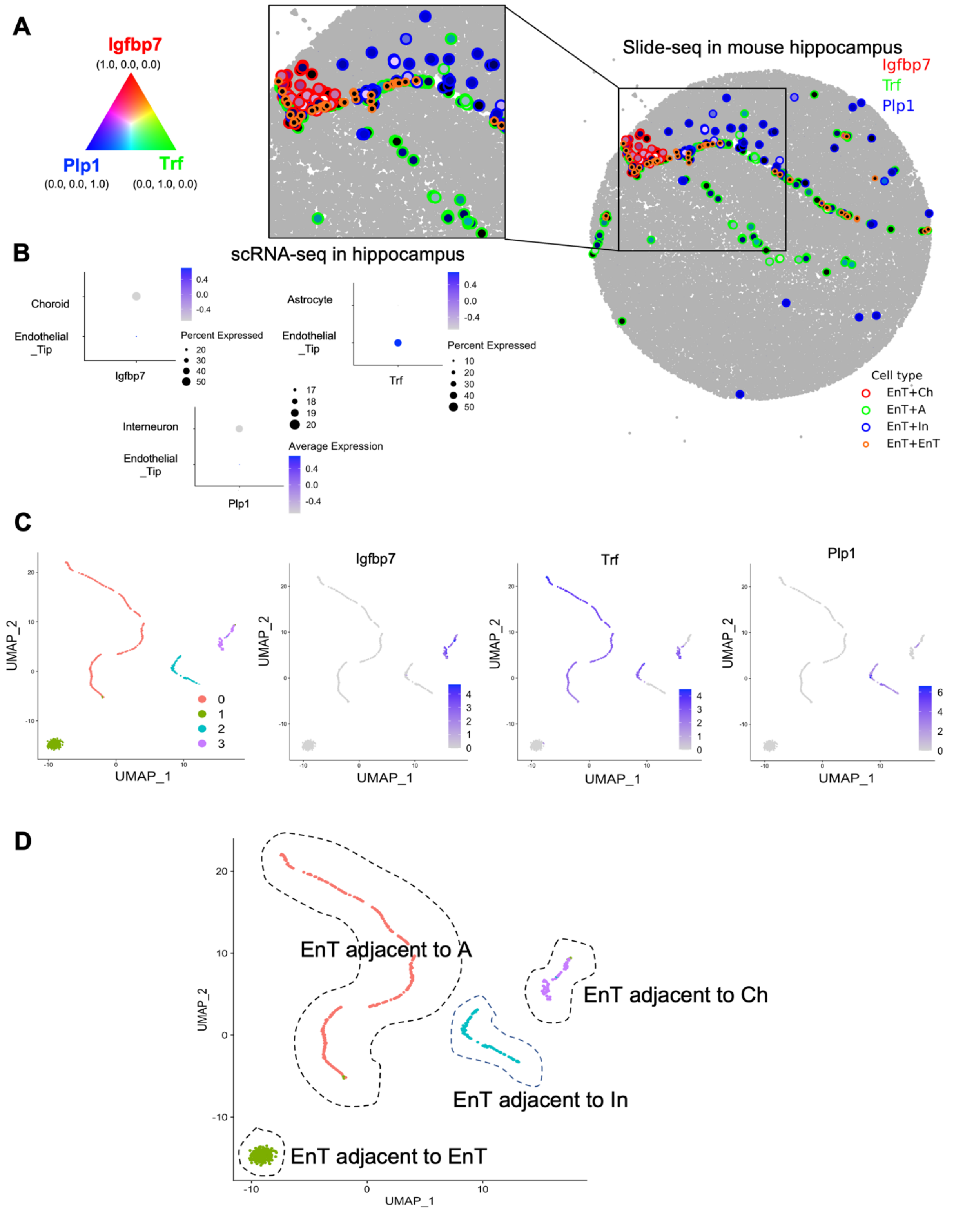
A
Igfbp7
(1.0, 0.0, 0.0)
Plp1
(0.0, 0.0, 1.0)
Trf
(0.0, 1.0, 0.0)
Slide-seq in mouse hippocampus
Igfbp7
Trf
Plp1
Cell type
EnT+Ch
EnT+A
EnT+In
EnT+EnT
B
scRNA-seq in hippocampus
Choroid
Endothelial
_Tip
Igfbp7
Percent Expressed
Astrocyte
Trf
Interneuron
Plp1
Average Expression
C
UMAP_1
UMAP_2
Igfbp7
Trf
Plp1
D
EnT adjacent to A
EnT adjacent to Ch
EnT adjacent to In
EnT adjacent to EnT
UMAP_1
UMAP_2

**Fig. S8. Heterogeneity of endothelial tip (EnT) cells in the mouse hippocampus Slide-seq data. A.** Neighboring cell type-dependent gene expression of EnT cells. EnT cells dominantly express *Igfbp7* (red) when proximal to Choroid (EnT+Ch), *Trf* (green) when proximal to Astrocyte (EnT+A), and *Plp1* (blue) when proximal to Interneuron (EnT+In). **B.** Expression of neighbor-dependent genes in the mouse hippocampus scRNA-seq data. It confirms that the three genes are expressed from EnT. **C.** UMAP of EnT cells. 4 clusters were obtained from clustering analysis: Cluster 0 to 3. *Igfbp7* is mostly expressed in Cluster 3, *Trf* in Cluster 0, *Plp1* in Cluster 2, and none of them is expressed in Cluster 1. **D.** Heterogeneity of EnT cells explained by niche-specific gene expression. Cluster 3 is EnT cells adjacent to Ch, Cluster 0 is EnT adjacent to A, Cluster 2 is EnT adjacent to In, and Cluster 0 is EnT adjacent to another EnT.

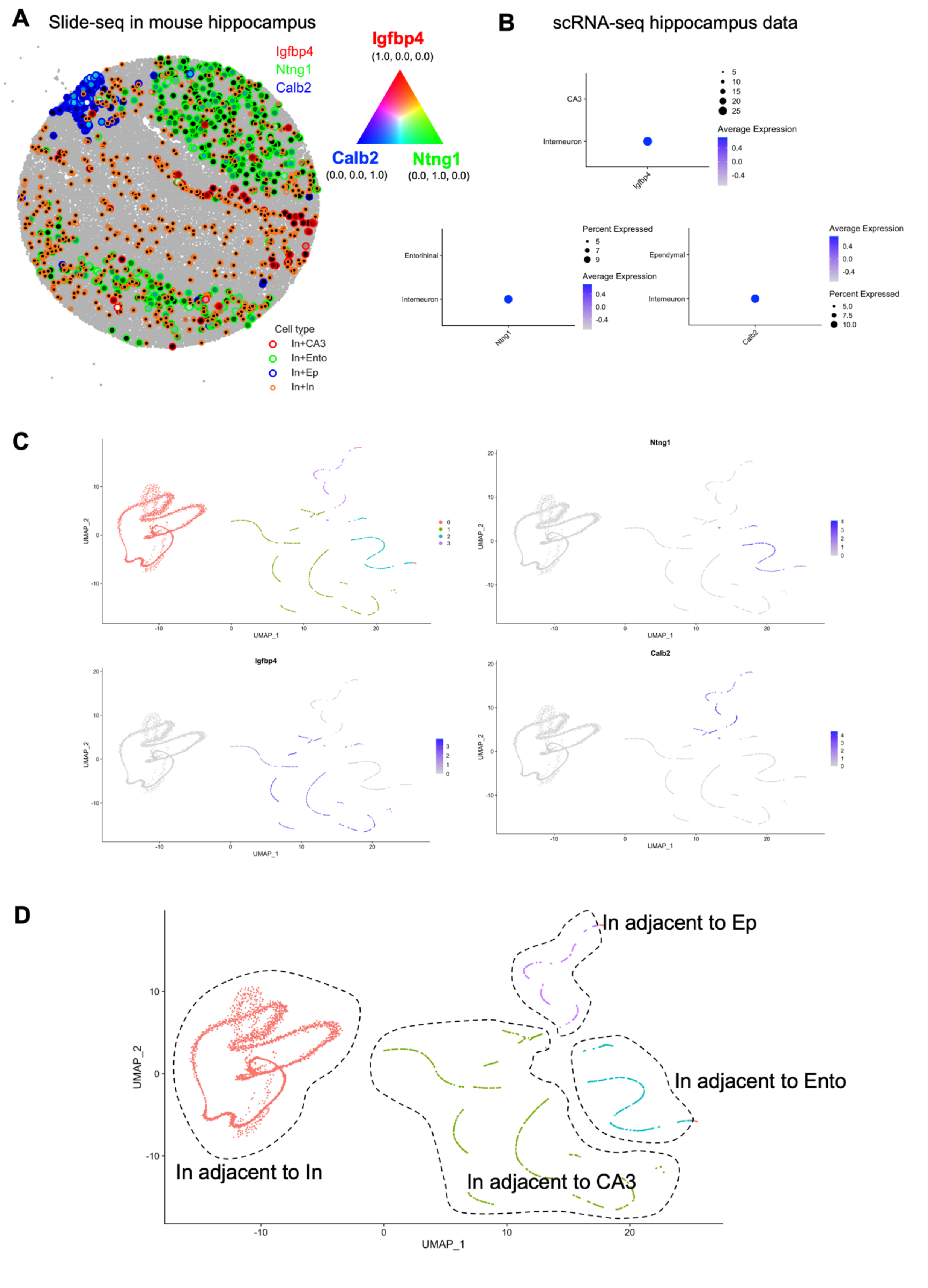

A
Slide-seq in mouse hippocampus
Igfbp4
Ntng1
Calb2
Igfbp4
(1.0, 0.0, 0.0)
Calb2
(0.0, 0.0, 1.0)
Ntng1
(0.0, 1.0, 0.0)
Cell type
In+CA3
In+Ento
In+Ep
In+In
B
scRNA-seq hippocampus data
CA3
Interneuron
Igfbp4
Average Expression
Percent Expressed
Entorihinal
Interneuron
Ntng1
Ependymal
Interneuron
Calb2
C
UMAP_1
UMAP_2
Ntng1
Igfbp4
Calb2
D
In adjacent to Ep
In adjacent to Ento
In adjacent to In
In adjacent to CA3
UMAP_1
UMAP_2

**Fig. S9. Heterogeneity of Interneuron (In) cells in the mouse hippocampus Slide-seq data. A.** Neighboring cell type-dependent gene expression of Interneuron cells. Interneuron cells dominantly express *Igfbp4* (red) when proximal to CA3 (In+CA3), *Ntng1* (green) when proximal to Entorhinal cells (In+Ento), and *Calb2* (blue) when proximal to Ependymal cells (In+Ep). **B.** Expression of neighbor-dependent genes in the mouse hippocampus scRNA-seq data. It was confirmed that the three genes are mostly expressed from Interneuron cells. **C.** UMAP of Interneuron cells. 4 clusters were obtained from clustering analysis: Cluster 0 to 3. *Igfbp4* is mostly expressed in Cluster 1, *Ntng1* in Cluster 2, *Calb2* in Cluster 3, and none of them is expressed in Cluster 0. **D.** Heterogeneity of Interneuron cells explained by niche-specific gene expression. It can be inferred from **(C)** that Cluster 1 is Interneuron cells adjacent to CA3, Cluster 2 is Interneuron adjacent to Ento, Cluster 3 is Interneuron adjacent to Ep, and Cluster 0 is Interneuron adjacent to another Interneuron.

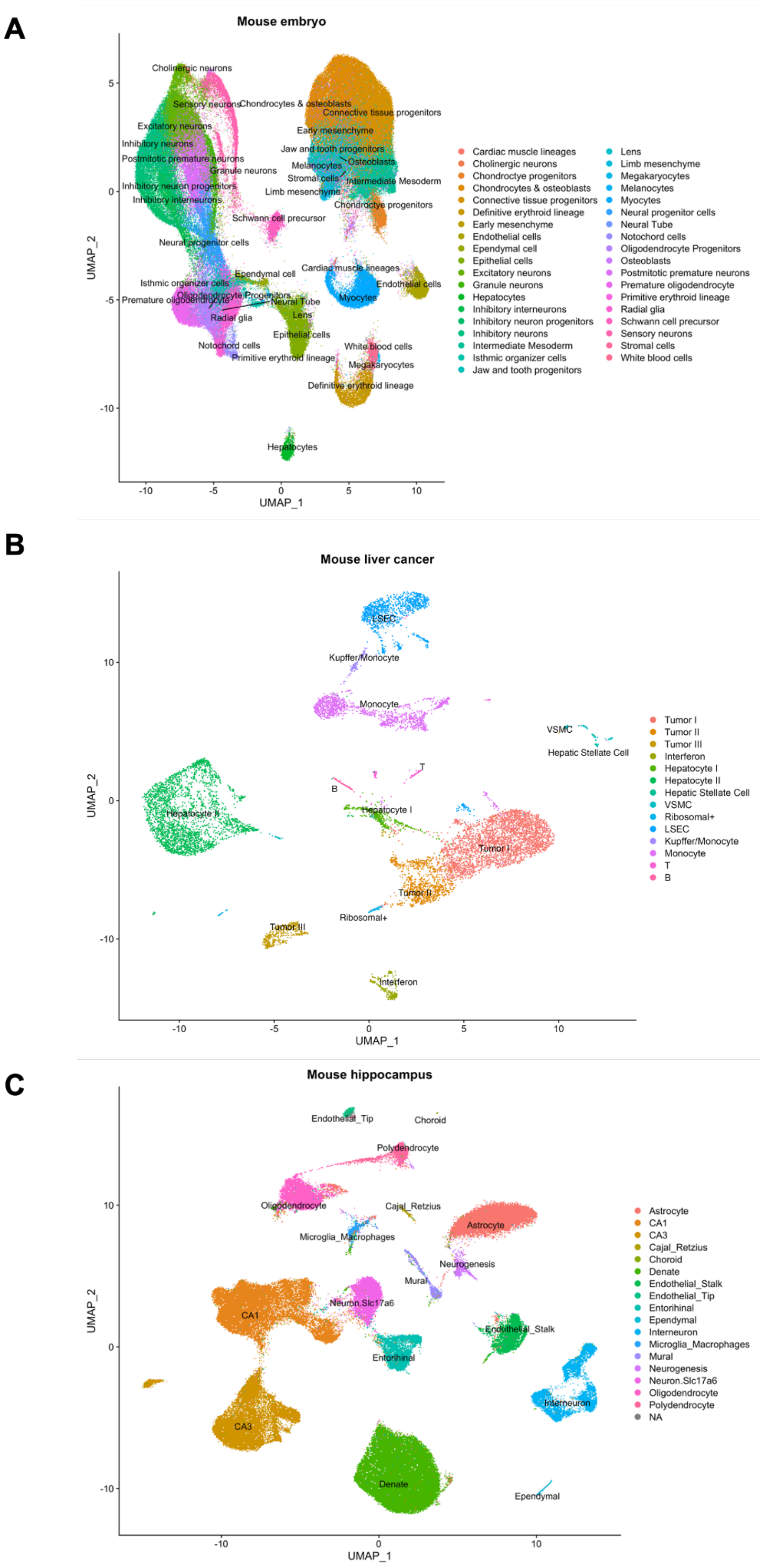
A
Mouse embryo
UMAP_1
UMAP_2
Cardiac muscle lineages
Cholinergic neurons
Chondrocyte progenitors
Chondrocytes & osteoblasts
Connective tissue progenitors
Definitive erythroid lineage
Early mesenchyme
Endothelial cells
Ependymal cell
Epithelial cells
Excitatory neurons
Granule neurons
Hepatocytes
Inhibitory interneurons
Inhibitory neuron progenitors
Inhibitory neurons
Intermediate Mesoderm
Isthmic organizer cells
Jaw and tooth progenitors
Lens
Limb mesenchyme
Megakaryocytes
Melanocytes
Myocytes
Neural progenitor cells
Neural Tube
Notochord cells
Oligodendrocyte Progenitors
Osteoblasts
Postmitotic premature neurons
Premature oligodendrocyte
Primitive erythroid lineage
Radial glia
Schwann cell precursor
Sensory neurons
Stromal cells
White blood cells
B
Mouse liver cancer
UMAP_1
UMAP_2
Tumor I
Tumor II
Tumor III
Interferon
Hepatocyte I
Hepatocyte II
Hepatic Stellate Cell
VSMC
Ribosomal+
LSEC
Kupffer/Monocyte
Monocyte
T
B
C
Mouse hippocampus
UMAP_1
UMAP_2
Astrocyte
CA1
CA3
Cajal_Retzius
Choroid
Denate
Endothelial_Stalk
Endothelial_Tip
Entorihinal
Ependymal
Interneuron
Microglia_Macrophages
Mural
Neurogenesis
Neuron.Slc17a6
Oligodendrocyte
Polydendrocyte
NA

**Fig. S10. UMAP projections for single cell/nucleus data. A.** UMAP of scRNA-seq data from a mouse embryo at E12.5. It consists of 37 cell types. **B.** UMAP of snRNA-seq data from mouse liver cancer. It has 14 cell types. **C.** UMAP of scRNA-seq data from mouse hippocampus. It is composed of 17 cell types.

**A**

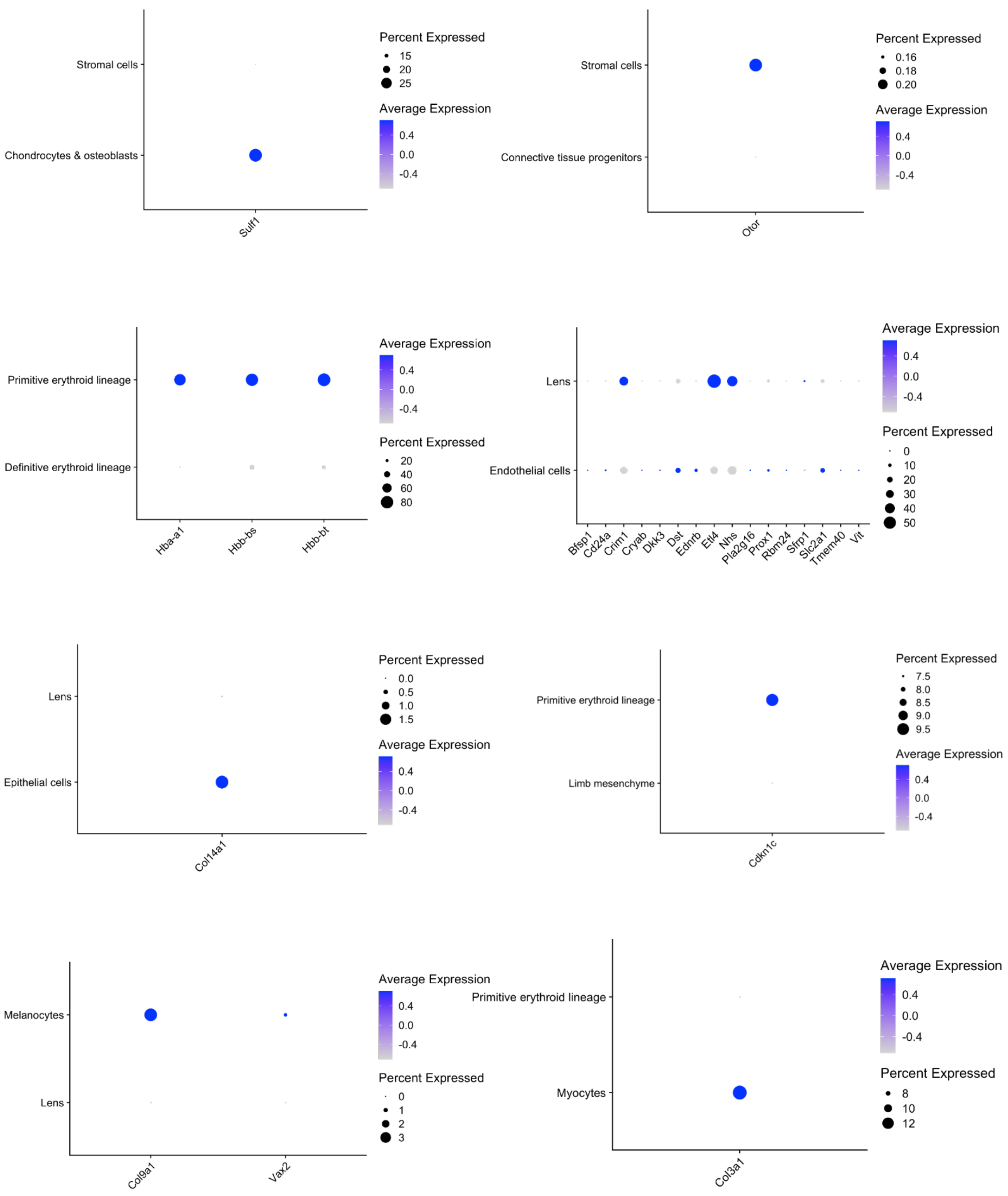

Stromal cells
Chondrocytes & osteoblasts
Sulf1
Percent Expressed
15
20
25
Average Expression
0.4
0.0
-0.4
Stromal cells
Connective tissue progenitors
Otor
Percent Expressed
0.16
0.18
0.20
Average Expression
0.4
0.0
-0.4
Primitive erythroid lineage
Definitive erythroid lineage
Hba-a1
Hbb-bs
Hbb-bt
Average Expression
0.4
0.0
-0.4
Percent Expressed
20
40
60
80
Lens
Endothelial cells
Bfsp1
Cd24a
Crim1
Cryab
Dkk3
Dst
Ednrb
Etl4
Nhs
Pla2g16
Prox1
Rbm24
Sfrp1
Slc2a1
Tmem40
Vit
Average Expression
0.4
0.0
-0.4
Percent Expressed
0
10
20
30
40
50
Lens
Epithelial cells
Col14a1
Percent Expressed
0.0
0.5
1.0
1.5
Average Expression
0.4
0.0
-0.4
Primitive erythroid lineage
Limb mesenchyme
Cdkn1c
Percent Expressed
7.5
8.0
8.5
9.0
9.5
Average Expression
0.4
0.0
-0.4
Melanocytes
Lens
Col9a1
Vax2
Average Expression
0.4
0.0
-0.4
Percent Expressed
0
1
2
3
Primitive erythroid lineage
Myocytes
Col3a1
Average Expression
0.4
0.0
-0.4
Percent Expressed
8
10
12

**B**

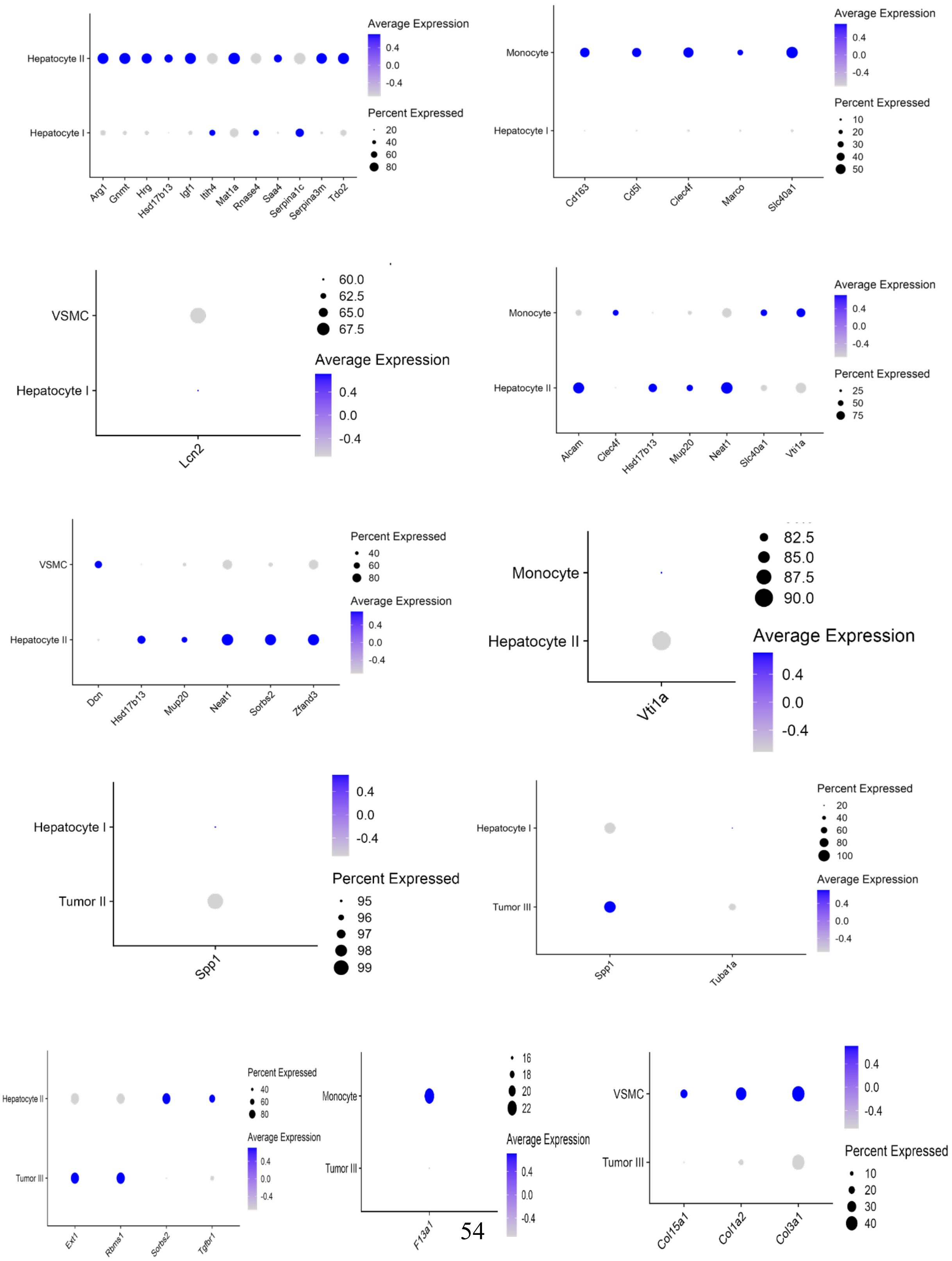
Hepatocyte II
Hepatocyte I
Arg1
Gnmt
Hrg
Hsd17b13
Igf1
Itih4
Mat1a
Rnase4
Saa4
Serpina1c
Serpina3m
Tdo2
Average Expression
0.4
0.0
-0.4
Percent Expressed
20
40
60
80
Monocyte
Hepatocyte I
Cd163
Cd5l
Clec4f
Marco
Slc40a1
Average Expression
Percent Expressed
10
20
30
40
50
VSMC
Hepatocyte I
Lcn2
60.0
62.5
65.0
67.5
Average Expression
Monocyte
Hepatocyte II
Alcam
Clec4f
Hsd17b13
Mup20
Neat1
Slc40a1
Vti1a
Percent Expressed
25
50
75
VSMC
Hepatocyte II
Dcn
Hsd17b13
Mup20
Neat1
Sorbs2
Zfand3
Percent Expressed
40
60
80
Average Expression
Monocyte
Hepatocyte II
Vti1a
82.5
85.0
87.5
90.0
Average Expression
Hepatocyte I
Tumor II
Spp1
Percent Expressed
95
96
97
98
99
Hepatocyte I
Tumor III
Spp1
Tuba1a
Percent Expressed
20
40
60
80
100
Average Expression
Hepatocyte II
Tumor III
Ext1
Rbms1
Sorbs2
Tgfbr1
Percent Expressed
40
60
80
Average Expression
Monocyte
Tumor III
F13a1
16
18
20
22
Average Expression
VSMC
Tumor III
Col15a1
Col1a2
Col3a1
Percent Expressed
10
20
30
40

**c**

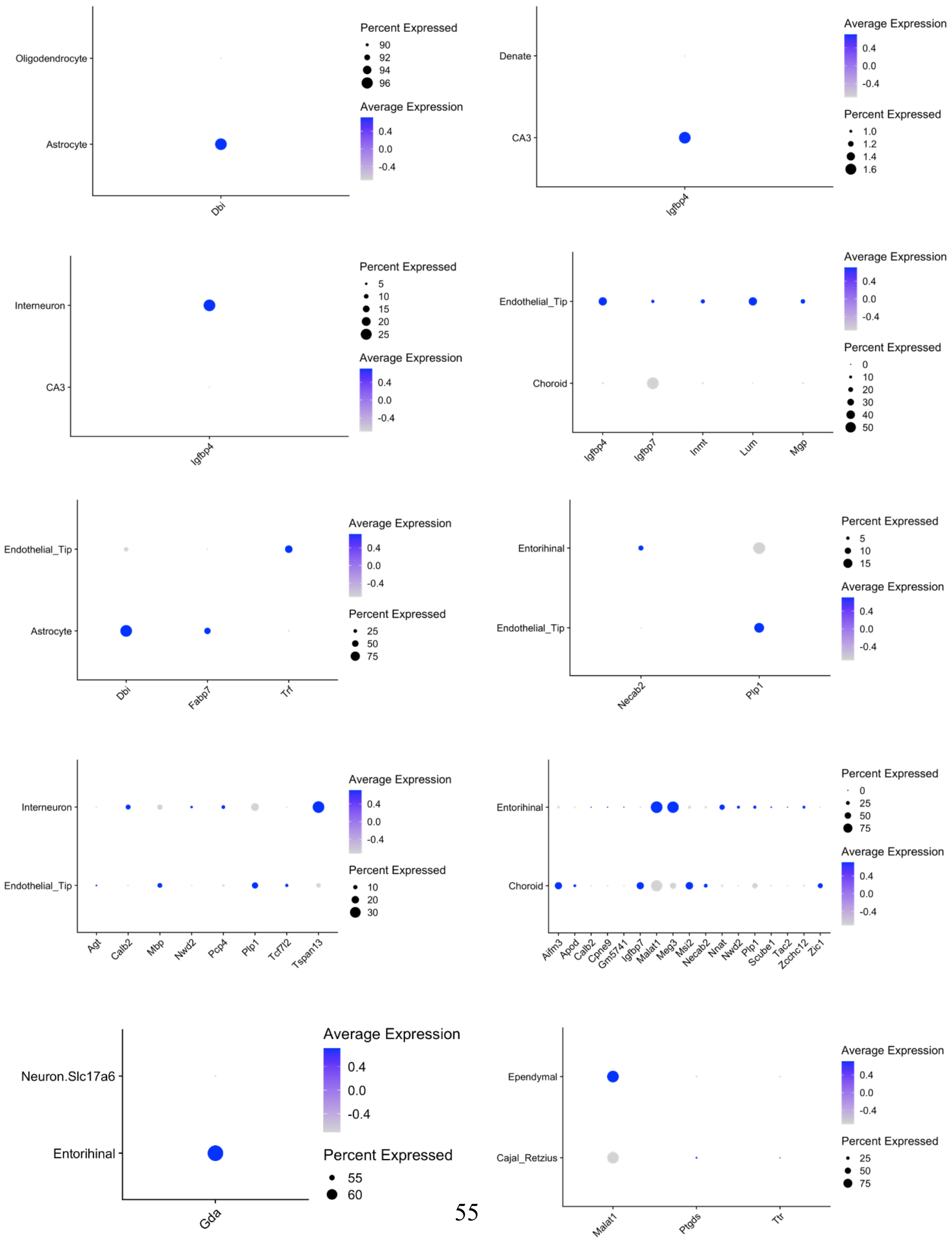
Oligodendrocyte
Astrocyte
Dbi
Percent Expressed
90
92
94
96
Average Expression
0.4
0.0
-0.4
Denate
CA3
Igfbp4
Average Expression
0.4
0.0
-0.4
Percent Expressed
1.0
1.2
1.4
1.6
Interneuron
CA3
Igfbp4
Percent Expressed
5
10
15
20
25
Average Expression
0.4
0.0
-0.4
Endothelial_Tip
Choroid
Igfbp4
Igfbp7
Inmt
Lum
Mgp
Average Expression
0.4
0.0
-0.4
Percent Expressed
0
10
20
30
40
50
Endothelial_Tip
Astrocyte
Dbi
Fabp7
Trf
Average Expression
0.4
0.0
-0.4
Percent Expressed
25
50
75
Entorihinal
Endothelial_Tip
Necab2
Plp1
Percent Expressed
5
10
15
Average Expression
0.4
0.0
-0.4
Interneuron
Endothelial_Tip
Agt
Calb2
Mbp
Nwd2
Pcp4
Plp1
Tcf7l2
Tspan13
Average Expression
0.4
0.0
-0.4
Percent Expressed
10
20
30
Entorihinal
Choroid
Aifm3
Apod
Calb2
Cpne9
Gm5741
Igfbp7
Malat1
Meg3
Msi2
Necab2
Nnat
Nwd2
Plp1
Scube1
Tac2
Zcchc12
Zic1
Percent Expressed
0
25
50
75
Average Expression
0.4
0.0
-0.4
Neuron.Slc17a6
Entorihinal
Gda
Average Expression
0.4
0.0
-0.4
Percent Expressed
55
60
Ependymal
Cajal_Retzius
Malat1
Ptgds
Ttr
Average Expression
0.4
0.0
-0.4
Percent Expressed
25
50
75

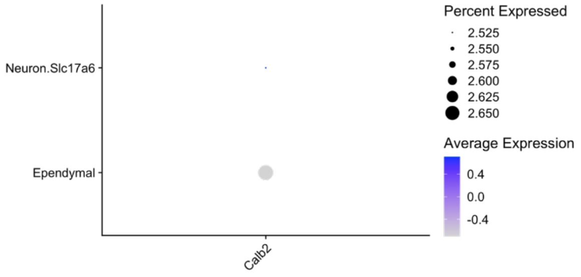
Percent Expressed
2.525
2.550
2.575
2.600
2.625
2.650
Average Expression
0.4
0.0
-0.4
Neuron.Slc17a6
Ependymal
Calb2

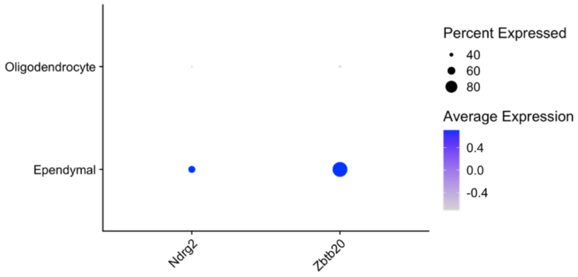
Percent Expressed
40
60
80
Average Expression
0.4
0.0
-0.4
Oligodendrocyte
Ependymal
Ndrg2
Zbtb20

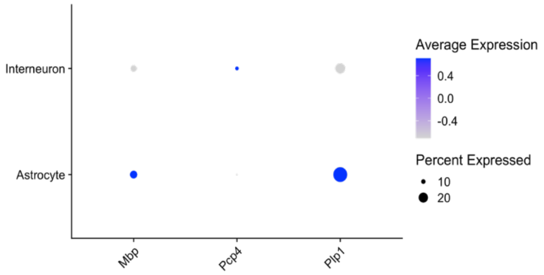
Average Expression
0.4
0.0
-0.4
Percent Expressed
10
20
Interneuron
Astrocyte
Mbp
Pcp4
Plp1

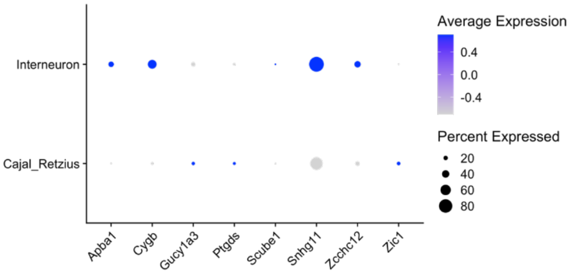
Average Expression
0.4
0.0
-0.4
Percent Expressed
20
40
60
80
Interneuron
Cajal_Retzius
Apba1
Cygb
Gucy1a3
Ptgds
Scube1
Snhg11
Zcchc12
Zic1

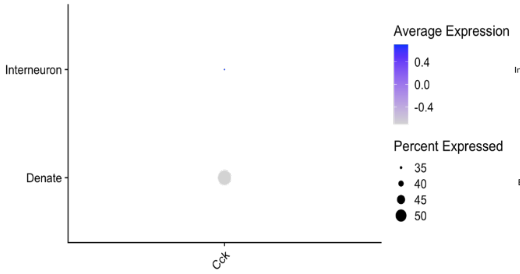
Average Expression
0.4
0.0
-0.4
Percent Expressed
35
40
45
50
Interneuron
Denate
Cck

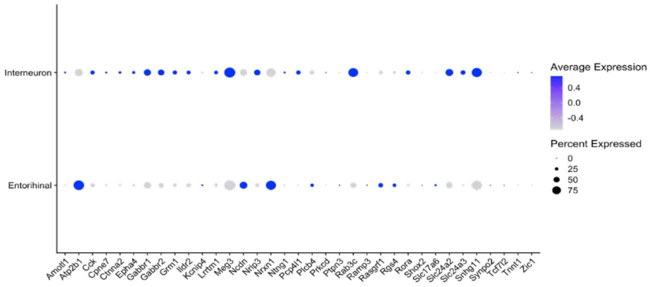
Average Expression
0.4
0.0
-0.4
Percent Expressed
0
25
50
75
Interneuron
Entorihinal
Amotl1
Atp2b1
Cck
Cpne7
Ctnna2
Epha4
Gabbr1
Gabbr2
Grm1
Ildr2
Kcnip4
Lrrtm1
Meg3
Ncdn
Nrip3
Nrxn1
Ntng1
Pcp4l1
Plcb4
Prkcd
Ptpn3
Rab3c
Ramp3
Rasgrf1
Rgs4
Rora
Shox2
Slc17a6
Slc24a2
Slc24a3
Snhg11
Synpo2
Tcf7l2
Tnnt1
Zic1

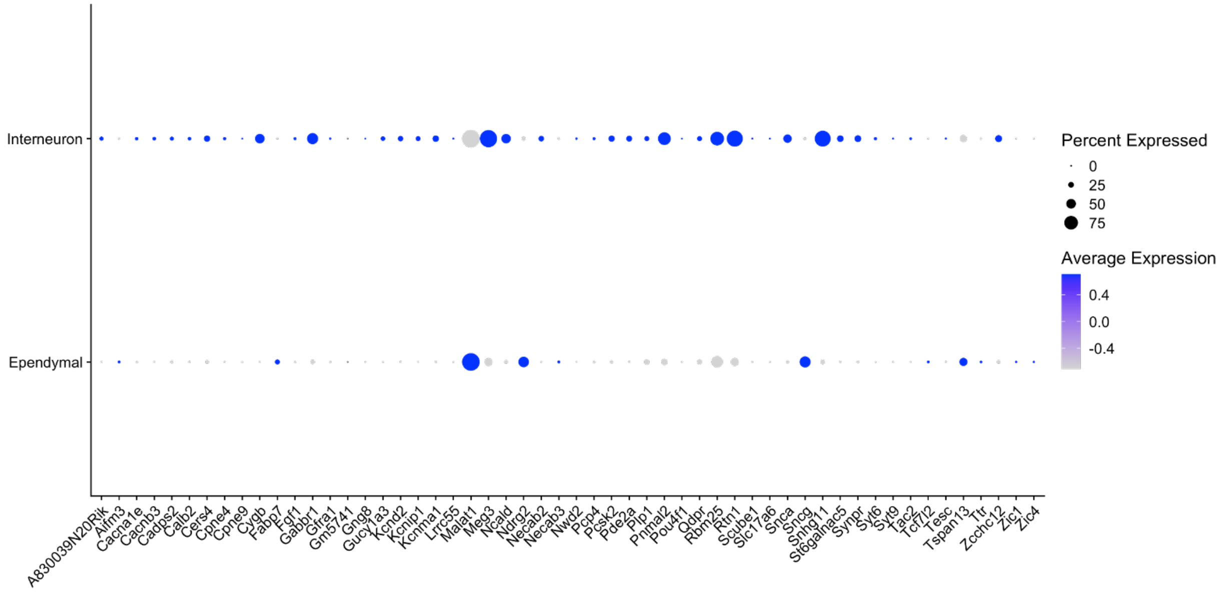
Percent Expressed
0
25
50
75
Average Expression
0.4
0.0
-0.4
Interneuron
Ependymal
A830039N20Rik
Aifm3
Cacna1e
Cacnb3
Cadps2
Calb2
Cers4
Cpne4
Cpne9
Cygb
Fabp7
Fgf1
Gabbr1
Gfra1
Gm5741
Gng8
Gucy1a3
Kcnd2
Kcnip1
Kcnma1
Lrrc55
Malat1
Meg3
Ncald
Ndrg2
Necab2
Necab3
Nwd2
Pcp4
Pcsk2
Pde2a
Plp1
Pnmal2
Pou4f1
Qdpr
Rbm25
Rtn1
Scube1
Slc17a6
Snca
Sncg
Snhg11
St6galnac5
Synpr
Syt6
Syt9
Tac2
Tcf7l2
Tesc
Tspan13
Ttr
Zcchc12
Zic1
Zic4

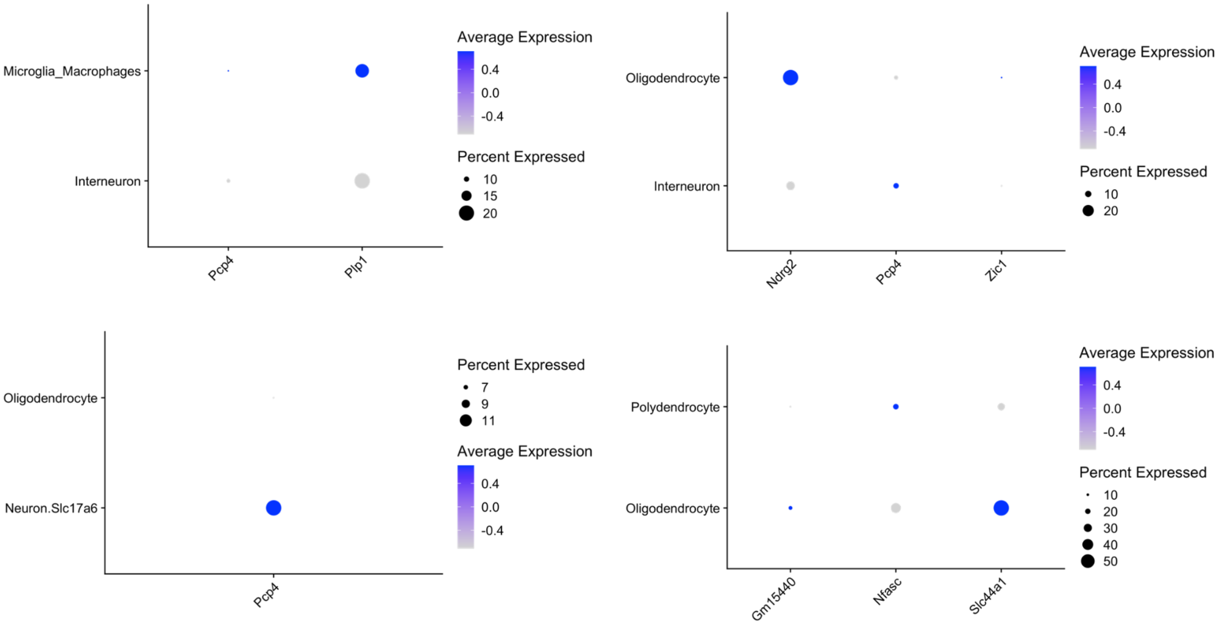

**Fig. S11. Origins of the expression of neighbor-dependent genes identified from single cell/nucleus data. A.** Expression of neighbor-dependent genes in the mouse embryo scRNA-seq data. The genes are the 28 up-regulated genes identified from the 9 heterotypic pairs in the mouse embryo Slide-seq data. **B.** Expression of neighbor-dependent genes in the mouse liver cancer snRNA-seq data. The genes are the 42 up-regulated genes identified from the 10 heterotypic pairs in the mouse liver cancer Slide-seq data. **C.** Expression of neighbor-dependent genes in the mouse hippocampus scRNA-seq data. The genes are the 155 up-regulated genes identified from the 21 heterotypic pairs in the mouse hippocampus Slide-seq data.

#### Supplementary Tables

#### Table S1A. Up-regulated genes identified from the mouse embryo seqFISH data

| Centered cell type/neighboring cell type | Up-regulated_gene | Homotypic1_logFC | Homotypic1_p.value | Homotypic1_fdr |
|---|---|---|---|---|
| Cardiomyocytes/Endothelium | Acvr1 | 0.57025 | 0.0027343 | 0.03943 |
| Cardiomyocytes/Endothelium | Akr1c19 | 0.61586 | 0.00055059 | 0.014436 |
| Cardiomyocytes/Endothelium | Cdh5 | 0.9569 | 1.77E-05 | 0.0012776 |
| Cardiomyocytes/Endothelium | Foxh1 | 0.66256 | 0.0029917 | 0.041087 |
| Cardiomyocytes/Endothelium | Fst | 0.61521 | 0.0014352 | 0.022996 |
| Cardiomyocytes/Endothelium | Hcn4 | 0.92915 | 0.00071837 | 0.015937 |
| Cardiomyocytes/Endothelium | Igf1 | 0.63315 | 0.00093871 | 0.018049 |
| Cardiomyocytes/Endothelium | Kmt2d | 0.8302 | 0.00046496 | 0.0149 |
| Cardiomyocytes/Endothelium | Pdgfa | 0.80587 | 0.00015449 | 0.0074263 |
| Cardiomyocytes/Endothelium | Pdgfra | 0.73204 | 0.0007913 | 0.016301 |
| Cardiomyocytes/Endothelium | Tbx5 | 1.1385 | 1.30E-05 | 0.0012486 |
| Cardiomyocytes/Mixed-mesenchymal-mesoderm | Bmp2 | 1.0469 | 0.00035217 | 0.017327 |
| Cardiomyocytes/Splanchnic-mesoderm | Aplnr | 1.4793 | 0.00033896 | 0.0064747 |
| Cardiomyocytes/Splanchnic-mesoderm | Car7 | 0.95652 | 0.0041978 | 0.034365 |
| Cardiomyocytes/Splanchnic-mesoderm | Cbfa2t3 | 0.99962 | 0.0065607 | 0.0482 |
| Cardiomyocytes/Splanchnic-mesoderm | Dusp6 | 1.1508 | 0.00010956 | 0.0031392 |
| Cardiomyocytes/Splanchnic-mesoderm | Fgfr2 | 1.0394 | 0.0029774 | 0.029418 |
| Cardiomyocytes/Splanchnic-mesoderm | Foxc2 | 1.1582 | 0.0032002 | 0.028654 |
| Cardiomyocytes/Splanchnic-mesoderm | Foxf1 | 2.3885 | 5.89E-07 | 3.37E-05 |
| Cardiomyocytes/Splanchnic-mesoderm | Gata6 | 0.73845 | 0.0024848 | 0.026368 |
| Cardiomyocytes/Splanchnic-mesoderm | Hapln1 | 1.2094 | 0.0002318 | 0.004744 |
| Cardiomyocytes/Splanchnic-mesoderm | Kmt2d | 1.185 | 0.0016279 | 0.020279 |
| Cardiomyocytes/Splanchnic-mesoderm | Lin28a | 0.96764 | 0.00014761 | 0.0035245 |
| Cardiomyocytes/Splanchnic-mesoderm | Marcks | 1.3539 | 0.00049587 | 0.0083576 |
| Cardiomyocytes/Splanchnic-mesoderm | Meis1 | 1.2147 | 0.00064699 | 0.010299 |
| Cardiomyocytes/Splanchnic-mesoderm | Meis2 | 1.519 | 3.35E-05 | 0.0012008 |
| Cardiomyocytes/Splanchnic-mesoderm | Mkrn1 | 0.80871 | 0.0037463 | 0.032528 |
| Cardiomyocytes/Splanchnic-mesoderm | Pcgf2 | 0.82535 | 0.0024973 | 0.025555 |
| Cardiomyocytes/Splanchnic-mesoderm | Pdgfa | 0.91687 | 0.0056973 | 0.042959 |
| Cardiomyocytes/Splanchnic-mesoderm | Pdgfra | 1.7755 | 2.88E-07 | 2.06E-05 |
| Cardiomyocytes/Splanchnic-mesoderm | Sfrp2 | 1.039 | 0.0053253 | 0.042384 |
| Cardiomyocytes/Splanchnic-mesoderm | Smim1 | 1.0542 | 0.0030598 | 0.028281 |
| Cardiomyocytes/Splanchnic-mesoderm | Tbx1 | 1.3839 | 0.0011375 | 0.014815 |
| Cardiomyocytes/Splanchnic-mesoderm | Tbx5 | 1.2803 | 0.0017204 | 0.020539 |
| Cardiomyocytes/Splanchnic-mesoderm | Tead2 | 1.3272 | 7.16E-18 | 2.05E-15 |
| Cranial-mesoderm/Gut-tube | Fgfr2 | 1.3417 | 0.00036116 | 0.037742 |
| Cranial-mesoderm/Gut-tube | Foxf1 | 1.8591 | 1.33E-07 | 4.17E-05 |
| Cranial-mesoderm/Gut-tube | Krt18 | 1.2869 | 0.00076703 | 0.034353 |
| Cranial-mesoderm/Gut-tube | Lin28a | 1.3488 | 0.00065793 | 0.034378 |
| Cranial-mesoderm/Gut-tube | Podxl | 1.4285 | 0.00038342 | 0.030052 |
| Dermomyotome/Presomitic-mesoderm | Cxcl12 | 1.5273 | 0.0014337 | 0.025487 |
| Dermomyotome/Presomitic-mesoderm | Dll3 | 2.101 | 0.00073385 | 0.017395 |
| Dermomyotome/Presomitic-mesoderm | Hoxa1 | 2.3938 | 0.000147 | 0.0069689 |
| Dermomyotome/Presomitic-mesoderm | Hoxa7 | 2.2709 | 0.00038549 | 0.012183 |
| Dermomyotome/Presomitic-mesoderm | Hoxb8 | 1.9937 | 0.0014806 | 0.024774 |
| Dermomyotome/Presomitic-mesoderm | Lef1 | 1.0533 | 0.0013442 | 0.027311 |
| Dermomyotome/Presomitic-mesoderm | Lfng | 2.1943 | 0.0021417 | 0.033844 |
| Dermomyotome/Presomitic-mesoderm | Marcks | 2.374 | 0.00086898 | 0.019014 |
| Dermomyotome/Presomitic-mesoderm | Meox1 | 0.57023 | 0.0001314 | 0.0074754 |
| Dermomyotome/Presomitic-mesoderm | Prrx2 | 2.3661 | 0.0002373 | 0.0096426 |
| Dermomyotome/Presomitic-mesoderm | Ptn | 1.6331 | 9.41E-05 | 0.0089227 |
| Forebrain-Midbrain-Hindbrain/Cranial-mesoderm | Akr1c19 | 1.6853 | 9.17E-10 | 3.77E-08 |
| Forebrain-Midbrain-Hindbrain/Cranial-mesoderm | Bmp7 | 0.78127 | 0.0070157 | 0.036858 |

| | | | | |
|---|---|---|---|---|
| Forebrain-Midbrain-Hindbrain/Cranial-mesoderm | Col26a1 | 0.90275 | 0.0013329 | 0.0091419 |
| Forebrain-Midbrain-Hindbrain/Cranial-mesoderm | Dnajb13 | 0.64716 | 0.0077114 | 0.038859 |
| Forebrain-Midbrain-Hindbrain/Cranial-mesoderm | Dnmt3a | 1.042 | 0.00063137 | 0.0051966 |
| Forebrain-Midbrain-Hindbrain/Cranial-mesoderm | Dusp6 | 2.4013 | 5.47E-09 | 1.50E-07 |
| Forebrain-Midbrain-Hindbrain/Cranial-mesoderm | En1 | 1.5571 | 0.00014681 | 0.0017262 |
| Forebrain-Midbrain-Hindbrain/Cranial-mesoderm | Foxa1 | 2.3251 | 3.37E-09 | 1.04E-07 |
| Forebrain-Midbrain-Hindbrain/Cranial-mesoderm | Foxa2 | 2.8649 | 5.29E-21 | 6.54E-19 |
| Forebrain-Midbrain-Hindbrain/Cranial-mesoderm | Gbx2 | 1.0603 | 0.00065576 | 0.0052232 |
| Forebrain-Midbrain-Hindbrain/Cranial-mesoderm | Gsn | 0.82967 | 0.0028433 | 0.018002 |
| Forebrain-Midbrain-Hindbrain/Cranial-mesoderm | Hoxb3 | 0.73195 | 0.0057778 | 0.032424 |
| Forebrain-Midbrain-Hindbrain/Cranial-mesoderm | Igfbp3 | 1.1881 | 0.00045169 | 0.0041308 |
| Forebrain-Midbrain-Hindbrain/Cranial-mesoderm | Itga3 | 0.79434 | 0.0090043 | 0.044467 |
| Forebrain-Midbrain-Hindbrain/Cranial-mesoderm | Kitl | 0.73753 | 0.0096221 | 0.046586 |
| Forebrain-Midbrain-Hindbrain/Cranial-mesoderm | Krt18 | 1.0378 | 0.0008533 | 0.0063847 |
| Forebrain-Midbrain-Hindbrain/Cranial-mesoderm | Marcks | 1.2627 | 8.20E-07 | 1.69E-05 |
| Forebrain-Midbrain-Hindbrain/Cranial-mesoderm | Myh9 | 0.81695 | 0.0072151 | 0.037116 |
| Forebrain-Midbrain-Hindbrain/Cranial-mesoderm | Prrx2 | 0.6626 | 0.0037442 | 0.022549 |
| Forebrain-Midbrain-Hindbrain/Cranial-mesoderm | Shh | 3.0708 | 1.34E-22 | 3.32E-20 |
| Forebrain-Midbrain-Hindbrain/Cranial-mesoderm | Sox2 | 0.82182 | 2.49E-08 | 6.15E-07 |
| Forebrain-Midbrain-Hindbrain/Cranial-mesoderm | Tcf7l1 | 1.0851 | 4.90E-05 | 0.00067257 |
| Forebrain-Midbrain-Hindbrain/Cranial-mesoderm | Tmem108 | 0.80003 | 0.0044496 | 0.025551 |
| Forebrain-Midbrain-Hindbrain/Cranial-mesoderm | Tmem119 | 0.6981 | 0.0069075 | 0.037078 |
| Forebrain-Midbrain-Hindbrain/Definitive-endoderm | Apln | 0.79416 | 0.0061697 | 0.02286 |
| Forebrain-Midbrain-Hindbrain/Definitive-endoderm | Atp1b1 | 0.83783 | 0.0068623 | 0.024337 |
| Forebrain-Midbrain-Hindbrain/Definitive-endoderm | Bmp7 | 1.3593 | 0.00014735 | 0.0010758 |
| Forebrain-Midbrain-Hindbrain/Definitive-endoderm | Col26a1 | 1.103 | 0.00066356 | 0.0040178 |
| Forebrain-Midbrain-Hindbrain/Definitive-endoderm | Col4a1 | 2.4915 | 7.83E-14 | 1.94E-12 |
| Forebrain-Midbrain-Hindbrain/Definitive-endoderm | Cpm | 0.99361 | 0.001892 | 0.0093937 |
| Forebrain-Midbrain-Hindbrain/Definitive-endoderm | Dnajb13 | 0.83048 | 0.0044637 | 0.017589 |
| Forebrain-Midbrain-Hindbrain/Definitive-endoderm | Dusp6 | 3.6812 | 4.85E-31 | 1.72E-29 |
| Forebrain-Midbrain-Hindbrain/Definitive-endoderm | En1 | 3.6102 | 4.57E-20 | 1.26E-18 |
| Forebrain-Midbrain-Hindbrain/Definitive-endoderm | Etv4 | 0.9129 | 0.0015719 | 0.0088685 |
| Forebrain-Midbrain-Hindbrain/Definitive-endoderm | Foxa1 | 3.4831 | 8.09E-38 | 4.01E-36 |
| Forebrain-Midbrain-Hindbrain/Definitive-endoderm | Foxa2 | 3.7021 | 5.55E-23 | 1.72E-21 |
| Forebrain-Midbrain-Hindbrain/Definitive-endoderm | Furin | 1.6269 | 9.55E-08 | 1.25E-06 |
| Forebrain-Midbrain-Hindbrain/Definitive-endoderm | Gbx2 | 0.941 | 0.0042148 | 0.017153 |
| Forebrain-Midbrain-Hindbrain/Definitive-endoderm | Gmpr | 0.81531 | 0.0045533 | 0.017662 |
| Forebrain-Midbrain-Hindbrain/Definitive-endoderm | Gpc4 | 0.80742 | 0.0052509 | 0.01975 |
| Forebrain-Midbrain-Hindbrain/Definitive-endoderm | Hoxa1 | 0.95607 | 0.0017531 | 0.0092595 |
| Forebrain-Midbrain-Hindbrain/Definitive-endoderm | Hoxb4 | 0.90947 | 0.0030945 | 0.013718 |
| Forebrain-Midbrain-Hindbrain/Definitive-endoderm | Igfbp3 | 1.9515 | 5.89E-06 | 5.42E-05 |
| Forebrain-Midbrain-Hindbrain/Definitive-endoderm | Itga3 | 2.4377 | 6.98E-09 | 1.02E-07 |
| Forebrain-Midbrain-Hindbrain/Definitive-endoderm | Kitl | 2.1643 | 5.86E-09 | 9.09E-08 |
| Forebrain-Midbrain-Hindbrain/Definitive-endoderm | Krt18 | 0.97228 | 0.0023212 | 0.010872 |
| Forebrain-Midbrain-Hindbrain/Definitive-endoderm | Marcks | 0.84996 | 0.0016964 | 0.0091549 |
| Forebrain-Midbrain-Hindbrain/Definitive-endoderm | Myh9 | 0.9937 | 0.0028569 | 0.012895 |
| Forebrain-Midbrain-Hindbrain/Definitive-endoderm | Nrp1 | 1.239 | 1.41E-05 | 0.00012459 |
| Forebrain-Midbrain-Hindbrain/Definitive-endoderm | Per2 | 1.3638 | 0.00010593 | 0.00079686 |
| Forebrain-Midbrain-Hindbrain/Definitive-endoderm | Setd2 | 1.5254 | 4.41E-06 | 4.21E-05 |
| Forebrain-Midbrain-Hindbrain/Definitive-endoderm | Shh | 4.9314 | 5.59E-53 | 4.62E-51 |
| Forebrain-Midbrain-Hindbrain/Definitive-endoderm | Tcf7l1 | 1.0002 | 0.00285 | 0.013102 |
| Forebrain-Midbrain-Hindbrain/Definitive-endoderm | Wnt5a | 1.8417 | 1.28E-10 | 2.44E-09 |
| Forebrain-Midbrain-Hindbrain/Definitive-endoderm | Xist | 0.97889 | 0.0066407 | 0.023892 |
| Forebrain-Midbrain-Hindbrain/Definitive-endoderm | Zfp57 | 0.88459 | 0.0035833 | 0.015337 |

| | | | | |
|---|---|---|---|---|
| Forebrain-Midbrain-Hindbrain/Neural-crest | Hoxb5 | 1.1338 | 0.0024989 | 0.043205 |
| Forebrain-Midbrain-Hindbrain/Neural-crest | Lef1 | 1.3291 | 0.00031939 | 0.0090363 |
| Forebrain-Midbrain-Hindbrain/Neural-crest | Lhx2 | 1.4767 | 7.82E-05 | 0.003042 |
| Forebrain-Midbrain-Hindbrain/Neural-crest | Otx2 | 1.4854 | 0.00020709 | 0.0071612 |
| Forebrain-Midbrain-Hindbrain/Neural-crest | Shh | 1.7975 | 8.46E-07 | 6.58E-05 |
| Forebrain-Midbrain-Hindbrain/Neural-crest | Six3 | 1.0679 | 0.0016337 | 0.029907 |
| Forebrain-Midbrain-Hindbrain/Neural-crest | Tagln | 1.3584 | 0.00040442 | 0.010488 |
| Forebrain-Midbrain-Hindbrain/Spinal-cord | Dll3 | 0.75958 | 3.90E-06 | 6.68E-05 |
| Forebrain-Midbrain-Hindbrain/Spinal-cord | Fgfr3 | 0.46046 | 0.0095755 | 0.030702 |
| Forebrain-Midbrain-Hindbrain/Spinal-cord | Hoxb3 | 0.88602 | 2.29E-08 | 6.71E-07 |
| Forebrain-Midbrain-Hindbrain/Spinal-cord | Hoxb4 | 0.50631 | 9.18E-05 | 0.0008974 |
| Forebrain-Midbrain-Hindbrain/Spinal-cord | Hoxd4 | 0.84656 | 4.88E-08 | 1.25E-06 |
| Forebrain-Midbrain-Hindbrain/Spinal-cord | Sfrp2 | 0.54313 | 0.0020003 | 0.0095457 |
| Gut-tube/Cranial-mesoderm | Bmp7 | 0.9359 | 0.0008048 | 0.0039802 |
| Gut-tube/Cranial-mesoderm | Cdh2 | 1.8662 | 2.02E-13 | 7.84E-12 |
| Gut-tube/Cranial-mesoderm | Chrd | 1.1746 | 0.00011482 | 0.00076256 |
| Gut-tube/Cranial-mesoderm | Col1a1 | 0.63275 | 0.008213 | 0.024793 |
| Gut-tube/Cranial-mesoderm | Col26a1 | 1.5235 | 2.49E-07 | 3.62E-06 |
| Gut-tube/Cranial-mesoderm | Fgfr2 | 0.93986 | 0.00074553 | 0.0037673 |
| Gut-tube/Cranial-mesoderm | Foxa1 | 0.63428 | 0.0097991 | 0.028832 |
| Gut-tube/Cranial-mesoderm | Foxc2 | 1.0869 | 5.72E-05 | 0.00047469 |
| Gut-tube/Cranial-mesoderm | Foxd4 | 0.7216 | 0.0026873 | 0.010075 |
| Gut-tube/Cranial-mesoderm | Fzd2 | 0.81182 | 2.59E-06 | 2.61E-05 |
| Gut-tube/Cranial-mesoderm | Igf1 | 1.7524 | 6.06E-08 | 9.38E-07 |
| Gut-tube/Cranial-mesoderm | Irx1 | 1.3031 | 9.61E-07 | 1.18E-05 |
| Gut-tube/Cranial-mesoderm | Irx2 | 0.85566 | 0.00061815 | 0.0033415 |
| Gut-tube/Cranial-mesoderm | Marcks | 1.381 | 6.68E-07 | 8.63E-06 |
| Gut-tube/Cranial-mesoderm | Mnt | 1.0673 | 8.66E-05 | 0.00062904 |
| Gut-tube/Cranial-mesoderm | Nr2f1 | 0.89099 | 0.0021096 | 0.0083113 |
| Gut-tube/Cranial-mesoderm | Piezo2 | 0.62308 | 0.0060767 | 0.019894 |
| Gut-tube/Cranial-mesoderm | Shisa2 | 1.0268 | 0.00069265 | 0.0036591 |
| Gut-tube/Cranial-mesoderm | Six1 | 1.4028 | 1.62E-06 | 1.88E-05 |
| Gut-tube/Cranial-mesoderm | Smoc2 | 1.8402 | 7.33E-12 | 1.89E-10 |
| Gut-tube/Cranial-mesoderm | Snai1 | 0.82804 | 0.0026427 | 0.01007 |
| Gut-tube/Cranial-mesoderm | Sox2 | 1.6751 | 1.23E-08 | 2.20E-07 |
| Gut-tube/Cranial-mesoderm | Sp5 | 0.71583 | 0.0076451 | 0.023694 |
| Gut-tube/Cranial-mesoderm | Tbx1 | 2.9051 | 2.07E-29 | 2.40E-27 |
| Gut-tube/Cranial-mesoderm | Tcf7l1 | 0.65581 | 0.0017807 | 0.0071365 |
| Gut-tube/Cranial-mesoderm | Tdo2 | 0.58168 | 0.0083463 | 0.024872 |
| Gut-tube/Endothelium | Bambi | 1.1973 | 1.79E-06 | 0.00012093 |
| Gut-tube/Endothelium | Bmp4 | 0.75782 | 0.0021821 | 0.025667 |
| Gut-tube/Endothelium | Cdh5 | 0.59222 | 0.005276 | 0.047578 |
| Gut-tube/Endothelium | Hoxa13 | 0.59915 | 0.0024044 | 0.027103 |
| Gut-tube/Endothelium | Hoxa9 | 0.84007 | 0.00057805 | 0.009199 |
| Gut-tube/Endothelium | Hoxb5 | 0.65765 | 0.002056 | 0.025283 |
| Gut-tube/Endothelium | Hoxb6 | 0.86201 | 0.00046058 | 0.0095849 |
| Gut-tube/Endothelium | Hoxc6 | 1.1073 | 3.40E-06 | 0.0001839 |
| Gut-tube/Endothelium | Hoxc8 | 0.72995 | 0.0011065 | 0.01663 |
| Gut-tube/Endothelium | Hoxd4 | 1.3517 | 4.00E-07 | 5.41E-05 |
| Gut-tube/Endothelium | Hoxd9 | 0.69354 | 0.0020026 | 0.025799 |
| Gut-tube/Endothelium | Isl1 | 0.83392 | 0.00048117 | 0.0092981 |
| Gut-tube/Endothelium | Nkx2-3 | 0.68946 | 0.0030874 | 0.032124 |
| Gut-tube/Endothelium | Tbx3 | 1.1072 | 2.10E-05 | 0.00070868 |
| Gut-tube/Endothelium | Wnt5a | 0.77817 | 0.0017877 | 0.024181 |

| | | | | |
|---|---|---|---|---|
| Gut-tube/Neural-crest | Ash2l | 1.2737 | 0.00053721 | 0.005899 |
| Gut-tube/Neural-crest | Bid | 1.0025 | 0.00089828 | 0.0080372 |
| Gut-tube/Neural-crest | Dlk1 | 2.4954 | 4.72E-10 | 2.85E-08 |
| Gut-tube/Neural-crest | Dusp6 | 1.9039 | 3.09E-07 | 6.21E-06 |
| Gut-tube/Neural-crest | Efna5 | 1.1249 | 0.0026355 | 0.019896 |
| Gut-tube/Neural-crest | Fli1 | 1.0041 | 0.008586 | 0.047141 |
| Gut-tube/Neural-crest | Fzd2 | 0.68254 | 0.001773 | 0.014769 |
| Gut-tube/Neural-crest | Gpc4 | 1.6702 | 3.42E-06 | 4.86E-05 |
| Gut-tube/Neural-crest | Hemgn | 1.1618 | 0.0037711 | 0.026795 |
| Gut-tube/Neural-crest | Hoxa11 | 0.78699 | 0.0069436 | 0.040913 |
| Gut-tube/Neural-crest | Lef1 | 1.6595 | 3.37E-06 | 5.09E-05 |
| Gut-tube/Neural-crest | Lin28a | 0.75311 | 2.94E-08 | 8.88E-07 |
| Gut-tube/Neural-crest | Meis2 | 1.9177 | 2.66E-06 | 4.28E-05 |
| Gut-tube/Neural-crest | Nkx2-3 | 1.9218 | 3.64E-07 | 6.76E-06 |
| Gut-tube/Neural-crest | Otx2 | 0.91507 | 0.006966 | 0.040068 |
| Gut-tube/Neural-crest | Pcgf2 | 1.0744 | 0.0054936 | 0.034924 |
| Gut-tube/Neural-crest | Pdgfa | 1.1889 | 0.0022013 | 0.017155 |
| Gut-tube/Neural-crest | Perp | 0.86723 | 0.0064773 | 0.03912 |
| Gut-tube/Neural-crest | Pitx1 | 3.5671 | 5.08E-10 | 2.45E-08 |
| Gut-tube/Neural-crest | Postn | 0.80757 | 0.0075977 | 0.042685 |
| Gut-tube/Neural-crest | Shisa2 | 1.3657 | 0.0012908 | 0.011137 |
| Gut-tube/Neural-crest | Sox2 | 2.062 | 1.24E-07 | 2.72E-06 |
| Gut-tube/Neural-crest | Tfap2a | 1.0207 | 0.0061981 | 0.038393 |
| Gut-tube/Splanchnic-mesoderm | Aldh1a2 | 1.0379 | 0.00019299 | 0.0011675 |
| Gut-tube/Splanchnic-mesoderm | Axin2 | 0.86828 | 0.00089664 | 0.0046844 |
| Gut-tube/Splanchnic-mesoderm | Bak1 | 0.7818 | 0.0035756 | 0.015221 |
| Gut-tube/Splanchnic-mesoderm | Fgfr2 | 0.85245 | 0.0021606 | 0.0099332 |
| Gut-tube/Splanchnic-mesoderm | Foxa1 | 0.70705 | 0.00074433 | 0.0040739 |
| Gut-tube/Splanchnic-mesoderm | Foxf1 | 1.9282 | 7.80E-09 | 1.20E-07 |
| Gut-tube/Splanchnic-mesoderm | Fzd2 | 0.62421 | 4.37E-05 | 0.00031361 |
| Gut-tube/Splanchnic-mesoderm | Gata4 | 1.0114 | 0.00056188 | 0.0031503 |
| Gut-tube/Splanchnic-mesoderm | Gata5 | 1.8057 | 3.31E-12 | 8.45E-11 |
| Gut-tube/Splanchnic-mesoderm | Gata6 | 1.2821 | 6.23E-06 | 4.94E-05 |
| Gut-tube/Splanchnic-mesoderm | Hoxa1 | 1.4015 | 4.02E-07 | 4.40E-06 |
| Gut-tube/Splanchnic-mesoderm | Hoxb1 | 1.7462 | 1.48E-07 | 1.70E-06 |
| Gut-tube/Splanchnic-mesoderm | Irx1 | 1.4286 | 5.84E-08 | 7.46E-07 |
| Gut-tube/Splanchnic-mesoderm | Irx3 | 1.1808 | 5.27E-06 | 4.33E-05 |
| Gut-tube/Splanchnic-mesoderm | Jarid2 | 0.90024 | 0.0013686 | 0.0068395 |
| Gut-tube/Splanchnic-mesoderm | Lin28a | 0.8291 | 1.29E-10 | 2.70E-09 |
| Gut-tube/Splanchnic-mesoderm | Meis1 | 1.291 | 9.83E-07 | 1.03E-05 |
| Gut-tube/Splanchnic-mesoderm | Meis2 | 1.3535 | 2.73E-06 | 2.32E-05 |
| Gut-tube/Splanchnic-mesoderm | Msx2 | 0.67719 | 0.0036379 | 0.015205 |
| Gut-tube/Splanchnic-mesoderm | Osr1 | 2.031 | 2.26E-14 | 8.67E-13 |
| Gut-tube/Splanchnic-mesoderm | Pcgf2 | 0.70618 | 0.0098925 | 0.032957 |
| Gut-tube/Splanchnic-mesoderm | Prrx2 | 0.82584 | 0.0021942 | 0.0098898 |
| Gut-tube/Splanchnic-mesoderm | Setd1a | 0.67454 | 0.0033928 | 0.014715 |
| Gut-tube/Splanchnic-mesoderm | Smarcd3 | 0.94052 | 2.74E-05 | 0.00021008 |
| Gut-tube/Splanchnic-mesoderm | Sox4 | 0.98806 | 8.45E-10 | 1.50E-08 |
| Gut-tube/Splanchnic-mesoderm | Tbx5 | 0.67251 | 0.004384 | 0.01768 |
| Lateral-plate-mesoderm/Endothelium | Hapln1 | 1.5986 | 2.96E-07 | 0.00010041 |
| Lateral-plate-mesoderm/Endothelium | Plp1 | 1.2048 | 0.00027248 | 0.030841 |
| Lateral-plate-mesoderm/Intermediate-mesoderm | Alx1 | 0.6047 | 0.00013986 | 0.00043603 |
| Lateral-plate-mesoderm/Intermediate-mesoderm | Apob | 0.53292 | 4.72E-05 | 0.00015801 |
| Lateral-plate-mesoderm/Intermediate-mesoderm | Aqp8 | 0.47292 | 0.00072235 | 0.0019496 |

| | | | | |
|---|---|---|---|---|
| Lateral-plate-mesoderm/Intermediate-mesoderm | Cdx1 | 0.44778 | 0.0083334 | 0.014216 |
| Lateral-plate-mesoderm/Intermediate-mesoderm | Col1a2 | 0.87516 | 5.14E-07 | 2.82E-06 |
| Lateral-plate-mesoderm/Intermediate-mesoderm | Dlk1 | 0.75011 | 4.91E-08 | 3.17E-07 |
| Lateral-plate-mesoderm/Intermediate-mesoderm | En1 | 0.41797 | 0.003309 | 0.006504 |
| Lateral-plate-mesoderm/Intermediate-mesoderm | Eomes | 0.44372 | 0.0011696 | 0.0029373 |
| Lateral-plate-mesoderm/Intermediate-mesoderm | Fezf1 | 0.45527 | 0.001905 | 0.0042009 |
| Lateral-plate-mesoderm/Intermediate-mesoderm | Fgf3 | 0.42246 | 0.0010404 | 0.0026876 |
| Lateral-plate-mesoderm/Intermediate-mesoderm | Fgfr2 | 1.1065 | 8.83E-10 | 7.98E-09 |
| Lateral-plate-mesoderm/Intermediate-mesoderm | Fxyd2 | 0.41054 | 0.0027471 | 0.0056449 |
| Lateral-plate-mesoderm/Intermediate-mesoderm | Gata6 | 0.66682 | 0.0001656 | 0.00049091 |
| Lateral-plate-mesoderm/Intermediate-mesoderm | Gbx2 | 0.46368 | 0.00016441 | 0.00049551 |
| Lateral-plate-mesoderm/Intermediate-mesoderm | Hoxb8 | 0.50613 | 0.0046627 | 0.0086922 |
| Lateral-plate-mesoderm/Intermediate-mesoderm | Hoxb9 | 0.40281 | 0.0041881 | 0.0079719 |
| Lateral-plate-mesoderm/Intermediate-mesoderm | Icam2 | 0.53932 | 0.00056947 | 0.0015603 |
| Lateral-plate-mesoderm/Intermediate-mesoderm | Irx3 | 2.0306 | 4.31E-31 | 7.80E-29 |
| Lateral-plate-mesoderm/Intermediate-mesoderm | Irx5 | 0.95695 | 1.26E-08 | 9.13E-08 |
| Lateral-plate-mesoderm/Intermediate-mesoderm | Lhx1 | 0.66243 | 5.19E-05 | 0.00017078 |
| Lateral-plate-mesoderm/Intermediate-mesoderm | Mesp2 | 0.49488 | 0.0014884 | 0.0035414 |
| Lateral-plate-mesoderm/Intermediate-mesoderm | Msx1 | 1.1663 | 8.25E-15 | 1.66E-13 |
| Lateral-plate-mesoderm/Intermediate-mesoderm | Nebl | 0.56116 | 0.00015796 | 0.00048413 |
| Lateral-plate-mesoderm/Intermediate-mesoderm | Nr1d1 | 0.516 | 0.00022909 | 0.00066817 |
| Lateral-plate-mesoderm/Intermediate-mesoderm | Ovol2 | 0.60489 | 3.07E-05 | 0.00010684 |
| Lateral-plate-mesoderm/Intermediate-mesoderm | Prrx1 | 1.1195 | 1.41E-11 | 1.59E-10 |
| Lateral-plate-mesoderm/Intermediate-mesoderm | Psmb8 | 0.64566 | 5.24E-06 | 2.49E-05 |
| Lateral-plate-mesoderm/Intermediate-mesoderm | Ptn | 0.44399 | 0.0026236 | 0.0054532 |
| Lateral-plate-mesoderm/Intermediate-mesoderm | Sfrp2 | 0.47686 | 0.0064725 | 0.011363 |
| Lateral-plate-mesoderm/Intermediate-mesoderm | Slc7a3 | 0.72275 | 4.89E-06 | 2.39E-05 |
| Lateral-plate-mesoderm/Intermediate-mesoderm | Wnt11 | 0.52035 | 0.00088242 | 0.0023466 |
| Lateral-plate-mesoderm/Intermediate-mesoderm | Wnt2 | 0.99581 | 6.71E-09 | 5.06E-08 |
| Neural-crest/Surface-ectoderm | Cdh1 | 0.75385 | 0.00057645 | 0.036134 |
| Neural-crest/Surface-ectoderm | Cldn4 | 0.92362 | 6.18E-05 | 0.0064533 |
| Neural-crest/Surface-ectoderm | Irx3 | 0.92542 | 0.00044808 | 0.035109 |
| Neural-crest/Surface-ectoderm | Krt18 | 1.0428 | 1.07E-05 | 0.0016718 |
| Presomitic-mesoderm/Dermomyotome | Aldh1a2 | 1.2495 | 1.58E-05 | 0.00098436 |
| Presomitic-mesoderm/Dermomyotome | Cer1 | 1.2468 | 2.25E-05 | 0.0010053 |
| Presomitic-mesoderm/Dermomyotome | Col26a1 | 0.96443 | 0.00038625 | 0.015074 |
| Spinal-cord/Forebrain-Midbrain-Hindbrain | Akr1c19 | 0.51772 | 0.00083698 | 0.00087424 |
| Spinal-cord/Forebrain-Midbrain-Hindbrain | Axin2 | 0.4576 | 0.0090091 | 0.0067996 |
| Spinal-cord/Forebrain-Midbrain-Hindbrain | Cntfr | 0.88263 | 1.15E-07 | 4.81E-07 |
| Spinal-cord/Forebrain-Midbrain-Hindbrain | Col4a1 | 0.60918 | 0.00082659 | 0.00087116 |
| Spinal-cord/Forebrain-Midbrain-Hindbrain | Dll3 | 1.0744 | 7.35E-08 | 3.44E-07 |
| Spinal-cord/Forebrain-Midbrain-Hindbrain | Eed | 0.59995 | 0.00028375 | 0.00036478 |
| Spinal-cord/Forebrain-Midbrain-Hindbrain | Efna5 | 0.5254 | 0.0047742 | 0.0040472 |
| Spinal-cord/Forebrain-Midbrain-Hindbrain | Fgfr3 | 1.3266 | 3.24E-13 | 5.41E-12 |
| Spinal-cord/Forebrain-Midbrain-Hindbrain | Fzd2 | 0.54071 | 0.0020069 | 0.0019244 |
| Spinal-cord/Forebrain-Midbrain-Hindbrain | Hoxb5 | 0.76577 | 2.76E-05 | 5.22E-05 |
| Spinal-cord/Forebrain-Midbrain-Hindbrain | Hoxb6 | 0.62255 | 0.00053743 | 0.0006104 |
| Spinal-cord/Forebrain-Midbrain-Hindbrain | Kmt2b | 0.87373 | 2.06E-06 | 5.88E-06 |
| Spinal-cord/Forebrain-Midbrain-Hindbrain | Lfng | 1.155 | 6.62E-10 | 5.53E-09 |
| Spinal-cord/Forebrain-Midbrain-Hindbrain | Marcks | 1.7571 | 1.14E-20 | 6.69E-19 |
| Spinal-cord/Forebrain-Midbrain-Hindbrain | Meis2 | 0.50508 | 0.0039384 | 0.0034129 |
| Spinal-cord/Forebrain-Midbrain-Hindbrain | Nr2f1 | 1.163 | 2.50E-13 | 4.88E-12 |
| Spinal-cord/Forebrain-Midbrain-Hindbrain | Pcgf2 | 0.88099 | 9.33E-07 | 3.12E-06 |
| Spinal-cord/Forebrain-Midbrain-Hindbrain | Pcgf3 | 0.69758 | 6.94E-05 | 0.00011591 |

| | | | | |
|---|---|---|---|---|
| Spinal-cord/Forebrain-Midbrain-Hindbrain | Setd2 | 0.73483 | 3.89E-05 | 6.89E-05 |
| Spinal-cord/Forebrain-Midbrain-Hindbrain | Sfrp1 | 1.0309 | 1.04E-09 | 8.11E-09 |
| Spinal-cord/Forebrain-Midbrain-Hindbrain | Sfrp2 | 0.74992 | 2.13E-05 | 4.22E-05 |
| Spinal-cord/Forebrain-Midbrain-Hindbrain | Sox4 | 0.85913 | 4.25E-06 | 1.08E-05 |
| Spinal-cord/Forebrain-Midbrain-Hindbrain | Tead2 | 0.79958 | 5.93E-08 | 2.89E-07 |
| Splanchnic-mesoderm/Cardiomyocytes | Aplnr | 1.1703 | 0.0042876 | 0.031551 |
| Splanchnic-mesoderm/Cardiomyocytes | Atp1b1 | 0.93407 | 0.0026506 | 0.021334 |
| Splanchnic-mesoderm/Cardiomyocytes | Bmp4 | 1.8167 | 1.37E-05 | 0.00019667 |
| Splanchnic-mesoderm/Cardiomyocytes | Bmp7 | 1.5152 | 8.81E-05 | 0.0010807 |
| Splanchnic-mesoderm/Cardiomyocytes | Clic6 | 0.95395 | 0.0054279 | 0.037784 |
| Splanchnic-mesoderm/Cardiomyocytes | Cxcl12 | 1.6084 | 1.27E-05 | 0.00019165 |
| Splanchnic-mesoderm/Cardiomyocytes | Dusp6 | 1.5439 | 6.39E-06 | 0.0001097 |
| Splanchnic-mesoderm/Cardiomyocytes | Gata5 | 1.5007 | 1.38E-06 | 3.24E-05 |
| Splanchnic-mesoderm/Cardiomyocytes | Gata6 | 1.5091 | 7.33E-10 | 3.15E-08 |
| Splanchnic-mesoderm/Cardiomyocytes | Hand2 | 2.086 | 2.81E-08 | 8.03E-07 |
| Splanchnic-mesoderm/Cardiomyocytes | Hapln1 | 2.391 | 3.93E-11 | 2.53E-09 |
| Splanchnic-mesoderm/Cardiomyocytes | Irx1 | 1.15 | 0.00067174 | 0.0064078 |
| Splanchnic-mesoderm/Cardiomyocytes | Irx3 | 2.2087 | 5.00E-11 | 2.57E-09 |
| Splanchnic-mesoderm/Cardiomyocytes | Irx5 | 1.2838 | 0.00076907 | 0.0070743 |
| Splanchnic-mesoderm/Cardiomyocytes | Isl1 | 1.2548 | 0.0005414 | 0.0053631 |
| Splanchnic-mesoderm/Cardiomyocytes | Mkrn1 | 0.92771 | 0.0033664 | 0.025501 |
| Splanchnic-mesoderm/Cardiomyocytes | Nebl | 0.88935 | 0.0075167 | 0.049641 |
| Splanchnic-mesoderm/Cardiomyocytes | Pdgfra | 1.4877 | 1.30E-08 | 4.19E-07 |
| Splanchnic-mesoderm/Cardiomyocytes | Per2 | 1.0007 | 0.0025437 | 0.021134 |
| Splanchnic-mesoderm/Cardiomyocytes | Popdc2 | 2.864 | 3.52E-19 | 9.06E-17 |
| Splanchnic-mesoderm/Cardiomyocytes | Prrx2 | 1.1567 | 0.0017097 | 0.014678 |
| Splanchnic-mesoderm/Cardiomyocytes | Smarcd3 | 1.5205 | 4.61E-06 | 8.49E-05 |
| Splanchnic-mesoderm/Cardiomyocytes | Tagln | 1.5105 | 0.00051432 | 0.0055195 |
| Splanchnic-mesoderm/Cardiomyocytes | Tbx1 | 1.4757 | 0.00052965 | 0.0054566 |
| Splanchnic-mesoderm/Cardiomyocytes | Tmem108 | 0.92857 | 0.0013527 | 0.012014 |
| Splanchnic-mesoderm/Cardiomyocytes | Ttn | 2.2451 | 2.73E-06 | 5.86E-05 |
| Splanchnic-mesoderm/Cardiomyocytes | Wnt5a | 1.523 | 9.14E-07 | 2.35E-05 |
| Splanchnic-mesoderm/Gut-tube | Akr1c19 | 1.4649 | 2.84E-09 | 3.32E-07 |
| Splanchnic-mesoderm/Gut-tube | Alas2 | 0.53601 | 0.006075 | 0.037386 |
| Splanchnic-mesoderm/Gut-tube | Aldh1a2 | 0.54555 | 0.00094327 | 0.0095907 |
| Splanchnic-mesoderm/Gut-tube | Aldh2 | 0.97647 | 6.84E-05 | 0.0013331 |
| Splanchnic-mesoderm/Gut-tube | Bid | 0.82746 | 8.94E-05 | 0.001609 |
| Splanchnic-mesoderm/Gut-tube | Cldn4 | 0.57603 | 0.0087777 | 0.047737 |
| Splanchnic-mesoderm/Gut-tube | Clic6 | 0.67343 | 0.0042363 | 0.029137 |
| Splanchnic-mesoderm/Gut-tube | Cxcl12 | 1.3246 | 7.36E-07 | 4.30E-05 |
| Splanchnic-mesoderm/Gut-tube | Dnmt3b | 0.82429 | 0.0023412 | 0.01825 |
| Splanchnic-mesoderm/Gut-tube | Ets1 | 0.92109 | 0.00070804 | 0.0082789 |
| Splanchnic-mesoderm/Gut-tube | Etv4 | 0.5853 | 0.0052455 | 0.034075 |
| Splanchnic-mesoderm/Gut-tube | Ezh1 | 0.79308 | 0.00078315 | 0.0083247 |
| Splanchnic-mesoderm/Gut-tube | Foxa1 | 1.2303 | 3.73E-06 | 0.00010891 |
| Splanchnic-mesoderm/Gut-tube | Foxa2 | 1.07 | 1.14E-05 | 0.00029596 |
| Splanchnic-mesoderm/Gut-tube | Foxh1 | 0.55398 | 0.0056098 | 0.035456 |
| Splanchnic-mesoderm/Gut-tube | Gfi1 | 0.55692 | 0.005034 | 0.033635 |
| Splanchnic-mesoderm/Gut-tube | Gypc | 0.60598 | 0.0074093 | 0.043317 |
| Splanchnic-mesoderm/Gut-tube | Hoxa1 | 1.0585 | 0.00010221 | 0.0017073 |
| Splanchnic-mesoderm/Gut-tube | Irx1 | 0.83414 | 0.00034625 | 0.004763 |
| Splanchnic-mesoderm/Gut-tube | Irx3 | 1.8923 | 1.45E-09 | 3.39E-07 |
| Splanchnic-mesoderm/Gut-tube | Irx5 | 0.81812 | 0.00097533 | 0.0095035 |
| Splanchnic-mesoderm/Gut-tube | Itga3 | 0.68263 | 0.0063017 | 0.037787 |

| | | | | |
|---|---|---|---|---|
| Splanchnic-mesoderm/Gut-tube | Krt18 | 0.88377 | 0.0001439 | 0.0022434 |
| Splanchnic-mesoderm/Gut-tube | Ldhb | 0.61289 | 0.0089947 | 0.047805 |
| Splanchnic-mesoderm/Gut-tube | Myh9 | 0.8777 | 7.90E-08 | 6.16E-06 |
| Splanchnic-mesoderm/Gut-tube | Nepn | 0.59803 | 0.001866 | 0.015585 |
| Splanchnic-mesoderm/Gut-tube | Osr1 | 0.93998 | 2.93E-06 | 9.78E-05 |
| Splanchnic-mesoderm/Gut-tube | Prrx1 | 0.64162 | 0.0020632 | 0.016638 |
| Splanchnic-mesoderm/Gut-tube | Prrx2 | 0.7895 | 0.00077862 | 0.0086706 |
| Splanchnic-mesoderm/Gut-tube | Setd1b | 0.68907 | 0.0034886 | 0.025494 |
| Splanchnic-mesoderm/Gut-tube | Shh | 0.80963 | 0.00028801 | 0.0042095 |
| Splanchnic-mesoderm/Gut-tube | Sox2 | 0.7037 | 0.0016923 | 0.015221 |
| Splanchnic-mesoderm/Gut-tube | Sox4 | 0.66631 | 3.08E-05 | 0.00065489 |
| Splanchnic-mesoderm/Gut-tube | Tbx1 | 1.3232 | 1.07E-06 | 4.18E-05 |
| Splanchnic-mesoderm/Gut-tube | Tbx5 | 0.82469 | 0.0030159 | 0.022751 |
| Splanchnic-mesoderm/Gut-tube | Tcf7l1 | 0.63106 | 0.0039017 | 0.027649 |
| Splanchnic-mesoderm/Gut-tube | Tjp2 | 1.0023 | 3.00E-05 | 0.00070111 |
| Splanchnic-mesoderm/Gut-tube | Wnt3 | 0.65412 | 0.0015275 | 0.014288 |
| Splanchnic-mesoderm/Gut-tube | Wnt5a | 0.79594 | 0.00044463 | 0.0057766 |
| Surface-ectoderm/Mixed-mesenchymal-mesoderm | Ahnak | 1.8915 | 4.97E-06 | 0.00074984 |
| Surface-ectoderm/Mixed-mesenchymal-mesoderm | Col1a1 | 1.5966 | 0.00024154 | 0.014572 |
| Surface-ectoderm/Mixed-mesenchymal-mesoderm | Hand1 | 2.2207 | 3.73E-06 | 0.0011245 |
| Surface-ectoderm/Mixed-mesenchymal-mesoderm | Hand2 | 1.5498 | 0.00012311 | 0.0092841 |
| Surface-ectoderm/Mixed-mesenchymal-mesoderm | Popdc2 | 1.3353 | 0.00041098 | 0.020662 |
| Surface-ectoderm/Mixed-mesenchymal-mesoderm | Prrx2 | 1.021 | 0.0013876 | 0.038053 |
| Surface-ectoderm/Mixed-mesenchymal-mesoderm | Smoc2 | 1.1704 | 0.0020103 | 0.035671 |
| Surface-ectoderm/Mixed-mesenchymal-mesoderm | Tfap2a | 0.93662 | 0.001545 | 0.03585 |
| Surface-ectoderm/Mixed-mesenchymal-mesoderm | Tgm1 | 0.937 | 0.0022175 | 0.035206 |
| Surface-ectoderm/Neural-crest | Nr2f1 | 1.3944 | 1.62E-06 | 0.00022888 |
| Surface-ectoderm/Neural-crest | Prrx1 | 0.8044 | 0.001608 | 0.041304 |
| Surface-ectoderm/Neural-crest | Prrx2 | 0.81931 | 0.0013976 | 0.039489 |
| Surface-ectoderm/Neural-crest | Snai1 | 1.0989 | 6.72E-05 | 0.0047472 |
| Surface-ectoderm/Neural-crest | Tfap2b | 1.0485 | 0.00048653 | 0.019638 |
| * The logFC was calculated based on the natural logarithm. | | | | |

#### Table S1B. Down-regulated genes identified from the mouse embryo seqFISH data

| Centered cell type/neighboring cell type | Down-regulated_gene | Homotypic1_logFC | Homotypic1_p.value | Homotypic1_fdr |
|---|---|---|---|---|
| Cardiomyocytes/Endothelium | Ahnak | -1.0695 | 7.47E-06 | 0.0010773 |
| Cardiomyocytes/Endothelium | Bmp4 | -0.83081 | 0.0013849 | 0.023496 |
| Cardiomyocytes/Endothelium | Dlk1 | -0.87429 | 0.00061071 | 0.014678 |
| Cardiomyocytes/Endothelium | Fzd2 | -0.79785 | 0.00052737 | 0.01521 |
| Cardiomyocytes/Endothelium | Hand2 | -0.73097 | 0.00099594 | 0.017953 |
| Cardiomyocytes/Endothelium | Isl1 | -0.63739 | 0.00040198 | 0.014492 |
| Cardiomyocytes/Endothelium | Itga3 | -0.93565 | 0.00028818 | 0.011874 |
| Cardiomyocytes/Endothelium | Krt18 | -0.99548 | 1.85E-05 | 0.0010672 |
| Cardiomyocytes/Endothelium | Mnt | -0.71648 | 0.001528 | 0.023194 |
| Cardiomyocytes/Endothelium | Shisa2 | -1.1706 | 1.90E-06 | 0.00054695 |
| Cardiomyocytes/Mixed-mesenchymal-mesoderm | Bambi | -0.99209 | 0.00068056 | 0.020091 |
| Cardiomyocytes/Mixed-mesenchymal-mesoderm | Bcl2 | -0.74418 | 0.00049577 | 0.016262 |
| Cardiomyocytes/Mixed-mesenchymal-mesoderm | Bmp7 | -0.96557 | 0.0022576 | 0.047604 |
| Cardiomyocytes/Mixed-mesenchymal-mesoderm | Hsf1 | -0.89094 | 0.00027804 | 0.02052 |
| Cardiomyocytes/Mixed-mesenchymal-mesoderm | Jarid2 | -0.92957 | 0.0009171 | 0.020826 |
| Cardiomyocytes/Mixed-mesenchymal-mesoderm | Kmt2b | -1.004 | 0.00045623 | 0.016836 |
| Cardiomyocytes/Mixed-mesenchymal-mesoderm | Pcdh19 | -0.40719 | 0.00043247 | 0.018238 |
| Cardiomyocytes/Mixed-mesenchymal-mesoderm | Prrx2 | -1.0259 | 0.00028016 | 0.016541 |
| Cardiomyocytes/Mixed-mesenchymal-mesoderm | Sfrp2 | -0.53095 | 0.00080589 | 0.019826 |
| Cardiomyocytes/Mixed-mesenchymal-mesoderm | Tead2 | -0.98597 | 0.00073007 | 0.019593 |
| Cardiomyocytes/Splanchnic-mesoderm | Atp1b1 | -1.5188 | 0.00069136 | 0.010426 |
| Cardiomyocytes/Splanchnic-mesoderm | Fezf1 | -0.63928 | 0.00038215 | 0.0068434 |
| Cardiomyocytes/Splanchnic-mesoderm | Fli1 | -0.43661 | 0.00010671 | 0.0033973 |
| Cardiomyocytes/Splanchnic-mesoderm | Gata3 | -0.45797 | 0.0053699 | 0.041584 |
| Cardiomyocytes/Splanchnic-mesoderm | Gypa | -0.5061 | 9.23E-06 | 0.00037782 |
| Cardiomyocytes/Splanchnic-mesoderm | Hoxb8 | -0.49177 | 0.0039188 | 0.033025 |
| Cardiomyocytes/Splanchnic-mesoderm | Irx2 | -0.90235 | 3.70E-06 | 0.00017658 |
| Cardiomyocytes/Splanchnic-mesoderm | Lhx1 | -0.96623 | 0.00012453 | 0.0032436 |
| Cardiomyocytes/Splanchnic-mesoderm | Lhx2 | -0.58195 | 0.0018583 | 0.021298 |
| Cardiomyocytes/Splanchnic-mesoderm | Wnt11 | -1.0474 | 0.00015905 | 0.0035055 |
| Cardiomyocytes/Splanchnic-mesoderm | Wnt3a | -0.64445 | 0.00077672 | 0.011127 |
| Cranial-mesoderm/Gut-tube | Morc4 | -0.8168 | 0.0004418 | 0.027701 |
| Cranial-mesoderm/Gut-tube | Rhoj | -1.2565 | 0.0012471 | 0.048873 |
| Dermomyotome/Presomitic-mesoderm | Arhgdib | -1.8602 | 0.00058006 | 0.015 |
| Dermomyotome/Presomitic-mesoderm | Col1a2 | -2.8606 | 6.22E-06 | 0.0017691 |
| Dermomyotome/Presomitic-mesoderm | F2r | -1.7568 | 0.0024687 | 0.036959 |
| Dermomyotome/Presomitic-mesoderm | Fst | -1.7676 | 0.0032831 | 0.044469 |
| Dermomyotome/Presomitic-mesoderm | Gdf3 | -2.1301 | 0.00011127 | 0.0079129 |
| Dermomyotome/Presomitic-mesoderm | Hand2 | -1.9869 | 0.0003442 | 0.012238 |
| Dermomyotome/Presomitic-mesoderm | Hcn4 | -1.3843 | 0.0027769 | 0.039493 |
| Dermomyotome/Presomitic-mesoderm | Hoxb5 | -1.656 | 0.0013471 | 0.025545 |
| Dermomyotome/Presomitic-mesoderm | Mpl | -2.1314 | 0.00042808 | 0.012177 |
| Dermomyotome/Presomitic-mesoderm | Usp28 | -2.5872 | 1.82E-05 | 0.002594 |
| Forebrain-Midbrain-Hindbrain/Cranial-mesoderm | Bambi | -0.55099 | 0.0009412 | 0.0066401 |
| Forebrain-Midbrain-Hindbrain/Cranial-mesoderm | Cdh2 | -0.99093 | 0.00022367 | 0.0023012 |
| Forebrain-Midbrain-Hindbrain/Cranial-mesoderm | Cntfr | -1.1514 | 0.00015806 | 0.0017741 |
| Forebrain-Midbrain-Hindbrain/Cranial-mesoderm | Dlk1 | -1.3221 | 2.40E-09 | 8.48E-08 |
| Forebrain-Midbrain-Hindbrain/Cranial-mesoderm | Dll1 | -1.029 | 0.0008322 | 0.0064214 |
| Forebrain-Midbrain-Hindbrain/Cranial-mesoderm | Eomes | -0.62545 | 1.45E-05 | 0.00023944 |
| Forebrain-Midbrain-Hindbrain/Cranial-mesoderm | Fezf1 | -1.171 | 8.64E-07 | 1.64E-05 |
| Forebrain-Midbrain-Hindbrain/Cranial-mesoderm | Fgf15 | -0.84011 | 1.74E-05 | 0.00026809 |
| Forebrain-Midbrain-Hindbrain/Cranial-mesoderm | Fgfr2 | -1.5559 | 9.81E-08 | 2.20E-06 |
| Forebrain-Midbrain-Hindbrain/Cranial-mesoderm | Fzd2 | -0.99987 | 0.00045477 | 0.0040105 |

| | | | | |
|---|---|---|---|---|
| Forebrain-Midbrain-Hindbrain/Cranial-mesoderm | Irx2 | -0.54961 | 0.0041735 | 0.024536 |
| Forebrain-Midbrain-Hindbrain/Cranial-mesoderm | Irx3 | -1.6766 | 2.11E-11 | 1.04E-09 |
| Forebrain-Midbrain-Hindbrain/Cranial-mesoderm | Irx5 | -0.69337 | 0.00019881 | 0.0021343 |
| Forebrain-Midbrain-Hindbrain/Cranial-mesoderm | Kcng1 | -0.63044 | 0.0062044 | 0.034044 |
| Forebrain-Midbrain-Hindbrain/Cranial-mesoderm | Lefty2 | -0.45111 | 3.72E-06 | 6.56E-05 |
| Forebrain-Midbrain-Hindbrain/Cranial-mesoderm | Lhx2 | -0.7492 | 0.0031382 | 0.019372 |
| Forebrain-Midbrain-Hindbrain/Cranial-mesoderm | Myb | -0.568 | 4.03E-05 | 0.00058495 |
| Forebrain-Midbrain-Hindbrain/Cranial-mesoderm | Nid1 | -0.49014 | 0.0015017 | 0.010022 |
| Forebrain-Midbrain-Hindbrain/Cranial-mesoderm | Nr2f1 | -0.99078 | 0.00023502 | 0.002232 |
| Forebrain-Midbrain-Hindbrain/Cranial-mesoderm | Otx2 | -1.2589 | 0.00022836 | 0.0022555 |
| Forebrain-Midbrain-Hindbrain/Cranial-mesoderm | Pou3f1 | -1.0981 | 0.00059092 | 0.0050313 |
| Forebrain-Midbrain-Hindbrain/Cranial-mesoderm | Ptn | -1.1842 | 8.20E-05 | 0.0010128 |
| Forebrain-Midbrain-Hindbrain/Cranial-mesoderm | Sfrp1 | -1.3134 | 7.50E-05 | 0.00097495 |
| Forebrain-Midbrain-Hindbrain/Cranial-mesoderm | Sfrp2 | -2.1252 | 2.91E-12 | 1.79E-10 |
| Forebrain-Midbrain-Hindbrain/Cranial-mesoderm | Thbs1 | -1.9929 | 2.50E-14 | 2.06E-12 |
| Forebrain-Midbrain-Hindbrain/Cranial-mesoderm | Tinagl1 | -0.44774 | 0.00089905 | 0.0065292 |
| Forebrain-Midbrain-Hindbrain/Definitive-endoderm | Afp | -0.40017 | 0.0039118 | 0.016185 |
| Forebrain-Midbrain-Hindbrain/Definitive-endoderm | Cavin3 | -0.46122 | 0.0010644 | 0.0061451 |
| Forebrain-Midbrain-Hindbrain/Definitive-endoderm | Cd8a | -0.40433 | 2.27E-05 | 0.00019462 |
| Forebrain-Midbrain-Hindbrain/Definitive-endoderm | Coro1a | -0.538 | 0.00076884 | 0.0045444 |
| Forebrain-Midbrain-Hindbrain/Definitive-endoderm | Dlk1 | -1.4588 | 1.52E-10 | 2.69E-09 |
| Forebrain-Midbrain-Hindbrain/Definitive-endoderm | Dll1 | -1.052 | 0.00015074 | 0.0010692 |
| Forebrain-Midbrain-Hindbrain/Definitive-endoderm | Eomes | -0.55072 | 0.001891 | 0.0095803 |
| Forebrain-Midbrain-Hindbrain/Definitive-endoderm | Fezf1 | -1.4399 | 5.91E-11 | 1.22E-09 |
| Forebrain-Midbrain-Hindbrain/Definitive-endoderm | Fgf15 | -0.94337 | 7.61E-07 | 7.87E-06 |
| Forebrain-Midbrain-Hindbrain/Definitive-endoderm | Fgfr3 | -1.047 | 0.004339 | 0.017373 |
| Forebrain-Midbrain-Hindbrain/Definitive-endoderm | Hoxa10 | -0.49135 | 1.27E-07 | 1.58E-06 |
| Forebrain-Midbrain-Hindbrain/Definitive-endoderm | Irx5 | -0.56203 | 0.0081825 | 0.02861 |
| Forebrain-Midbrain-Hindbrain/Definitive-endoderm | Kcng1 | -1.0524 | 2.74E-07 | 3.09E-06 |
| Forebrain-Midbrain-Hindbrain/Definitive-endoderm | Lfng | -0.98684 | 0.004627 | 0.017671 |
| Forebrain-Midbrain-Hindbrain/Definitive-endoderm | Lhx2 | -1.1799 | 5.39E-08 | 7.43E-07 |
| Forebrain-Midbrain-Hindbrain/Definitive-endoderm | Meis2 | -0.95912 | 0.0065427 | 0.023886 |
| Forebrain-Midbrain-Hindbrain/Definitive-endoderm | Myb | -0.6177 | 8.60E-05 | 0.00068878 |
| Forebrain-Midbrain-Hindbrain/Definitive-endoderm | Nefl | -0.63237 | 0.0018124 | 0.0093737 |
| Forebrain-Midbrain-Hindbrain/Definitive-endoderm | Nfe2 | -0.47886 | 1.15E-60 | 2.85E-58 |
| Forebrain-Midbrain-Hindbrain/Definitive-endoderm | Nr2f1 | -1.9862 | 2.04E-11 | 4.61E-10 |
| Forebrain-Midbrain-Hindbrain/Definitive-endoderm | Otx2 | -1.3251 | 0.00040787 | 0.0027366 |
| Forebrain-Midbrain-Hindbrain/Definitive-endoderm | Pax6 | -0.77397 | 1.86E-06 | 1.84E-05 |
| Forebrain-Midbrain-Hindbrain/Definitive-endoderm | Perp | -0.46617 | 0.0016021 | 0.0088383 |
| Forebrain-Midbrain-Hindbrain/Definitive-endoderm | Pou3f1 | -1.7135 | 2.25E-09 | 3.72E-08 |
| Forebrain-Midbrain-Hindbrain/Definitive-endoderm | Ptn | -1.1428 | 0.00053099 | 0.0034689 |
| Forebrain-Midbrain-Hindbrain/Definitive-endoderm | Rgl1 | -0.75825 | 0.0019723 | 0.0094156 |
| Forebrain-Midbrain-Hindbrain/Definitive-endoderm | Sfrp1 | -1.8159 | 5.43E-07 | 5.86E-06 |
| Forebrain-Midbrain-Hindbrain/Definitive-endoderm | Sfrp2 | -1.3281 | 0.00025019 | 0.0017252 |
| Forebrain-Midbrain-Hindbrain/Definitive-endoderm | Six3 | -0.85516 | 4.23E-05 | 0.00034965 |
| Forebrain-Midbrain-Hindbrain/Definitive-endoderm | Sord | -0.69914 | 0.00057751 | 0.003676 |
| Forebrain-Midbrain-Hindbrain/Definitive-endoderm | Tfap2c | -0.46237 | 2.41E-56 | 2.99E-54 |
| Forebrain-Midbrain-Hindbrain/Definitive-endoderm | Thbs1 | -1.5663 | 2.12E-07 | 2.51E-06 |
| Forebrain-Midbrain-Hindbrain/Definitive-endoderm | Tinagl1 | -0.46064 | 0.0019011 | 0.0092541 |
| Forebrain-Midbrain-Hindbrain/Neural-crest | Cdh2 | -1.1075 | 0.00042937 | 0.010279 |
| Forebrain-Midbrain-Hindbrain/Neural-crest | En1 | -0.80113 | 0.00090336 | 0.018742 |
| Forebrain-Midbrain-Hindbrain/Neural-crest | Fst | -0.92053 | 4.09E-05 | 0.0018204 |
| Forebrain-Midbrain-Hindbrain/Neural-crest | Hcn4 | -0.53248 | 1.10E-05 | 0.00068322 |
| Forebrain-Midbrain-Hindbrain/Neural-crest | Hes3 | -0.52541 | 0.00023879 | 0.0074313 |

| | | | | |
|---|---|---|---|---|
| Forebrain-Midbrain-Hindbrain/Neural-crest | Hoxa1 | -0.55357 | 2.35E-69 | 7.30E-67 |
| Forebrain-Midbrain-Hindbrain/Neural-crest | Hoxb1 | -0.61646 | 3.65E-05 | 0.0018955 |
| Forebrain-Midbrain-Hindbrain/Neural-crest | Lfng | -1.2345 | 0.00092182 | 0.01793 |
| Forebrain-Midbrain-Hindbrain/Neural-crest | Nr2f1 | -1.4174 | 0.00080206 | 0.017829 |
| Forebrain-Midbrain-Hindbrain/Neural-crest | Rab27b | -0.41509 | 3.40E-52 | 3.53E-50 |
| Forebrain-Midbrain-Hindbrain/Neural-crest | Tbx1 | -0.48515 | 7.98E-59 | 1.24E-56 |
| Forebrain-Midbrain-Hindbrain/Spinal-cord | Bambi | -0.40394 | 0.00012417 | 0.0011078 |
| Forebrain-Midbrain-Hindbrain/Spinal-cord | Cdh2 | -0.42265 | 0.0089626 | 0.030653 |
| Forebrain-Midbrain-Hindbrain/Spinal-cord | Dlk1 | -0.52128 | 0.0017538 | 0.0087776 |
| Forebrain-Midbrain-Hindbrain/Spinal-cord | Dnmt3b | -0.61004 | 1.27E-05 | 0.00017426 |
| Forebrain-Midbrain-Hindbrain/Spinal-cord | Dusp6 | -0.43684 | 0.00094221 | 0.0053708 |
| Forebrain-Midbrain-Hindbrain/Spinal-cord | Eed | -0.59964 | 0.00011006 | 0.0010266 |
| Forebrain-Midbrain-Hindbrain/Spinal-cord | En1 | -0.62914 | 2.63E-07 | 5.40E-06 |
| Forebrain-Midbrain-Hindbrain/Spinal-cord | Fezf1 | -0.99315 | 7.63E-13 | 5.22E-11 |
| Forebrain-Midbrain-Hindbrain/Spinal-cord | Hoxa10 | -0.41384 | 1.34E-11 | 4.57E-10 |
| Forebrain-Midbrain-Hindbrain/Spinal-cord | Icam2 | -0.42147 | 0.00048232 | 0.0035348 |
| Forebrain-Midbrain-Hindbrain/Spinal-cord | Irx2 | -0.42554 | 0.00015942 | 0.0013085 |
| Forebrain-Midbrain-Hindbrain/Spinal-cord | Kcng1 | -0.4572 | 0.00064115 | 0.0043856 |
| Forebrain-Midbrain-Hindbrain/Spinal-cord | Lef1 | -0.51133 | 0.0020793 | 0.0096972 |
| Forebrain-Midbrain-Hindbrain/Spinal-cord | Lhx2 | -1.225 | 9.38E-31 | 1.92E-28 |
| Forebrain-Midbrain-Hindbrain/Spinal-cord | Myh9 | -0.4202 | 0.0095686 | 0.031167 |
| Forebrain-Midbrain-Hindbrain/Spinal-cord | Otx2 | -1.2325 | 8.29E-12 | 3.40E-10 |
| Forebrain-Midbrain-Hindbrain/Spinal-cord | Ovol2 | -0.42616 | 1.90E-05 | 0.0002289 |
| Forebrain-Midbrain-Hindbrain/Spinal-cord | Ptn | -0.98825 | 5.73E-08 | 1.31E-06 |
| Forebrain-Midbrain-Hindbrain/Spinal-cord | Shh | -0.47044 | 0.00065923 | 0.0043638 |
| Forebrain-Midbrain-Hindbrain/Spinal-cord | Shisa2 | -0.69565 | 1.72E-05 | 0.00022002 |
| Forebrain-Midbrain-Hindbrain/Spinal-cord | Six3 | -0.8811 | 6.22E-16 | 6.38E-14 |
| Forebrain-Midbrain-Hindbrain/Spinal-cord | Tet1 | -0.48543 | 3.88E-05 | 0.00044229 |
| Forebrain-Midbrain-Hindbrain/Spinal-cord | Thbs1 | -0.5628 | 0.0011687 | 0.0063112 |
| Forebrain-Midbrain-Hindbrain/Spinal-cord | Wdr5 | -0.44649 | 0.0041649 | 0.017093 |
| Forebrain-Midbrain-Hindbrain/Spinal-cord | Wnt5b | -0.88825 | 5.72E-12 | 2.93E-10 |
| Gut-tube/Cranial-mesoderm | Aldh1a2 | -0.40972 | 0.0074678 | 0.023457 |
| Gut-tube/Cranial-mesoderm | Aqp3 | -0.7773 | 3.40E-07 | 4.65E-06 |
| Gut-tube/Cranial-mesoderm | Bambi | -1.746 | 9.84E-13 | 3.27E-11 |
| Gut-tube/Cranial-mesoderm | Bin2 | -0.46387 | 0.0016477 | 0.0067193 |
| Gut-tube/Cranial-mesoderm | Bmp4 | -1.0382 | 2.23E-05 | 0.00021619 |
| Gut-tube/Cranial-mesoderm | Cdx1 | -0.41928 | 0.005843 | 0.019402 |
| Gut-tube/Cranial-mesoderm | Cdx2 | -1.7635 | 5.81E-33 | 1.35E-30 |
| Gut-tube/Cranial-mesoderm | Cpm | -2.0321 | 1.44E-13 | 6.68E-12 |
| Gut-tube/Cranial-mesoderm | Cpn1 | -1.0938 | 0.00012251 | 0.00079102 |
| Gut-tube/Cranial-mesoderm | Dlk1 | -0.78017 | 0.0012637 | 0.0057596 |
| Gut-tube/Cranial-mesoderm | Dll1 | -0.46839 | 0.0010886 | 0.0050609 |
| Gut-tube/Cranial-mesoderm | Dnmt3l | -0.45144 | 0.00054376 | 0.0030094 |
| Gut-tube/Cranial-mesoderm | Fgfr4 | -0.98427 | 2.31E-06 | 2.56E-05 |
| Gut-tube/Cranial-mesoderm | Folr1 | -0.59521 | 0.00030136 | 0.0017512 |
| Gut-tube/Cranial-mesoderm | Gata2 | -0.54613 | 0.00094171 | 0.0045603 |
| Gut-tube/Cranial-mesoderm | Gata3 | -1.1196 | 6.44E-05 | 0.00051653 |
| Gut-tube/Cranial-mesoderm | Gata4 | -0.62742 | 0.00028839 | 0.0017188 |
| Gut-tube/Cranial-mesoderm | Gata6 | -1.3968 | 2.04E-08 | 3.38E-07 |
| Gut-tube/Cranial-mesoderm | Gjb3 | -0.45293 | 0.00070297 | 0.0036311 |
| Gut-tube/Cranial-mesoderm | Gjb5 | -0.41383 | 0.0099235 | 0.028833 |
| Gut-tube/Cranial-mesoderm | Hcn4 | -0.4698 | 0.00010042 | 0.0007073 |
| Gut-tube/Cranial-mesoderm | Hoxa9 | -1.2489 | 1.47E-12 | 4.28E-11 |
| Gut-tube/Cranial-mesoderm | Hoxb3 | -1.0966 | 4.37E-09 | 8.46E-08 |

| | | | | |
|---|---|---|---|---|
| Gut-tube/Cranial-mesoderm | Hoxb4 | -0.77375 | 8.16E-05 | 0.00063194 |
| Gut-tube/Cranial-mesoderm | Hoxb5 | -0.67492 | 0.0002351 | 0.0014769 |
| Gut-tube/Cranial-mesoderm | Hoxb6 | -0.57046 | 0.0016327 | 0.0067771 |
| Gut-tube/Cranial-mesoderm | Hoxb8 | -0.77128 | 4.18E-05 | 0.00036014 |
| Gut-tube/Cranial-mesoderm | Hoxb9 | -1.4095 | 1.21E-21 | 7.05E-20 |
| Gut-tube/Cranial-mesoderm | Hoxc6 | -0.85053 | 3.07E-05 | 0.00028551 |
| Gut-tube/Cranial-mesoderm | Hoxc8 | -0.87768 | 2.38E-06 | 2.51E-05 |
| Gut-tube/Cranial-mesoderm | Hoxc9 | -0.62469 | 0.00035968 | 0.0020391 |
| Gut-tube/Cranial-mesoderm | Hoxd4 | -1.4254 | 9.80E-12 | 2.28E-10 |
| Gut-tube/Cranial-mesoderm | Hoxd9 | -0.73927 | 0.00023638 | 0.0014459 |
| Gut-tube/Cranial-mesoderm | Icam2 | -0.52276 | 0.002237 | 0.0086661 |
| Gut-tube/Cranial-mesoderm | Isl1 | -0.86814 | 0.0013626 | 0.0058651 |
| Gut-tube/Cranial-mesoderm | Kitl | -0.91827 | 0.00010561 | 0.00072197 |
| Gut-tube/Cranial-mesoderm | Krt18 | -0.50682 | 0.0044378 | 0.015396 |
| Gut-tube/Cranial-mesoderm | Ldhb | -0.77324 | 0.0042861 | 0.015095 |
| Gut-tube/Cranial-mesoderm | Lef1 | -1.1813 | 3.41E-05 | 0.00030503 |
| Gut-tube/Cranial-mesoderm | Mpl | -0.46626 | 0.0077846 | 0.023809 |
| Gut-tube/Cranial-mesoderm | Msx1 | -0.55058 | 0.0045404 | 0.015295 |
| Gut-tube/Cranial-mesoderm | Nanog | -0.42415 | 0.0034189 | 0.012417 |
| Gut-tube/Cranial-mesoderm | Nepn | -0.55834 | 0.0010875 | 0.005159 |
| Gut-tube/Cranial-mesoderm | Nkx2-3 | -0.73326 | 8.30E-05 | 0.00062263 |
| Gut-tube/Cranial-mesoderm | Osr1 | -0.84841 | 0.0037078 | 0.013259 |
| Gut-tube/Cranial-mesoderm | Sox18 | -0.55523 | 1.77E-09 | 3.74E-08 |
| Gut-tube/Cranial-mesoderm | Tbx3 | -1.8478 | 1.13E-21 | 8.77E-20 |
| Gut-tube/Cranial-mesoderm | Tmem37 | -0.4909 | 0.0028894 | 0.010661 |
| Gut-tube/Endothelium | Cdh1 | -0.69905 | 0.0016568 | 0.02359 |
| Gut-tube/Endothelium | Col4a1 | -1.3968 | 4.35E-09 | 1.18E-06 |
| Gut-tube/Endothelium | Fzd2 | -0.81152 | 0.00012998 | 0.0035164 |
| Gut-tube/Endothelium | Irx1 | -0.67937 | 0.0002535 | 0.0057151 |
| Gut-tube/Endothelium | Kcng1 | -0.46857 | 0.00055634 | 0.010034 |
| Gut-tube/Endothelium | Pdgfa | -0.6979 | 0.0035063 | 0.033878 |
| Gut-tube/Endothelium | Smoc2 | -0.78353 | 1.36E-05 | 0.00052414 |
| Gut-tube/Endothelium | Sox2 | -1.138 | 9.02E-07 | 8.14E-05 |
| Gut-tube/Endothelium | Sp5 | -0.66651 | 0.0005679 | 0.0096022 |
| Gut-tube/Endothelium | Tbx1 | -0.63488 | 9.29E-05 | 0.0027928 |
| Gut-tube/Endothelium | Thbs1 | -0.7553 | 1.18E-05 | 0.00053316 |
| Gut-tube/Neural-crest | Afp | -0.9013 | 1.39E-09 | 5.62E-08 |
| Gut-tube/Neural-crest | Aqp3 | -0.8349 | 9.73E-05 | 0.0013059 |
| Gut-tube/Neural-crest | Cdx2 | -1.6924 | 8.53E-13 | 1.03E-10 |
| Gut-tube/Neural-crest | Coro1a | -0.57096 | 7.02E-26 | 1.70E-23 |
| Gut-tube/Neural-crest | Cpm | -1.41 | 0.00085889 | 0.0079804 |
| Gut-tube/Neural-crest | Cpn1 | -1.8951 | 2.26E-06 | 3.90E-05 |
| Gut-tube/Neural-crest | Dll1 | -0.62088 | 0.00011687 | 0.0014117 |
| Gut-tube/Neural-crest | Dnmt3b | -1.2788 | 0.00070707 | 0.0068325 |
| Gut-tube/Neural-crest | Folr1 | -0.63863 | 0.0042002 | 0.028991 |
| Gut-tube/Neural-crest | Foxa1 | -2.2933 | 1.63E-10 | 1.31E-08 |
| Gut-tube/Neural-crest | Foxa2 | -1.1658 | 0.0021052 | 0.016952 |
| Gut-tube/Neural-crest | Gata6 | -1.7276 | 2.98E-08 | 7.99E-07 |
| Gut-tube/Neural-crest | Hoxa9 | -1.3401 | 1.22E-08 | 4.22E-07 |
| Gut-tube/Neural-crest | Hoxb6 | -0.79725 | 0.00010117 | 0.0012863 |
| Gut-tube/Neural-crest | Hoxc6 | -1.0204 | 0.00019805 | 0.0022783 |
| Gut-tube/Neural-crest | Hoxc8 | -0.80673 | 0.0042793 | 0.028716 |
| Gut-tube/Neural-crest | Kmt2d | -1.0252 | 0.0028875 | 0.021138 |
| Gut-tube/Neural-crest | Marcks | -1.3211 | 0.00070189 | 0.007065 |

| | | | | |
|---|---|---|---|---|
| Gut-tube/Neural-crest | Osr1 | -1.415 | 0.00062846 | 0.0066009 |
| Gut-tube/Neural-crest | Smoc2 | -1.2076 | 1.23E-07 | 2.97E-06 |
| Gut-tube/Neural-crest | Tgm1 | -0.43626 | 0.0050481 | 0.03296 |
| Gut-tube/Splanchnic-mesoderm | Aplnr | -0.42409 | 0.0010595 | 0.005412 |
| Gut-tube/Splanchnic-mesoderm | Aqp3 | -0.91554 | 9.22E-12 | 2.12E-10 |
| Gut-tube/Splanchnic-mesoderm | Bmp2 | -0.43982 | 0.0093359 | 0.032031 |
| Gut-tube/Splanchnic-mesoderm | Cdh1 | -1.2555 | 1.02E-06 | 1.02E-05 |
| Gut-tube/Splanchnic-mesoderm | Cdx2 | -1.5934 | 1.98E-18 | 2.28E-16 |
| Gut-tube/Splanchnic-mesoderm | Cdx4 | -0.42432 | 0.0032453 | 0.014346 |
| Gut-tube/Splanchnic-mesoderm | Cxcl12 | -1.7418 | 1.98E-12 | 5.70E-11 |
| Gut-tube/Splanchnic-mesoderm | Dkk1 | -0.63271 | 1.83E-06 | 1.68E-05 |
| Gut-tube/Splanchnic-mesoderm | Dusp6 | -0.88821 | 0.0019609 | 0.0093909 |
| Gut-tube/Splanchnic-mesoderm | Evx1 | -0.46454 | 0.0072643 | 0.026506 |
| Gut-tube/Splanchnic-mesoderm | Furin | -0.69829 | 0.0060911 | 0.022954 |
| Gut-tube/Splanchnic-mesoderm | Gpc4 | -0.781 | 7.79E-05 | 0.00051149 |
| Gut-tube/Splanchnic-mesoderm | Hoxa11 | -0.52273 | 9.41E-05 | 0.00060092 |
| Gut-tube/Splanchnic-mesoderm | Hoxa7 | -0.9113 | 1.71E-08 | 2.32E-07 |
| Gut-tube/Splanchnic-mesoderm | Hoxa9 | -1.3613 | 1.12E-17 | 8.61E-16 |
| Gut-tube/Splanchnic-mesoderm | Hoxb5 | -0.55649 | 0.0042913 | 0.017615 |
| Gut-tube/Splanchnic-mesoderm | Hoxb6 | -0.64145 | 5.55E-05 | 0.00038665 |
| Gut-tube/Splanchnic-mesoderm | Hoxb8 | -1.1189 | 5.42E-17 | 3.11E-15 |
| Gut-tube/Splanchnic-mesoderm | Hoxb9 | -1.3014 | 4.69E-15 | 2.16E-13 |
| Gut-tube/Splanchnic-mesoderm | Hoxc6 | -0.93102 | 1.61E-06 | 1.54E-05 |
| Gut-tube/Splanchnic-mesoderm | Hoxc8 | -1.06 | 7.67E-10 | 1.47E-08 |
| Gut-tube/Splanchnic-mesoderm | Hoxc9 | -0.6731 | 7.28E-05 | 0.00049239 |
| Gut-tube/Splanchnic-mesoderm | Hoxd4 | -1.6208 | 1.60E-19 | 3.69E-17 |
| Gut-tube/Splanchnic-mesoderm | Hoxd9 | -1.112 | 1.40E-13 | 4.61E-12 |
| Gut-tube/Splanchnic-mesoderm | Kitl | -0.94555 | 4.00E-05 | 0.00029663 |
| Gut-tube/Splanchnic-mesoderm | Lypd6b | -0.60483 | 3.75E-09 | 6.16E-08 |
| Gut-tube/Splanchnic-mesoderm | Notch1 | -0.52149 | 0.00015444 | 0.00095948 |
| Gut-tube/Splanchnic-mesoderm | Otx2 | -0.6127 | 0.00081599 | 0.0043622 |
| Gut-tube/Splanchnic-mesoderm | Perp | -0.50209 | 0.0058594 | 0.022829 |
| Gut-tube/Splanchnic-mesoderm | Pitx1 | -0.46566 | 0.0074178 | 0.026643 |
| Gut-tube/Splanchnic-mesoderm | Pou5f1 | -0.44225 | 1.23E-08 | 1.76E-07 |
| Gut-tube/Splanchnic-mesoderm | Sall3 | -0.50144 | 0.0050456 | 0.019997 |
| Gut-tube/Splanchnic-mesoderm | Six3 | -0.43703 | 2.24E-06 | 1.98E-05 |
| Gut-tube/Splanchnic-mesoderm | Tagln | -0.77556 | 0.00040353 | 0.0023785 |
| Gut-tube/Splanchnic-mesoderm | Wnt5a | -1.1831 | 1.31E-07 | 1.59E-06 |
| Gut-tube/Splanchnic-mesoderm | Zfp57 | -0.69974 | 0.0082353 | 0.028683 |
| Lateral-plate-mesoderm/Endothelium | Pou5f1 | -0.52003 | 1.30E-06 | 0.00021991 |
| Lateral-plate-mesoderm/Endothelium | Zic3 | -0.50596 | 0.00046803 | 0.03973 |
| Lateral-plate-mesoderm/Intermediate-mesoderm | Acvr1 | -0.41503 | 0.0058157 | 0.010516 |
| Lateral-plate-mesoderm/Intermediate-mesoderm | Akr1c19 | -0.491 | 0.0032678 | 0.0064935 |
| Lateral-plate-mesoderm/Intermediate-mesoderm | Aldh1a2 | -0.44259 | 0.0060426 | 0.010819 |
| Lateral-plate-mesoderm/Intermediate-mesoderm | Ash2l | -0.49326 | 0.0044811 | 0.0084407 |
| Lateral-plate-mesoderm/Intermediate-mesoderm | Atp1b1 | -0.54333 | 7.43E-05 | 0.00023982 |
| Lateral-plate-mesoderm/Intermediate-mesoderm | Bak1 | -0.97016 | 2.85E-08 | 1.91E-07 |
| Lateral-plate-mesoderm/Intermediate-mesoderm | Bambi | -1.2097 | 2.70E-11 | 2.88E-10 |
| Lateral-plate-mesoderm/Intermediate-mesoderm | Bmp4 | -0.69205 | 2.22E-05 | 8.35E-05 |
| Lateral-plate-mesoderm/Intermediate-mesoderm | Cdh2 | -0.54138 | 0.0017202 | 0.0038883 |
| Lateral-plate-mesoderm/Intermediate-mesoderm | Cdx2 | -1.0515 | 2.16E-09 | 1.78E-08 |
| Lateral-plate-mesoderm/Intermediate-mesoderm | Cdx4 | -0.80518 | 5.63E-06 | 2.61E-05 |
| Lateral-plate-mesoderm/Intermediate-mesoderm | Cers4 | -0.51956 | 0.0010801 | 0.002751 |
| Lateral-plate-mesoderm/Intermediate-mesoderm | Cpm | -0.4991 | 0.0015145 | 0.0035568 |

| | | | | |
|---|---|---|---|---|
| Lateral-plate-mesoderm/Intermediate-mesoderm | Dnmt3a | -0.8755 | 4.14E-07 | 2.42E-06 |
| Lateral-plate-mesoderm/Intermediate-mesoderm | Dusp6 | -2.0687 | 3.87E-27 | 3.50E-25 |
| Lateral-plate-mesoderm/Intermediate-mesoderm | Efna5 | -0.52511 | 0.0021609 | 0.004597 |
| Lateral-plate-mesoderm/Intermediate-mesoderm | Ep300 | -0.70798 | 2.90E-05 | 0.00010288 |
| Lateral-plate-mesoderm/Intermediate-mesoderm | Ets1 | -0.48035 | 0.0016611 | 0.0038021 |
| Lateral-plate-mesoderm/Intermediate-mesoderm | Etv4 | -0.71853 | 5.76E-06 | 2.60E-05 |
| Lateral-plate-mesoderm/Intermediate-mesoderm | Evx1 | -0.7608 | 2.35E-05 | 8.49E-05 |
| Lateral-plate-mesoderm/Intermediate-mesoderm | Fgfr1 | -0.65864 | 4.56E-05 | 0.00015563 |
| Lateral-plate-mesoderm/Intermediate-mesoderm | Foxf1 | -1.9446 | 4.81E-26 | 2.90E-24 |
| Lateral-plate-mesoderm/Intermediate-mesoderm | Fzd2 | -0.60973 | 0.00011626 | 0.00036884 |
| Lateral-plate-mesoderm/Intermediate-mesoderm | Gata3 | -1.1444 | 7.35E-12 | 9.50E-11 |
| Lateral-plate-mesoderm/Intermediate-mesoderm | Hand2 | -0.44812 | 0.0019365 | 0.0042189 |
| Lateral-plate-mesoderm/Intermediate-mesoderm | Hoxa10 | -1.2514 | 1.53E-13 | 2.51E-12 |
| Lateral-plate-mesoderm/Intermediate-mesoderm | Hoxa11 | -1.4741 | 3.33E-17 | 1.00E-15 |
| Lateral-plate-mesoderm/Intermediate-mesoderm | Hoxa13 | -0.93567 | 2.39E-10 | 2.27E-09 |
| Lateral-plate-mesoderm/Intermediate-mesoderm | Hoxa9 | -0.55157 | 0.0015606 | 0.003618 |
| Lateral-plate-mesoderm/Intermediate-mesoderm | Hoxb6 | -0.79546 | 2.59E-06 | 1.34E-05 |
| Lateral-plate-mesoderm/Intermediate-mesoderm | Hoxc6 | -0.81328 | 4.19E-06 | 2.10E-05 |
| Lateral-plate-mesoderm/Intermediate-mesoderm | Hoxd4 | -0.51261 | 0.00090583 | 0.0023739 |
| Lateral-plate-mesoderm/Intermediate-mesoderm | Hoxd9 | -0.5002 | 0.0039938 | 0.0076829 |
| Lateral-plate-mesoderm/Intermediate-mesoderm | Hsf1 | -0.53519 | 0.0021454 | 0.0046184 |
| Lateral-plate-mesoderm/Intermediate-mesoderm | Igfbp3 | -0.79917 | 5.93E-07 | 3.16E-06 |
| Lateral-plate-mesoderm/Intermediate-mesoderm | Isl1 | -1.5379 | 1.84E-17 | 8.30E-16 |
| Lateral-plate-mesoderm/Intermediate-mesoderm | Itga3 | -0.52437 | 0.0012039 | 0.0029823 |
| Lateral-plate-mesoderm/Intermediate-mesoderm | Jarid2 | -0.77696 | 8.49E-06 | 3.41E-05 |
| Lateral-plate-mesoderm/Intermediate-mesoderm | Krt18 | -0.80651 | 5.94E-06 | 2.62E-05 |
| Lateral-plate-mesoderm/Intermediate-mesoderm | Ldhb | -0.71365 | 7.44E-06 | 3.06E-05 |
| Lateral-plate-mesoderm/Intermediate-mesoderm | Lin28a | -0.67512 | 1.73E-07 | 1.08E-06 |
| Lateral-plate-mesoderm/Intermediate-mesoderm | Lmo2 | -0.48776 | 0.00043732 | 0.0012166 |
| Lateral-plate-mesoderm/Intermediate-mesoderm | Marcks | -0.8053 | 7.41E-06 | 3.19E-05 |
| Lateral-plate-mesoderm/Intermediate-mesoderm | Mcl1 | -0.75401 | 1.54E-05 | 6.05E-05 |
| Lateral-plate-mesoderm/Intermediate-mesoderm | Meis2 | -1.2285 | 1.06E-11 | 1.28E-10 |
| Lateral-plate-mesoderm/Intermediate-mesoderm | Myh9 | -1.2694 | 5.60E-14 | 1.01E-12 |
| Lateral-plate-mesoderm/Intermediate-mesoderm | Nid1 | -0.69136 | 2.19E-05 | 8.42E-05 |
| Lateral-plate-mesoderm/Intermediate-mesoderm | Nkx2-3 | -0.74904 | 7.41E-06 | 3.12E-05 |
| Lateral-plate-mesoderm/Intermediate-mesoderm | Osr1 | -0.91976 | 1.81E-07 | 1.09E-06 |
| Lateral-plate-mesoderm/Intermediate-mesoderm | Pcgf3 | -0.53449 | 0.0017811 | 0.0039761 |
| Lateral-plate-mesoderm/Intermediate-mesoderm | Pdgfra | -1.4901 | 1.53E-15 | 3.46E-14 |
| Lateral-plate-mesoderm/Intermediate-mesoderm | Plvap | -1.4767 | 5.42E-17 | 1.40E-15 |
| Lateral-plate-mesoderm/Intermediate-mesoderm | Prdm1 | -0.99514 | 1.42E-10 | 1.43E-09 |
| Lateral-plate-mesoderm/Intermediate-mesoderm | Setd1a | -0.99597 | 1.75E-08 | 1.22E-07 |
| Lateral-plate-mesoderm/Intermediate-mesoderm | Snai1 | -0.57942 | 0.0014529 | 0.0035503 |
| Lateral-plate-mesoderm/Intermediate-mesoderm | Sox4 | -1.3165 | 3.78E-13 | 5.69E-12 |
| Lateral-plate-mesoderm/Intermediate-mesoderm | Spry4 | -0.55229 | 0.0003993 | 0.0011282 |
| Lateral-plate-mesoderm/Intermediate-mesoderm | Tbx3 | -1.1138 | 1.07E-09 | 9.23E-09 |
| Lateral-plate-mesoderm/Intermediate-mesoderm | Tbx4 | -1.0749 | 4.57E-09 | 3.59E-08 |
| Lateral-plate-mesoderm/Intermediate-mesoderm | Tead2 | -1.5378 | 2.41E-17 | 8.70E-16 |
| Lateral-plate-mesoderm/Intermediate-mesoderm | Tmem119 | -0.76802 | 2.34E-05 | 8.64E-05 |
| Lateral-plate-mesoderm/Intermediate-mesoderm | Twist2 | -0.75589 | 5.12E-07 | 2.90E-06 |
| Lateral-plate-mesoderm/Intermediate-mesoderm | Wdr5 | -0.63523 | 0.0003532 | 0.0010138 |
| Lateral-plate-mesoderm/Intermediate-mesoderm | Wnt5a | -1.2182 | 3.66E-12 | 5.09E-11 |
| Lateral-plate-mesoderm/Intermediate-mesoderm | Wnt5b | -0.51533 | 0.0029182 | 0.0059291 |
| Lateral-plate-mesoderm/Intermediate-mesoderm | Zfp57 | -0.55717 | 0.0014564 | 0.0035115 |
| Neural-crest/Surface-ectoderm | Hand2 | -1.2871 | 8.03E-06 | 0.0025178 |

| | | | | |
|---|---|---|---|---|
| Presomitic-mesoderm/Dermomyotome | Dll1 | -1.5313 | 3.49E-07 | 5.44E-05 |
| Presomitic-mesoderm/Dermomyotome | Dll3 | -1.0695 | 2.19E-05 | 0.0011392 |
| Presomitic-mesoderm/Dermomyotome | Hes7 | -1.3536 | 4.41E-07 | 4.59E-05 |
| Presomitic-mesoderm/Dermomyotome | Mesp2 | -1.9676 | 1.40E-10 | 4.38E-08 |
| Presomitic-mesoderm/Dermomyotome | T | -1.2071 | 6.20E-07 | 4.84E-05 |
| Spinal-cord/Forebrain-Midbrain-Hindbrain | Abcc4 | -0.48646 | 0.00018485 | 0.00025744 |
| Spinal-cord/Forebrain-Midbrain-Hindbrain | Acvr1 | -0.88134 | 3.77E-13 | 5.51E-12 |
| Spinal-cord/Forebrain-Midbrain-Hindbrain | Aldh1a2 | -0.40829 | 0.00145 | 0.0014019 |
| Spinal-cord/Forebrain-Midbrain-Hindbrain | Atp1b1 | -0.66653 | 2.77E-08 | 1.54E-07 |
| Spinal-cord/Forebrain-Midbrain-Hindbrain | Bcl2 | -0.43808 | 0.00032424 | 0.00040786 |
| Spinal-cord/Forebrain-Midbrain-Hindbrain | Bin2 | -0.65697 | 1.01E-10 | 1.31E-09 |
| Spinal-cord/Forebrain-Midbrain-Hindbrain | Ccng1 | -0.72096 | 1.12E-10 | 1.31E-09 |
| Spinal-cord/Forebrain-Midbrain-Hindbrain | Cdh1 | -0.59018 | 1.25E-09 | 8.58E-09 |
| Spinal-cord/Forebrain-Midbrain-Hindbrain | Cdx1 | -0.75966 | 1.06E-15 | 2.47E-14 |
| Spinal-cord/Forebrain-Midbrain-Hindbrain | Clic6 | -0.4464 | 6.20E-06 | 1.42E-05 |
| Spinal-cord/Forebrain-Midbrain-Hindbrain | Cxcl12 | -0.47667 | 1.08E-05 | 2.29E-05 |
| Spinal-cord/Forebrain-Midbrain-Hindbrain | Dnmt3b | -1.0074 | 2.66E-10 | 2.59E-09 |
| Spinal-cord/Forebrain-Midbrain-Hindbrain | Dusp6 | -0.80255 | 3.55E-07 | 1.34E-06 |
| Spinal-cord/Forebrain-Midbrain-Hindbrain | Elf5 | -0.53097 | 1.61E-07 | 6.49E-07 |
| Spinal-cord/Forebrain-Midbrain-Hindbrain | Emcn | -0.67366 | 2.41E-09 | 1.57E-08 |
| Spinal-cord/Forebrain-Midbrain-Hindbrain | En1 | -0.7284 | 8.59E-08 | 3.86E-07 |
| Spinal-cord/Forebrain-Midbrain-Hindbrain | Eng | -0.4457 | 1.85E-06 | 5.40E-06 |
| Spinal-cord/Forebrain-Midbrain-Hindbrain | Epor | -0.40445 | 0.00035557 | 0.00043786 |
| Spinal-cord/Forebrain-Midbrain-Hindbrain | Erg | -0.40363 | 4.43E-05 | 7.62E-05 |
| Spinal-cord/Forebrain-Midbrain-Hindbrain | Esam | -0.45957 | 1.37E-05 | 2.82E-05 |
| Spinal-cord/Forebrain-Midbrain-Hindbrain | Fgf10 | -0.44037 | 0.00010738 | 0.00016104 |
| Spinal-cord/Forebrain-Midbrain-Hindbrain | Fgf17 | -0.48919 | 3.53E-05 | 6.35E-05 |
| Spinal-cord/Forebrain-Midbrain-Hindbrain | Fgf3 | -0.43519 | 0.00015667 | 0.00022627 |
| Spinal-cord/Forebrain-Midbrain-Hindbrain | Fgf5 | -0.4085 | 1.70E-05 | 3.42E-05 |
| Spinal-cord/Forebrain-Midbrain-Hindbrain | Foxh1 | -0.51922 | 9.60E-06 | 2.08E-05 |
| Spinal-cord/Forebrain-Midbrain-Hindbrain | Gata4 | -0.67085 | 2.27E-08 | 1.33E-07 |
| Spinal-cord/Forebrain-Midbrain-Hindbrain | Gata6 | -0.40758 | 8.01E-05 | 0.0001284 |
| Spinal-cord/Forebrain-Midbrain-Hindbrain | Gfi1b | -0.64179 | 5.33E-10 | 4.80E-09 |
| Spinal-cord/Forebrain-Midbrain-Hindbrain | Gjb5 | -0.45312 | 9.69E-05 | 0.00014919 |
| Spinal-cord/Forebrain-Midbrain-Hindbrain | Gng3 | -0.45218 | 6.57E-05 | 0.00011139 |
| Spinal-cord/Forebrain-Midbrain-Hindbrain | Gpc4 | -0.61186 | 6.02E-07 | 2.13E-06 |
| Spinal-cord/Forebrain-Midbrain-Hindbrain | Gypa | -0.45464 | 0.00017397 | 0.0002452 |
| Spinal-cord/Forebrain-Midbrain-Hindbrain | Hemgn | -0.59989 | 4.98E-06 | 1.19E-05 |
| Spinal-cord/Forebrain-Midbrain-Hindbrain | Hes3 | -0.73724 | 5.68E-08 | 2.89E-07 |
| Spinal-cord/Forebrain-Midbrain-Hindbrain | Hes7 | -0.41779 | 0.00047883 | 0.00056015 |
| Spinal-cord/Forebrain-Midbrain-Hindbrain | Hoxa10 | -0.40273 | 3.72E-07 | 1.36E-06 |
| Spinal-cord/Forebrain-Midbrain-Hindbrain | Hoxb1 | -0.86328 | 1.84E-07 | 7.17E-07 |
| Spinal-cord/Forebrain-Midbrain-Hindbrain | Hoxb9 | -2.8149 | 1.00E-38 | 1.17E-36 |
| Spinal-cord/Forebrain-Midbrain-Hindbrain | Hoxc6 | -1.162 | 2.13E-17 | 8.31E-16 |
| Spinal-cord/Forebrain-Midbrain-Hindbrain | Hoxc8 | -1.1239 | 1.72E-16 | 5.02E-15 |
| Spinal-cord/Forebrain-Midbrain-Hindbrain | Hoxc9 | -0.61835 | 5.77E-06 | 1.35E-05 |
| Spinal-cord/Forebrain-Midbrain-Hindbrain | Igfbp3 | -0.50822 | 9.49E-05 | 0.00014807 |
| Spinal-cord/Forebrain-Midbrain-Hindbrain | Ikzf1 | -0.42181 | 0.00045744 | 0.00054054 |
| Spinal-cord/Forebrain-Midbrain-Hindbrain | Itga3 | -0.41872 | 0.00019162 | 0.00026373 |
| Spinal-cord/Forebrain-Midbrain-Hindbrain | Kcng1 | -0.55211 | 0.00020061 | 0.00027289 |
| Spinal-cord/Forebrain-Midbrain-Hindbrain | Kdr | -0.43531 | 8.75E-06 | 1.93E-05 |
| Spinal-cord/Forebrain-Midbrain-Hindbrain | Kitl | -0.51453 | 0.00033066 | 0.00041151 |
| Spinal-cord/Forebrain-Midbrain-Hindbrain | Lef1 | -0.8371 | 3.92E-06 | 1.02E-05 |
| Spinal-cord/Forebrain-Midbrain-Hindbrain | Lypd6b | -0.40606 | 9.90E-05 | 0.00015042 |

| | | | | |
|---|---|---|---|---|
| Spinal-cord/Forebrain-Midbrain-Hindbrain | Mesp2 | -0.55087 | 2.31E-06 | 6.44E-06 |
| Spinal-cord/Forebrain-Midbrain-Hindbrain | Msx1 | -0.57469 | 2.17E-10 | 2.31E-09 |
| Spinal-cord/Forebrain-Midbrain-Hindbrain | Nebl | -0.43258 | 0.00023032 | 0.00029938 |
| Spinal-cord/Forebrain-Midbrain-Hindbrain | Nfe2 | -0.42342 | 2.27E-05 | 4.43E-05 |
| Spinal-cord/Forebrain-Midbrain-Hindbrain | Nrp1 | -0.60121 | 2.61E-05 | 5.00E-05 |
| Spinal-cord/Forebrain-Midbrain-Hindbrain | Osr1 | -0.48291 | 2.78E-06 | 7.57E-06 |
| Spinal-cord/Forebrain-Midbrain-Hindbrain | Pcdh19 | -0.83865 | 1.05E-09 | 7.71E-09 |
| Spinal-cord/Forebrain-Midbrain-Hindbrain | Perp | -0.56364 | 1.56E-06 | 4.81E-06 |
| Spinal-cord/Forebrain-Midbrain-Hindbrain | Piezo2 | -0.65396 | 1.12E-07 | 4.85E-07 |
| Spinal-cord/Forebrain-Midbrain-Hindbrain | Pitx1 | -0.45216 | 0.000359 | 0.00043748 |
| Spinal-cord/Forebrain-Midbrain-Hindbrain | Plac1 | -0.44281 | 3.08E-06 | 8.20E-06 |
| Spinal-cord/Forebrain-Midbrain-Hindbrain | Plp1 | -0.73081 | 1.59E-06 | 4.76E-06 |
| Spinal-cord/Forebrain-Midbrain-Hindbrain | Rgl1 | -0.48732 | 0.0008451 | 0.0008749 |
| Spinal-cord/Forebrain-Midbrain-Hindbrain | Rspo3 | -0.42561 | 3.47E-05 | 6.34E-05 |
| Spinal-cord/Forebrain-Midbrain-Hindbrain | Runx1 | -0.50925 | 1.22E-06 | 3.96E-06 |
| Spinal-cord/Forebrain-Midbrain-Hindbrain | Smim1 | -0.70005 | 4.04E-09 | 2.48E-08 |
| Spinal-cord/Forebrain-Midbrain-Hindbrain | Snai1 | -0.61937 | 6.58E-07 | 2.26E-06 |
| Spinal-cord/Forebrain-Midbrain-Hindbrain | Sox18 | -0.41568 | 0.00030048 | 0.00038209 |
| Spinal-cord/Forebrain-Midbrain-Hindbrain | Tagln | -0.51884 | 1.53E-06 | 4.83E-06 |
| Spinal-cord/Forebrain-Midbrain-Hindbrain | Vamp5 | -0.46035 | 4.52E-06 | 1.10E-05 |
| Spinal-cord/Forebrain-Midbrain-Hindbrain | Vcam1 | -0.537 | 1.15E-05 | 2.40E-05 |
| Spinal-cord/Forebrain-Midbrain-Hindbrain | Wnt3a | -0.53355 | 4.93E-08 | 2.62E-07 |
| Splanchnic-mesoderm/Cardiomyocytes | Aldh1a2 | -1.5433 | 1.40E-05 | 0.00018912 |
| Splanchnic-mesoderm/Cardiomyocytes | Dlk1 | -1.4783 | 0.00045937 | 0.005144 |
| Splanchnic-mesoderm/Cardiomyocytes | Fgfr2 | -1.0741 | 0.0051279 | 0.036687 |
| Splanchnic-mesoderm/Cardiomyocytes | Hoxb1 | -2.174 | 1.29E-09 | 4.74E-08 |
| Splanchnic-mesoderm/Cardiomyocytes | Hoxb4 | -1.6024 | 1.26E-05 | 0.00020229 |
| Splanchnic-mesoderm/Cardiomyocytes | Hoxb5 | -1.2972 | 4.20E-06 | 8.31E-05 |
| Splanchnic-mesoderm/Cardiomyocytes | Kcng1 | -1.5944 | 8.40E-05 | 0.0010816 |
| Splanchnic-mesoderm/Cardiomyocytes | Nr2f1 | -1.1317 | 0.000412 | 0.0048234 |
| Splanchnic-mesoderm/Cardiomyocytes | Zic3 | -0.46373 | 4.48E-15 | 5.77E-13 |
| Splanchnic-mesoderm/Gut-tube | Cdh2 | -0.69181 | 0.0080602 | 0.044879 |
| Splanchnic-mesoderm/Gut-tube | Fgfr2 | -1.0504 | 9.42E-07 | 4.40E-05 |
| Splanchnic-mesoderm/Gut-tube | Gata4 | -0.93962 | 0.0018117 | 0.015692 |
| Splanchnic-mesoderm/Gut-tube | Sfrp1 | -0.66519 | 0.0075646 | 0.043147 |
| Surface-ectoderm/Mixed-mesenchymal-mesoderm | Dusp6 | -1.2952 | 0.00089254 | 0.033655 |
| Surface-ectoderm/Mixed-mesenchymal-mesoderm | F2rl2 | -0.65639 | 8.97E-05 | 0.0090214 |
| Surface-ectoderm/Mixed-mesenchymal-mesoderm | Gfi1 | -0.71279 | 0.0018935 | 0.035698 |
| Surface-ectoderm/Mixed-mesenchymal-mesoderm | Hoxb5 | -0.60954 | 0.0015593 | 0.033597 |
| Surface-ectoderm/Mixed-mesenchymal-mesoderm | Hsf1 | -1.1543 | 0.0011002 | 0.033189 |
| Surface-ectoderm/Mixed-mesenchymal-mesoderm | Osr1 | -0.70426 | 0.00076088 | 0.032788 |
| Surface-ectoderm/Mixed-mesenchymal-mesoderm | Pitx1 | -1.57 | 0.00097836 | 0.032792 |
| Surface-ectoderm/Mixed-mesenchymal-mesoderm | Postn | -1.1946 | 0.0018052 | 0.036303 |
| Surface-ectoderm/Mixed-mesenchymal-mesoderm | Pou3f1 | -0.76394 | 0.0014248 | 0.035817 |
| Surface-ectoderm/Mixed-mesenchymal-mesoderm | Xist | -1.267 | 0.0021306 | 0.035705 |
| Surface-ectoderm/Neural-crest | Acvr2a | -0.96273 | 0.00012006 | 0.0067847 |
| Surface-ectoderm/Neural-crest | Cdh2 | -0.86815 | 0.0017106 | 0.040276 |
| Surface-ectoderm/Neural-crest | Cxcl12 | -1.0668 | 0.00024651 | 0.011608 |
| Surface-ectoderm/Neural-crest | Dlk1 | -1.3499 | 3.51E-05 | 0.0033059 |
| Surface-ectoderm/Neural-crest | Fgf17 | -1.217 | 1.60E-06 | 0.00045343 |
| Surface-ectoderm/Neural-crest | Pitx1 | -1.1876 | 0.00061536 | 0.021734 |
| Surface-ectoderm/Neural-crest | Six3 | -1.0509 | 0.00063497 | 0.019934 |
| * The logFC was calculated based on the natural logarithm. | | | | |

### Table S1C. Up-regulated genes identified from the mouse embryo Slide-seq data

| Heterotypic pair | Up-regulated_gene | Expression in scRNA-seq | Homotypic1_logFC | Hmotypic1_p.value | Homotypic2_logFC | Homotypic2_p.value | Artificial_logFC | Artificial_fdr |
|---|---|---|---|---|---|---|---|---|
| CO+Sc | Sulf1 | CO | 0.53617 | 0.0091785 | 0.42489 | 0.0040106 | 0.46775 | 0.0026889 |
| CTP+Sc | Otor | Sc | 0.61525 | 0.00024382 | 0.43413 | 0.00018499 | 0.49472 | 4.27E-05 |
| DEL+PEL | Hba-a1 | PEL | 0.45666 | 0.0037843 | 1.1599 | 7.54E-22 | 7.94E-01 | 3.10E-15 |
| DEL+PEL | Hbb-bs | PEL | 0.50477 | 0.008049 | 0.99815 | 9.78E-27 | 7.42E-01 | 6.35E-26 |
| DEL+PEL | Hbb-bt | PEL | 1.2001 | 0.0015925 | 0.45918 | 7.01E-05 | 8.72E-01 | 2.89E-16 |
| En+L | Bfsp1 | En | 1.4098 | 2.02E-17 | 1.05E+00 | 1.19E-05 | 1.19E+00 | 1.31E-13 |
| En+L | Cd24a | En | 0.47046 | 1.35E-05 | 4.40E-01 | 0.00025139 | 0.45119 | 9.24E-05 |
| En+L | Crim1 | L | 0.95448 | 9.04E-13 | 7.74E-01 | 2.07E-06 | 8.33E-01 | 1.38E-09 |
| En+L | Cryab | En | 0.83046 | 1.75E-10 | 6.20E-01 | 0.0003532 | 0.69808 | 3.09E-07 |
| En+L | Crygs | n/a | 1.0205 | 2.40E-11 | 8.02E-01 | 6.85E-05 | 8.80E-01 | 5.01E-09 |
| En+L | Dkk3 | En | 0.50235 | 5.09E-06 | 5.43E-01 | 2.09E-07 | 5.25E-01 | 3.40E-06 |
| En+L | Dst | En | 0.60235 | 1.78E-08 | 5.39E-01 | 9.13E-06 | 5.60E-01 | 9.74E-07 |
| En+L | Ednrb | En | 0.66736 | 9.45E-07 | 5.44E-01 | 0.0016906 | 0.59494 | 4.17E-06 |
| En+L | Etl4 | L | 0.49304 | 5.76E-05 | 4.49E-01 | 0.0023909 | 0.46584 | 0.00014904 |
| En+L | Nhs | L | 0.49303 | 7.14E-07 | 4.30E-01 | 0.00021093 | 0.45276 | 3.25E-05 |
| En+L | Pla2g16 | En | 0.49065 | 4.01E-05 | 4.38E-01 | 0.0016948 | 0.45923 | 0.00014759 |
| En+L | Prox1 | En | 0.66281 | 9.49E-08 | 4.75E-01 | 0.0049179 | 0.53676 | 1.43E-05 |
| En+L | Rbm24 | En | 0.63996 | 1.86E-07 | 6.74E-01 | 1.88E-08 | 6.59E-01 | 2.62E-07 |
| En+L | Sfrp1 | L | 0.47534 | 0.00051307 | 0.5258 | 0.00032101 | 0.51026 | 3.83E-05 |
| En+L | Slc2a1 | En | 1.2617 | 1.68E-12 | 7.51E-01 | 0.0024363 | 0.95308 | 5.55E-10 |
| En+L | Tmem40 | En | 0.63456 | 9.64E-06 | 5.37E-01 | 0.0031956 | 0.5749 | 1.48E-05 |
| En+L | Vit | En | 1.3282 | 7.48E-16 | 8.94E-01 | 0.00023405 | 1.0661 | 2.16E-11 |
| Ept+L | Col14a1 | Ept | 0.45834 | 0.0031017 | 0.53229 | 0.00066133 | 0.477 | 0.0064847 |
| L+PEL | Crygs | n/a | 0.5126 | 0.00041996 | 0.57592 | 1.71E-05 | 5.44E-01 | 0.00020594 |
| LM+PEL | Cdkn1c | PEL | 0.4502 | 0.00069251 | 0.51514 | 4.09E-08 | 4.67E-01 | 4.93E-25 |
| Ml+L | Col9a1 | Ml | 0.47361 | 0.00277 | 0.54563 | 0.00081676 | 0.50396 | 0.0049674 |
| Ml+L | Vax2 | Ml | 0.6437 | 0.0014546 | 1.0571 | 1.35E-07 | 8.20E-01 | 7.42E-05 |
| My+PEL | Col3a1 | My | 0.41583 | 4.79E-08 | 1.34E+00 | 1.56E-46 | 7.24E-01 | 2.25E-25 |

*n/a: It is not included in the gene list of scRNA-seq.
* The full names of heterotypic pairs can be found in Table S5.
* The logFC was calculated based on the natural logarithm.

### Table S1D. Down-regulated genes identified from the mouse embryo Slide-seq data

| Heterotypic pair | Down-regulated_gene | Homotypic1_logFC | Hmotypic1_p.value | Homotypic2_logFC | Homotypic2_p.value | Artificial_logFC | Artificial_fdr |
|---|---|---|---|---|---|---|---|
| CO+Sc | Hba-a1 | -0.98848 | 0.0016881 | -0.51959 | 5.80E-20 | -0.69947 | 3.01E-27 |
| CO+Sc | Hba-x | -1.0419 | 0.0029211 | -0.79957 | 2.33E-08 | -0.88447 | 2.49E-11 |
| CO+Sc | Hbb-bs | -0.88471 | 0.0068785 | -0.68574 | 4.16E-09 | -0.76782 | 5.90E-10 |
| CO+Sc | Hbb-y | -1.3501 | 0.00041326 | -1.1259 | 2.46E-19 | -1.2258 | 1.53E-20 |
| CO+Sc | Crybb3 | -2.4289 | 4.60E-11 | -1.8425 | 3.38E-30 | -2.0741 | 6.7089e-2010 |
| CTP+En | Hbb-bt | -1.2299 | 0.0010564 | -1.0157 | 2.27E-05 | -1.1638 | 2.46E-10 |
| CTP+En | Hbb-y | -1.2441 | 0.00035368 | -1.0457 | 0.0011807 | -1.1802 | 0.00026305 |
| CTP+Sc | Hba-x | -1.2581 | 0.00051367 | -0.71971 | 2.52E-15 | -0.93843 | 7.48E-26 |
| CTP+Sc | Hbb-bt | -1.2233 | 0.0011426 | -1.0779 | 1.07E-34 | -1.1489 | 9.43E-37 |
| CTP+Sc | Hbb-y | -2.1357 | 2.40E-12 | -1.3519 | 8.95E-20 | -1.6748 | 4.46E-26 |
| En+LM | Hba-x | -1.1181 | 0.0010388 | -0.74801 | 0.0089333 | -0.83917 | 0.00033351 |
| En+LM | Hbb-bs | -0.95879 | 0.0066835 | -0.84398 | 0.0074749 | -0.8758 | 0.00083342 |
| En+LM | Hbb-y | -1.4151 | 4.01E-05 | -1.3194 | 0.00040162 | -1.3428 | 0.00082487 |
| Ept+Sc | Hba-a1 | -0.65968 | 1.21E-12 | -0.58502 | 1.78E-14 | -0.63054 | 1.63E-20 |
| Ept+Sc | Hba-x | -0.82914 | 7.06E-15 | -0.58164 | 6.43E-11 | -0.69023 | 1.69E-14 |
| Ept+Sc | Hbb-bs | -0.55372 | 9.50E-10 | -0.49658 | 3.52E-11 | -0.52789 | 1.64E-12 |
| Ept+Sc | Hbb-bt | -1.0417 | 2.61E-27 | -0.88425 | 9.41E-30 | -0.95728 | 6.17E-40 |
| Ept+Sc | Hbb-y | -1.3275 | 1.16E-29 | -1.0817 | 1.10E-21 | -1.1934 | 4.80E-25 |
| IO+Ml | Crybb3 | -1.9211 | 5.33E-12 | -0.88111 | 0.00042461 | -1.5925 | 8.7429e-17402 |
| IO+Sc | Gpx1 | -0.53903 | 0.00041264 | -0.54152 | 1.47E-05 | -0.56814 | 1.68E-05 |
| JTP+O | Fth1 | -0.58151 | 0.0086744 | -1.1146 | 0.00081282 | -0.73544 | 0.0010683 |
| JTP+O | Hba-x | -0.9451 | 0.0088024 | -1.853 | 4.14E-05 | -1.2526 | 1.92E-06 |
| JTP+O | Hbb-y | -0.97924 | 1.04E-06 | -0.86671 | 0.0013237 | -0.94426 | 1.60E-08 |
| JTP+Sc | Fth1 | -0.61703 | 5.20E-05 | -1.1294 | 9.50E-91 | -0.89866 | 2.14E-67 |
| JTP+Sc | Hba-x | -0.65172 | 0.00033793 | -1.8333 | 9.97E-116 | -1.3027 | 3.44E-47 |
| JTP+Sc | Hbb-y | -0.47563 | 0.0045083 | -0.51965 | 2.26E-14 | -0.49905 | 6.19E-14 |
| My+Sc | Hbb-y | -0.51683 | 2.56E-09 | -0.85617 | 7.02E-26 | -0.72839 | 5.50E-20 |
| Sc+En | Hba-x | -0.47477 | 2.13E-17 | -0.87995 | 3.24E-05 | -0.58358 | 4.65E-29 |

* The full names of heterotypic pairs can be found in Table S5.
* The logFC was calculated based on the natural logarithm.

### Table S1E. Up-regulated genes identified from the mouse liver cancer Slide-seq data

| Heterotypic pair | Up-regulated_gene | Expression in snRNA-seq | Homotypic1_logFC | Hmotypic1_p.value | Homotypic2_logFC | Homotypic2_p.value | Artificial_logFC | Artificial_fdr |
|---|---|---|---|---|---|---|---|---|
| HepatocyteI+HepatocyteII | Arg1 | HepatocyteII | 0.62288 | 8.49E-55 | 0.40004 | 4.94E-10 | 0.52801 | 3.09E-46 |
| HepatocyteI+HepatocyteII | Gnmt | HepatocyteII | 0.59362 | 3.28E-52 | 0.65088 | 2.61E-26 | 0.61473 | 2.40E-63 |
| HepatocyteI+HepatocyteII | Hrg | HepatocyteII | 0.43573 | 2.44E-33 | 0.78861 | 1.18E-36 | 0.57859 | 7.65E-76 |
| HepatocyteI+HepatocyteII | Hsd17b13 | HepatocyteII | 0.54793 | 3.67E-42 | 0.76199 | 7.51E-33 | 0.6384 | 1.29E-63 |
| HepatocyteI+HepatocyteII | Igf1 | HepatocyteII | 0.4219 | 8.91E-29 | 0.5333 | 1.60E-21 | 0.46621 | 1.05E-36 |
| HepatocyteI+HepatocyteII | Itih4 | HepatocyteI | 0.4228 | 1.78E-28 | 0.56151 | 3.33E-23 | 0.48642 | 3.72E-41 |
| HepatocyteI+HepatocyteII | Mat1a | HepatocyteII | 0.49711 | 3.80E-42 | 0.78745 | 6.53E-37 | 0.61794 | 5.32E-87 |
| HepatocyteI+HepatocyteII | Rnase4 | HepatocyteI | 0.40484 | 4.08E-29 | 0.4621 | 1.35E-18 | 0.42903 | 8.00E-37 |
| HepatocyteI+HepatocyteII | Saa4 | HepatocyteII | 0.49382 | 1.24E-54 | 0.55726 | 3.07E-21 | 0.52132 | 4.51E-76 |
| HepatocyteI+HepatocyteII | Serpina1c | HepatocyteI | 0.55279 | 2.60E-48 | 0.57904 | 4.42E-22 | 0.56122 | 1.49E-56 |
| HepatocyteI+HepatocyteII | Serpina3m | HepatocyteII | 0.58118 | 2.16E-54 | 0.56146 | 2.16E-22 | 0.57417 | 1.02E-59 |
| HepatocyteI+HepatocyteII | Tdo2 | HepatocyteII | 0.55293 | 5.31E-49 | 0.71628 | 1.93E-32 | 0.62124 | 3.27E-69 |
| HepatocyteI+Monocyte | Cd163 | Monocyte | 0.71129 | 1.77E-28 | 0.40011 | 0.00062417 | 0.59263 | 7.14E-20 |
| HepatocyteI+Monocyte | Cd5l | Monocyte | 1.1518 | 5.03E-58 | 0.54006 | 2.24E-05 | 0.91359 | 1.18E-38 |
| HepatocyteI+Monocyte | Clec4f | Monocyte | 1.4463 | 7.12E-77 | 0.79691 | 5.67E-08 | 1.1963 | 2.27E-56 |
| HepatocyteI+Monocyte | Marco | Monocyte | 1.0521 | 1.93E-49 | 0.45206 | 0.00044519 | 0.82002 | 1.15E-31 |
| HepatocyteI+Monocyte | Slc40a1 | Monocyte | 0.95693 | 1.52E-47 | 0.59722 | 4.56E-07 | 0.81846 | 1.01E-35 |
| HepatocyteI+VSMC | Lcn2 | HepatocyteI | 0.66993 | 4.49E-05 | 0.79342 | 0.0016655 | 0.71638 | 0.00011444 |
| HepatocyteII+Monocyte | Alcam | HepatocyteII | 0.5703 | 2.11E-05 | 0.67359 | 4.45E-05 | 0.60317 | 7.94E-05 |
| HepatocyteII+Monocyte | Clec4f | Monocyte | 0.42068 | 0.0018482 | 0.82395 | 1.30E-08 | 0.59209 | 3.57E-05 |
| HepatocyteII+Monocyte | Hsd17b13 | HepatocyteII | 0.40411 | 0.00046946 | 0.51224 | 3.79E-05 | 0.4596 | 0.00020635 |
| HepatocyteII+Monocyte | Mup20 | HepatocyteII | 1.7582 | 8.76E-25 | 0.65591 | 0.00051659 | 1.2888 | 2.23E-15 |
| HepatocyteII+Monocyte | Neat1 | HepatocyteII | 0.53608 | 2.26E-05 | 0.97412 | 1.85E-10 | 0.71883 | 8.46E-07 |
| HepatocyteII+Monocyte | Slc40a1 | Monocyte | 0.61883 | 6.71E-05 | 1.3162 | 4.25E-14 | 0.91609 | 7.10E-09 |
| HepatocyteII+Monocyte | Vti1a | Monocyte | 1.0223 | 5.99E-13 | 0.50805 | 0.0016665 | 0.81251 | 1.26E-08 |
| HepatocyteII+VSMC | Dcn | VSMC | 1.2078 | 0.00015387 | 0.94178 | 0.0031563 | 1.1196 | 0.0013113 |
| HepatocyteII+VSMC | Hsd17b13 | HepatocyteII | 1.1466 | 3.97E-06 | 1.5216 | 1.91E-05 | 1.2954 | 0.00036754 |
| HepatocyteII+VSMC | Mup20 | HepatocyteII | 0.85234 | 0.0027371 | 0.80629 | 0.0031393 | 0.84806 | 0.0092178 |
| HepatocyteII+VSMC | Neat1 | HepatocyteII | 0.81442 | 0.00046198 | 1.1755 | 0.00011142 | 0.95804 | 0.0023528 |
| HepatocyteII+VSMC | Sorbs2 | HepatocyteII | 1.0912 | 0.00015079 | 1.3817 | 8.41E-06 | 1.1873 | 3.16E-07 |
| HepatocyteII+VSMC | Zfand3 | HepatocyteII | 0.92087 | 0.00050076 | 0.66193 | 0.0093101 | 0.84644 | 0.0040735 |
| TumorII+HepatocyteI | Spp1 | HepatocyteI | 0.897 | 1.67E-23 | 1.3584 | 8.59E-50 | 1.054 | 1.74E-32 |
| TumorIII+HepatocyteI | Spp1 | TumorIII | 0.54735 | 8.11E-11 | 1.9888 | 3.90E-85 | 1.0455 | 2.18E-32 |
| TumorIII+HepatocyteI | Tuba1a | HepatocyteI | 0.41681 | 2.08E-09 | 0.90641 | 5.61E-34 | 0.58493 | 1.10E-15 |
| TumorIII+HepatocyteII | Ext1 | TumorIII | 1.3015 | 6.91E-05 | 0.66989 | 0.0061089 | 1.1311 | 0.0010699 |
| TumorIII+HepatocyteII | Rbms1 | TumorIII | 1.0154 | 0.0011967 | 0.87535 | 0.0043947 | 0.97584 | 0.0044819 |
| TumorIII+HepatocyteII | Sorbs2 | HepatocyteII | 1.0562 | 0.0015383 | 1.1609 | 0.0006242 | 1.0846 | 0.0031706 |
| TumorIII+HepatocyteII | Tgfbr1 | HepatocyteII | 0.93878 | 0.00026837 | 1.4034 | 6.70E-05 | 1.0853 | 0.003113 |
| TumorIII+Monocyte | F13a1 | Monocyte | 0.94129 | 1.04E-42 | 0.96804 | 4.79E-20 | 0.94445 | 1.01E-43 |
| VSMC+TumorIII | Col15a1 | VSMC | 0.9453 | 0.00012559 | 2.0481 | 2.28E-29 | 1.7886 | 1.19E-23 |
| VSMC+TumorIII | Col1a2 | VSMC | 0.66008 | 0.0082911 | 1.7252 | 2.99E-23 | 1.4648 | 1.38E-17 |
| VSMC+TumorIII | Col3a1 | VSMC | 0.49994 | 0.00028959 | 0.64032 | 2.04E-08 | 0.6118 | 5.84E-07 |

* The logFC was calculated based on the natural logarithm.

### Table S1F. Down-regulated genes identified from the mouse liver cancer Slide-seq data

| Heterotypic pair | Down-regulated_gene | Homotypic1_logFC | Hmotypic1_p.value | Homotypic2_logFC | Homotypic2_p.value | Artificial_logFC | Artificial_fdr |
|---|---|---|---|---|---|---|---|
| TumorII+TumorIII | Ahnak | -0.92843 | 2.37E-11 | -0.57872 | 0.00022657 | -0.79487 | 3.83E-07 |
| TumorII+TumorIII | Fgb | -0.43377 | 0.0090293 | -0.58678 | 0.00029071 | -0.51076 | 0.0028919 |
| TumorIII+TumorI | Birc5 | -0.80291 | 8.09E-40 | -0.53643 | 1.61E-05 | -0.66475 | 1.24E-28 |

* The logFC was calculated based on the natural logarithm.

### Table S1G. Up-regulated genes identified from the mouse hippocampus Slide-seq data

| Heterotypic pair | Up-regulated_gene | Expression in scRNA-seq | Homotypic1_logFC | Hmotypic1_p.value | Homotypic2_logFC | Homotypic2_p.value | Artificial_logFC | Artificial_fdr |
|---|---|---|---|---|---|---|---|---|
| A+O | Dbi | A | 0.51438 | 2.65E-16 | 4.57E-01 | 3.83E-13 | 4.86E-01 | 4.99E-16 |
| CA3+D | Igfbp4 | CA3 | 0.75045 | 0.0007722 | 0.74287 | 0.00085865 | 0.74925 | 0.0015419 |
| CA3+In | Igfbp4 | In | 0.75959 | 1.12E-05 | 5.19E-01 | 0.0020205 | 0.65019 | 0.00061727 |
| Ch+EnT | Igfbp4 | EnT | 0.99726 | 2.45E-07 | 6.78E-01 | 0.0037621 | 0.91887 | 8.88E-06 |
| Ch+EnT | Igfbp7 | EnT | 0.49497 | 0.00026967 | 0.49497 | 0.00026967 | 0.49497 | 0.0033361 |
| Ch+EnT | Inmt | EnT | 1.2347 | 1.79E-07 | 9.16E-01 | 0.00077618 | 1.1679 | 1.24E-06 |
| Ch+EnT | Lum | EnT | 0.46733 | 0.00094173 | 0.46139 | 0.0012779 | 0.46014 | 0.0065985 |
| Ch+EnT | Mgp | EnT | 0.93889 | 4.63E-07 | 7.82E-01 | 0.00016544 | 0.90475 | 5.85E-06 |
| EnT+A | Dbi | A | 0.55148 | 6.50E-05 | 5.74E-01 | 1.44E-06 | 5.70E-01 | 1.59E-05 |
| EnT+A | Fabp7 | A | 1.0725 | 6.59E-05 | 1.11E+00 | 4.62E-14 | 1.10E+00 | 1.25E-09 |
| EnT+A | Trf | EnT | 0.62715 | 0.00070979 | 0.69793 | 5.67E-07 | 6.65E-01 | 1.58E-05 |
| EnT+Ento | Necab2 | Ento | 0.83981 | 0.0012975 | 0.50315 | 0.00037474 | 0.61047 | 4.10E-05 |
| EnT+Ento | Plp1 | EnT | 0.61612 | 1.86E-08 | 5.24E-01 | 2.27E-07 | 5.51E-01 | 2.81E-07 |
| EnT+In | Agt | EnT | 0.54965 | 8.47E-05 | 4.89E-01 | 0.00043651 | 0.50911 | 0.0016552 |
| EnT+In | Calb2 | In | 0.8114 | 0.0061183 | 0.77837 | 2.11E-05 | 8.16E-01 | 0.00063365 |
| EnT+In | Mbp | EnT | 1.0856 | 8.18E-07 | 9.62E-01 | 1.68E-06 | 9.86E-01 | 7.74E-06 |
| EnT+In | Nwd2 | In | 0.68325 | 3.28E-06 | 4.59E-01 | 0.0013244 | 0.52674 | 0.001497 |
| EnT+In | Pcp4 | In | 0.6552 | 0.0001213 | 0.42971 | 0.0068586 | 0.49034 | 0.0098786 |
| EnT+In | Plp1 | EnT | 0.42473 | 0.0028153 | 0.43804 | 0.00089355 | 0.43732 | 0.0051391 |
| EnT+In | Tcf7l2 | EnT | 1.2846 | 5.60E-06 | 7.38E-01 | 0.00014468 | 0.88967 | 0.00013052 |
| EnT+In | Tspan13 | In | 1.0226 | 0.0015014 | 0.75457 | 0.00056653 | 0.82991 | 0.0011757 |
| Ento+Ch | Aifm3 | Ch | 0.75172 | 5.48E-13 | 6.70E-01 | 2.91E-08 | 7.13E-01 | 5.96E-11 |
| Ento+Ch | Apod | Ch | 0.76661 | 1.81E-14 | 7.17E-01 | 7.19E-11 | 7.35E-01 | 1.76E-12 |
| Ento+Ch | Calb2 | Ento | 0.68246 | 3.53E-06 | 1.53E+00 | 2.20E-15 | 1.11E+00 | 1.82E-12 |
| Ento+Ch | Cpne9 | Ento | 0.42268 | 4.11E-06 | 4.29E-01 | 0.00067771 | 0.42709 | 1.29E-05 |
| Ento+Ch | Gm5741 | Ento | 0.76755 | 8.77E-13 | 8.10E-01 | 5.08E-12 | 7.89E-01 | 2.53E-12 |
| Ento+Ch | Igfbp7 | Ch | 0.62413 | 6.36E-12 | 6.45E-01 | 1.43E-12 | 6.33E-01 | 3.99E-11 |
| Ento+Ch | Malat1 | Ento | 0.41172 | 1.29E-07 | 4.37E-01 | 4.27E-08 | 4.27E-01 | 3.70E-07 |
| Ento+Ch | Meg3 | Ento | 0.85009 | 3.07E-14 | 9.11E-01 | 4.00E-13 | 8.80E-01 | 5.82E-14 |
| Ento+Ch | Msi2 | Ch | 0.8164 | 1.14E-13 | 7.55E-01 | 3.77E-08 | 7.84E-01 | 6.94E-12 |
| Ento+Ch | Necab2 | Ch | 0.80895 | 1.89E-13 | 7.43E-01 | 7.51E-08 | 7.74E-01 | 1.07E-11 |
| Ento+Ch | Nnat | Ento | 0.48987 | 1.94E-08 | 5.05E-01 | 2.17E-06 | 4.93E-01 | 1.12E-07 |
| Ento+Ch | Nwd2 | Ento | 0.41923 | 1.54E-06 | 5.52E-01 | 3.83E-10 | 4.84E-01 | 2.51E-07 |
| Ento+Ch | Plp1 | Ento | 0.57372 | 5.37E-10 | 5.43E-01 | 1.28E-06 | 5.60E-01 | 8.44E-09 |
| Ento+Ch | Scube1 | Ento | 0.40368 | 0.0087653 | 2.404 | 1.71E-23 | 1.42E+00 | 4.06E-29 |
| Ento+Ch | Tac2 | Ento | 0.5235 | 5.37E-10 | 4.48E-01 | 8.15E-06 | 4.87E-01 | 5.03E-08 |
| Ento+Ch | Zcchc12 | Ento | 0.40744 | 1.57E-06 | 4.44E-01 | 7.11E-05 | 4.23E-01 | 2.97E-06 |
| Ento+Ch | Zic1 | Ch | 0.54002 | 6.06E-05 | 1.59E+00 | 6.73E-09 | 1.08E+00 | 8.48E-15 |
| Ento+N | Gda | Ento | 0.44386 | 9.23E-06 | 4.42E-01 | 0.00014454 | 0.43996 | 0.00010034 |
| Ep+Ca | Malat1 | Ep | 1.6242 | 0.00020585 | 2.6767 | 0.00012208 | 2.1159 | 7.33E-06 |
| Ep+Ca | Ptgds | Ca | 1.1297 | 0.0012242 | 2.0266 | 0.00019889 | 1.5522 | 1.61E-05 |
| Ep+Ca | Ttr | Ca | 1.0958 | 0.00010998 | 2.1654 | 0.0021327 | 1.5615 | 1.82E-20 |
| Ep+N | Calb2 | N | 0.81646 | 0.00018498 | 0.83612 | 5.20E-05 | 8.17E-01 | 0.00021087 |
| Ep+O | Ndrg2 | Ep | 0.79589 | 0.0033705 | 1.0556 | 3.23E-05 | 9.28E-01 | 0.0013186 |
| Ep+O | Zbtb20 | Ep | 1.3841 | 1.34E-05 | 8.95E-01 | 0.0011392 | 1.1491 | 0.00022428 |
| In+A | Mbp | A | 0.64121 | 1.90E-22 | 6.47E-01 | 1.29E-35 | 6.48E-01 | 1.77E-53 |
| In+A | Pcp4 | In | 0.79467 | 2.45E-30 | 5.85E-01 | 5.99E-26 | 7.09E-01 | 1.90E-61 |
| In+A | Plp1 | A | 0.74492 | 4.94E-22 | 1.19E+00 | 3.37E-98 | 9.25E-01 | 4.72E-85 |
| In+Ca | Apba1 | In | 0.46601 | 3.69E-08 | 1.07E+00 | 0.0024075 | 0.69532 | 1.06E-18 |
| In+Ca | Cygb | In | 0.90323 | 7.99E-33 | 9.52E-01 | 0.0031601 | 0.92194 | 5.84E-33 |
| In+Ca | Gucy1a3 | Ca | 1.8559 | 3.22E-77 | 1.27E+00 | 0.0063407 | 1.6316 | 5.59E-64 |
| In+Ca | Ptgds | Ca | 0.6197 | 1.82E-14 | 8.52E-01 | 0.0059904 | 0.70573 | 7.91E-21 |
| In+Ca | Scube1 | In | 0.64103 | 3.57E-24 | 7.26E-01 | 0.0066859 | 0.67308 | 3.46E-26 |
| In+Ca | Snhg11 | In | 0.69575 | 5.17E-19 | 9.05E-01 | 0.0057038 | 0.7736 | 1.70E-25 |
| In+Ca | Zcchc12 | In | 0.53152 | 5.07E-07 | 1.66E+00 | 0.001002 | 0.95525 | 7.09E-25 |
| In+Ca | Zic1 | Ca | 0.52066 | 6.98E-14 | 8.18E-01 | 0.0042108 | 0.62491 | 4.69E-21 |
| In+D | Cck | In | 0.47476 | 0.0032986 | 0.69024 | 0.00010898 | 0.60595 | 0.001849 |
| In+Ento | Amotl1 | In | 0.97627 | 3.93E-29 | 1.06E+00 | 2.42E-37 | 1.02E+00 | 2.17E-37 |
| In+Ento | Atp2b1 | Ento | 0.51685 | 1.29E-15 | 6.71E-01 | 1.91E-28 | 5.90E-01 | 1.18E-23 |
| In+Ento | Cck | In | 1.2405 | 2.27E-53 | 4.21E-01 | 5.51E-08 | 8.34E-01 | 9.80E-33 |
| In+Ento | Cpne7 | In | 0.81806 | 4.21E-32 | 8.99E-01 | 7.82E-41 | 8.58E-01 | 1.87E-38 |
| In+Ento | Ctnna2 | In | 1.0394 | 6.50E-43 | 1.03E+00 | 1.03E-43 | 1.03E+00 | 7.92E-45 |
| In+Ento | Epha4 | In | 0.70947 | 2.52E-28 | 7.67E-01 | 3.23E-35 | 7.34E-01 | 7.76E-34 |
| In+Ento | Gabbr1 | In | 0.90835 | 5.13E-29 | 1.05E+00 | 4.42E-43 | 9.82E-01 | 1.91E-40 |
| In+Ento | Gabbr2 | In | 0.43634 | 9.85E-11 | 4.39E-01 | 2.14E-12 | 4.39E-01 | 7.64E-14 |
| In+Ento | Grm1 | In | 0.52168 | 7.75E-18 | 5.54E-01 | 1.86E-21 | 5.40E-01 | 1.25E-21 |
| In+Ento | Ildr2 | In | 0.95981 | 1.27E-32 | 7.46E-01 | 7.83E-25 | 8.55E-01 | 2.90E-34 |
| In+Ento | Kcnip4 | Ento | 0.46702 | 1.50E-15 | 5.56E-01 | 5.61E-24 | 5.12E-01 | 8.79E-22 |
| In+Ento | Lrrtm1 | In | 0.78838 | 4.49E-33 | 7.98E-01 | 1.05E-35 | 7.92E-01 | 1.38E-36 |

| | | | | | | | | |
|---|---|---|---|---|---|---|---|---|
| In+Ento | Meg3 | In | 0.77715 | 3.93E-37 | 6.96E-01 | 7.50E-31 | 7.37E-01 | 4.67E-35 |
| In+Ento | Ncdn | Ento | 0.44254 | 4.30E-16 | 4.36E-01 | 4.20E-17 | 4.38E-01 | 1.44E-18 |
| In+Ento | Nrip3 | In | 0.68042 | 2.73E-13 | 6.48E-01 | 3.57E-15 | 6.55E-01 | 5.18E-20 |
| In+Ento | Nrxn1 | Ento | 0.73787 | 3.72E-14 | 7.78E-01 | 9.66E-20 | 7.47E-01 | 8.30E-25 |
| In+Ento | Ntng1 | In | 0.9339 | 3.03E-37 | 8.07E-01 | 8.20E-30 | 8.67E-01 | 7.23E-37 |
| In+Ento | Pcp4l1 | In | 0.59169 | 5.83E-15 | 5.10E-01 | 1.07E-13 | 5.51E-01 | 1.84E-17 |
| In+Ento | Plcb4 | Ento | 0.49114 | 5.11E-14 | 7.25E-01 | 2.44E-33 | 6.07E-01 | 2.27E-25 |
| In+Ento | Prkcd | In | 0.48007 | 5.52E-19 | 5.22E-01 | 8.96E-24 | 4.99E-01 | 2.43E-22 |
| In+Ento | Ptpn3 | Ento | 0.49644 | 5.99E-18 | 5.65E-01 | 4.83E-25 | 5.28E-01 | 1.90E-23 |
| In+Ento | Rab3c | In | 0.45947 | 7.81E-21 | 4.35E-01 | 2.19E-19 | 4.46E-01 | 5.41E-21 |
| In+Ento | Ramp3 | Ento | 0.73401 | 1.01E-32 | 6.31E-01 | 1.38E-25 | 6.80E-01 | 4.00E-31 |
| In+Ento | Rasgrf1 | Ento | 0.55157 | 3.71E-17 | 5.50E-01 | 3.41E-19 | 5.52E-01 | 1.47E-21 |
| In+Ento | Rgs4 | Ento | 0.82514 | 1.95E-35 | 7.26E-01 | 2.32E-29 | 7.73E-01 | 6.95E-35 |
| In+Ento | Rora | In | 0.47058 | 1.05E-08 | 1.11E+00 | 1.39E-51 | 7.88E-01 | 5.05E-31 |
| In+Ento | Shox2 | Ento | 0.62778 | 1.31E-23 | 6.21E-01 | 8.64E-25 | 6.25E-01 | 6.04E-27 |
| In+Ento | Slc17a6 | Ento | 0.56125 | 8.05E-25 | 4.96E-01 | 1.16E-20 | 5.30E-01 | 1.29E-24 |
| In+Ento | Slc24a2 | In | 0.64627 | 5.49E-19 | 6.54E-01 | 4.41E-22 | 6.48E-01 | 8.50E-25 |
| In+Ento | Slc24a3 | In | 0.83725 | 2.66E-32 | 4.50E-01 | 1.89E-11 | 6.46E-01 | 9.43E-25 |
| In+Ento | Snhg11 | In | 0.44477 | 8.96E-08 | 1.05E+00 | 1.51E-44 | 7.42E-01 | 1.90E-28 |
| In+Ento | Synpo2 | Ento | 0.55535 | 2.13E-11 | 6.30E-01 | 1.30E-16 | 5.89E-01 | 4.98E-19 |
| In+Ento | Tcf7l2 | Ento | 0.7083 | 4.28E-17 | 6.50E-01 | 6.08E-17 | 6.86E-01 | 2.14E-24 |
| In+Ento | Tnnt1 | In | 0.52988 | 1.11E-13 | 6.79E-01 | 7.13E-25 | 6.04E-01 | 4.13E-23 |
| In+Ento | Zic1 | In | 0.79954 | 1.62E-23 | 7.68E-01 | 1.54E-28 | 7.87E-01 | 2.25E-32 |
| In+Ep | A830039N20Rik | In | 1.2334 | 3.65E-12 | 8.36E-01 | 0.0045363 | 1.0766 | 2.04E-09 |
| In+Ep | Aifm3 | Ep | 0.73046 | 1.45E-09 | 8.56E-01 | 8.61E-09 | 7.84E-01 | 3.22E-10 |
| In+Ep | Cacna1e | In | 1.0881 | 5.76E-20 | 1.00E+00 | 1.51E-12 | 1.05E+00 | 5.77E-18 |
| In+Ep | Cacnb3 | In | 0.856 | 4.28E-17 | 8.42E-01 | 2.79E-13 | 8.43E-01 | 8.99E-16 |
| In+Ep | Cadps2 | In | 1.163 | 2.90E-20 | 8.21E-01 | 0.00030781 | 1.0403 | 3.04E-14 |
| In+Ep | Calb2 | In | 0.58599 | 3.79E-09 | 7.33E-01 | 2.75E-09 | 6.45E-01 | 2.98E-10 |
| In+Ep | Cers4 | In | 0.53984 | 2.24E-10 | 5.31E-01 | 1.05E-07 | 5.34E-01 | 1.94E-09 |
| In+Ep | Cpne4 | In | 1.6237 | 1.13E-33 | 1.77E+00 | 1.19E-29 | 1.68E+00 | 6.81E-34 |
| In+Ep | Cpne9 | In | 0.96003 | 6.77E-20 | 9.28E-01 | 4.28E-15 | 9.44E-01 | 2.24E-18 |
| In+Ep | Cygb | In | 0.7135 | 7.85E-15 | 7.35E-01 | 7.71E-15 | 7.23E-01 | 3.49E-14 |
| In+Ep | Fabp7 | Ep | 0.7728 | 1.60E-14 | 8.78E-01 | 6.29E-18 | 8.13E-01 | 7.67E-15 |
| In+Ep | Fgf1 | In | 0.4671 | 1.95E-09 | 5.34E-01 | 6.50E-12 | 4.93E-01 | 1.40E-09 |
| In+Ep | Gabbr1 | In | 0.54238 | 1.09E-07 | 7.36E-01 | 1.89E-07 | 6.11E-01 | 4.04E-09 |
| In+Ep | Gfra1 | In | 0.6636 | 1.16E-12 | 6.97E-01 | 9.52E-14 | 6.75E-01 | 4.66E-12 |
| In+Ep | Gm5741 | In | 1.3212 | 3.06E-25 | 1.33E+00 | 1.62E-14 | 1.32E+00 | 1.86E-24 |
| In+Ep | Gng8 | In | 0.61247 | 5.36E-13 | 5.82E-01 | 3.01E-08 | 6.01E-01 | 1.02E-11 |
| In+Ep | Gucy1a3 | In | 1.3246 | 1.54E-26 | 1.25E+00 | 1.96E-13 | 1.29E+00 | 9.13E-25 |
| In+Ep | Kcnd2 | In | 0.41336 | 1.96E-06 | 5.57E-01 | 2.50E-06 | 4.71E-01 | 1.26E-07 |
| In+Ep | Kcnip1 | In | 0.73477 | 3.08E-13 | 5.79E-01 | 0.00010862 | 0.67777 | 5.67E-11 |
| In+Ep | Kcnma1 | In | 0.4922 | 4.27E-08 | 6.08E-01 | 1.43E-11 | 5.34E-01 | 1.36E-08 |
| In+Ep | Lrrc55 | In | 0.69844 | 5.04E-14 | 6.05E-01 | 4.99E-07 | 6.56E-01 | 6.85E-12 |
| In+Ep | Malat1 | Ep | 0.4154 | 1.30E-05 | 5.74E-01 | 1.97E-05 | 4.72E-01 | 1.21E-06 |
| In+Ep | Meg3 | In | 0.55618 | 1.68E-11 | 5.83E-01 | 1.82E-12 | 5.67E-01 | 6.54E-11 |
| In+Ep | Ncald | In | 0.67261 | 9.71E-12 | 7.93E-01 | 1.68E-15 | 7.18E-01 | 3.02E-12 |
| In+Ep | Ndrg2 | Ep | 0.4943 | 2.47E-07 | 8.78E-01 | 1.03E-11 | 6.56E-01 | 3.50E-10 |
| In+Ep | Necab2 | In | 0.46244 | 1.95E-08 | 5.11E-01 | 6.77E-07 | 4.86E-01 | 9.92E-09 |
| In+Ep | Necab3 | Ep | 0.71065 | 1.50E-12 | 8.99E-01 | 1.93E-17 | 7.84E-01 | 3.70E-14 |
| In+Ep | Nwd2 | In | 0.41882 | 3.14E-08 | 5.03E-01 | 2.62E-11 | 4.53E-01 | 1.04E-08 |
| In+Ep | Pcp4 | In | 0.69581 | 8.82E-13 | 8.33E-01 | 3.10E-17 | 7.50E-01 | 1.26E-13 |
| In+Ep | Pcsk2 | In | 0.64838 | 1.74E-09 | 9.61E-01 | 3.86E-15 | 7.70E-01 | 3.04E-12 |
| In+Ep | Pde2a | In | 0.63786 | 3.78E-12 | 6.37E-01 | 4.70E-08 | 6.37E-01 | 1.90E-11 |
| In+Ep | Plp1 | In | 0.70289 | 5.03E-14 | 7.69E-01 | 1.55E-14 | 7.28E-01 | 5.14E-14 |
| In+Ep | Pnmal2 | In | 1.0502 | 1.81E-16 | 2.64E+00 | 1.03E-40 | 1.68E+00 | 8.93E-65 |
| In+Ep | Pou4f1 | In | 1.0117 | 2.55E-16 | 2.96E+00 | 4.46E-27 | 1.79E+00 | 8.77E-42 |
| In+Ep | Qdpr | In | 0.93349 | 7.37E-17 | 9.72E-01 | 1.53E-11 | 9.47E-01 | 1.27E-16 |
| In+Ep | Rbm25 | In | 0.86148 | 2.68E-15 | 8.50E-01 | 2.31E-08 | 8.55E-01 | 1.30E-14 |
| In+Ep | Rtn1 | In | 0.44583 | 7.28E-07 | 6.75E-01 | 9.23E-14 | 5.29E-01 | 1.40E-08 |
| In+Ep | Scube1 | In | 0.47246 | 5.76E-06 | 7.26E-01 | 2.31E-07 | 5.77E-01 | 5.60E-08 |
| In+Ep | Slc17a6 | In | 0.54904 | 3.00E-10 | 6.59E-01 | 1.38E-12 | 5.95E-01 | 5.28E-11 |
| In+Ep | Snca | In | 0.60609 | 4.90E-11 | 6.61E-01 | 1.00E-09 | 6.27E-01 | 5.36E-11 |
| In+Ep | Sncg | Ep | 0.63444 | 1.59E-11 | 7.35E-01 | 3.65E-14 | 6.78E-01 | 4.07E-12 |
| In+Ep | Snhg11 | In | 0.47433 | 7.04E-08 | 6.04E-01 | 1.55E-09 | 5.27E-01 | 6.78E-09 |
| In+Ep | St6galnac5 | In | 0.50624 | 2.38E-09 | 5.61E-01 | 5.78E-10 | 5.26E-01 | 2.92E-09 |
| In+Ep | Synpr | In | 0.45275 | 5.56E-07 | 6.61E-01 | 5.79E-10 | 5.38E-01 | 6.98E-09 |
| In+Ep | Syt6 | In | 0.5713 | 2.03E-08 | 6.76E-01 | 0.0001678 | 0.61818 | 3.08E-08 |
| In+Ep | Syt9 | In | 0.92919 | 1.05E-12 | 1.80E+00 | 3.59E-17 | 1.27E+00 | 7.28E-21 |
| In+Ep | Tac2 | In | 0.47208 | 3.75E-06 | 5.60E-01 | 0.00067594 | 0.50504 | 1.29E-06 |

| | | | | | | | | |
|---|---|---|---|---|---|---|---|---|
| In+Ep | Tcf7l2 | Ep | 0.57185 | 4.35E-06 | 1.50E+00 | 2.03E-18 | 9.35E-01 | 1.22E-14 |
| In+Ep | Tesc | In | 0.46556 | 4.84E-06 | 8.48E-01 | 1.72E-08 | 6.19E-01 | 4.89E-09 |
| In+Ep | Tspan13 | Ep | 0.58504 | 7.55E-12 | 4.59E-01 | 7.66E-05 | 5.34E-01 | 1.76E-09 |
| In+Ep | Ttr | Ep | 0.44813 | 8.14E-05 | 9.86E-01 | 1.56E-09 | 6.57E-01 | 5.16E-09 |
| In+Ep | Zcchc12 | In | 0.73912 | 1.11E-08 | 9.82E-01 | 6.30E-05 | 8.36E-01 | 2.39E-14 |
| In+Ep | Zic1 | Ep | 1.0819 | 5.12E-19 | 9.73E-01 | 0.00054513 | 1.0453 | 1.10E-20 |
| In+Ep | Zic4 | Ep | 0.49514 | 4.63E-05 | 9.34E-01 | 6.36E-06 | 6.65E-01 | 2.33E-08 |
| In+MM | Pcp4 | MM | 0.46836 | 0.004549 | 0.83857 | 0.00010221 | 0.59475 | 0.0047964 |
| In+MM | Plp1 | MM | 0.67076 | 0.00072455 | 1.0985 | 1.14E-05 | 7.92E-01 | 0.00077291 |
| In+O | Ndrg2 | O | 0.43176 | 2.51E-22 | 6.24E-01 | 1.52E-67 | 5.12E-01 | 1.51E-47 |
| In+O | Pcp4 | In | 0.87329 | 2.18E-26 | 1.55E+00 | 1.79E-132 | 1.15E+00 | 1.54E-89 |
| In+O | Zic1 | O | 0.66487 | 2.06E-20 | 6.15E-01 | 2.28E-28 | 6.46E-01 | 2.75E-43 |
| N+O | Pcp4 | N | 0.71763 | 1.29E-16 | 7.86E-01 | 4.59E-27 | 7.41E-01 | 2.58E-26 |
| O+P | Gm15440 | O | 0.47888 | 0.00053118 | 0.72965 | 3.65E-07 | 5.99E-01 | 0.00011635 |
| O+P | Nfasc | P | 0.59026 | 0.00030248 | 0.90565 | 1.21E-06 | 7.46E-01 | 4.02E-05 |
| O+P | Slc44a1 | O | 0.6113 | 0.00010042 | 0.70517 | 6.72E-05 | 6.60E-01 | 0.00017071 |
| * The full names of heterotypic pairs can be found in Table S5. | | | | | | | | |
| * The logFC was calculated based on the natural logarithm. | | | | | | | | |

#### Table S1H. Down-regulated genes identified from the mouse hippocampus Slide-seq data

| Heterotypic pair | Down-regulated_gene | Homotypic1_logFC | Hmotypic1_p.value | Homotypic2_logFC | Homotypic2_p.value | Artificial_logFC | Artificial_fdr |
|---|---|---|---|---|---|---|---|
| CA1+Ch | Pcp4 | -0.58582 | 0.0079524 | -0.78776 | 0.0069514 | -0.63339 | 0.005624 |
| CA1+MM | Pcp4 | -0.6172 | 1.59E-05 | -0.51347 | 0.0075461 | -0.58006 | 1.03E-05 |
| CA3+D | Atp2b1 | -0.66151 | 0.0052972 | -0.66139 | 0.0024592 | -0.66558 | 6.97E-03 |
| CA3+D | Cpne6 | -0.86892 | 0.00010443 | -0.5882 | 0.0026289 | -0.74567 | 4.27E-06 |
| CA3+D | Epha4 | -0.5355 | 9.01E-07 | -1.1234 | 1.38E-16 | -0.80337 | 1.30E-10 |
| CA3+D | Gabbr1 | -0.66971 | 0.00012336 | -0.64821 | 0.00093077 | -0.66823 | 2.47E-05 |
| CA3+D | Lppr4 | -0.60825 | 0.00021775 | -0.66808 | 5.68E-05 | -0.61822 | 4.45E-05 |
| CA3+D | Nefm | -1.508 | 3.06E-13 | -0.67626 | 5.12E-05 | -1.1398 | 4.85E-11 |
| CA3+D | Olfm1 | -1.0923 | 4.22E-05 | -1.8462 | 9.90E-08 | -1.4314 | 2.94E-05 |
| CA3+D | Syn2 | -0.62215 | 0.0034671 | -0.61528 | 0.0022759 | -0.63805 | 6.03E-05 |
| D+CA1 | Ank3 | -0.56066 | 0.0032544 | -0.59921 | 0.00093614 | -0.57609 | 3.16E-05 |
| D+CA1 | Cadm2 | -0.62087 | 1.14E-05 | -0.74449 | 4.02E-07 | -0.7203 | 1.97E-07 |
| D+CA1 | Camk2b | -0.98162 | 2.57E-07 | -1.2909 | 3.23E-10 | -1.139 | 3.93E-11 |
| D+CA1 | Celf2 | -0.57061 | 0.007249 | -1.0785 | 3.47E-07 | -0.86191 | 8.21E-05 |
| D+CA1 | Chrm1 | -0.72545 | 6.33E-08 | -0.93789 | 7.11E-11 | -0.84953 | 1.89E-09 |
| D+CA1 | Cpne6 | -0.97212 | 1.82E-05 | -0.78967 | 0.00022903 | -0.90635 | 1.61E-04 |
| D+CA1 | Cttnbp2 | -1.0202 | 8.74E-06 | -1.1192 | 8.06E-08 | -1.0768 | 3.23E-11 |
| D+CA1 | Dlgap1 | -1.0202 | 0.0013434 | -1.2658 | 0.00010774 | -1.1576 | 3.67E-04 |
| D+CA1 | Eif5b | -0.74301 | 1.57E-07 | -0.49556 | 0.00016641 | -0.61913 | 3.14E-06 |
| D+CA1 | Enc1 | -0.55533 | 0.0042526 | -0.771 | 5.83E-05 | -0.67128 | 0.0015851 |
| D+CA1 | Epha4 | -0.53823 | 0.0055088 | -0.8215 | 3.10E-05 | -0.69141 | 0.0010319 |
| D+CA1 | Epha7 | -0.76187 | 2.21E-06 | -0.78226 | 1.38E-06 | -0.773 | 4.08E-08 |
| D+CA1 | Gabrb3 | -0.80567 | 0.00029572 | -1.3351 | 6.35E-06 | -1.1169 | 1.09E-04 |
| D+CA1 | Gda | -0.95569 | 2.29E-06 | -1.1175 | 8.76E-09 | -1.0406 | 1.52E-12 |
| D+CA1 | Gria2 | -0.60211 | 0.0091294 | -0.86912 | 0.0020729 | -0.75132 | 5.71E-03 |
| D+CA1 | Grin2a | -0.80116 | 3.03E-06 | -0.96684 | 6.79E-08 | -0.91889 | 4.22E-10 |
| D+CA1 | Icam5 | -0.93452 | 2.29E-117 | -0.55984 | 2.16E-63 | -0.73472 | 0 |
| D+CA1 | Kcnd2 | -0.51995 | 0.0071534 | -0.72916 | 0.00015409 | -0.62157 | 1.28E-03 |
| D+CA1 | Kcnj3 | -0.57204 | 0.0030766 | -0.6625 | 0.00045212 | -0.6031 | 0.0014029 |
| D+CA1 | Lppr4 | -0.53637 | 0.0045713 | -0.66768 | 0.000305 | -0.60667 | 0.002319 |
| D+CA1 | Lsamp | -0.4398 | 0.0091814 | -0.41741 | 0.0082966 | -0.44662 | 3.18E-04 |
| D+CA1 | Map1b | -0.62265 | 0.0015832 | -0.7223 | 0.00017728 | -0.69015 | 3.85E-04 |
| D+CA1 | Mbp | -0.54233 | 4.38E-05 | -0.41466 | 0.0049404 | -0.47487 | 5.88E-05 |
| D+CA1 | Miat | -0.50163 | 0.0030033 | -0.40952 | 0.0081873 | -0.4587 | 0.0001598 |
| D+CA1 | Nbea | -0.64709 | 0.0016029 | -0.51163 | 0.0080409 | -0.58191 | 0.0093151 |
| D+CA1 | Ndrg2 | -0.61451 | 0.0011866 | -0.62141 | 0.00059546 | -0.63262 | 5.75E-06 |
| D+CA1 | Nedd4l | -0.79059 | 4.44E-08 | -0.54183 | 5.32E-05 | -0.67836 | 7.78E-07 |
| D+CA1 | Nptx1 | -0.72888 | 1.43E-08 | -0.6796 | 7.76E-08 | -0.695 | 1.01E-07 |
| D+CA1 | Nrxn1 | -0.40707 | 0.0011784 | -0.46491 | 0.00027704 | -0.43087 | 2.42E-04 |
| D+CA1 | Olfm1 | -1.1805 | 5.94E-05 | -0.93608 | 0.0010094 | -1.0524 | 0.00028563 |
| D+CA1 | Plp1 | -0.76903 | 3.03E-17 | -0.51164 | 1.91E-10 | -0.63866 | 3.51E-13 |
| D+CA1 | Ppp3ca | -0.43595 | 9.45E-05 | -0.62243 | 1.99E-07 | -0.54941 | 2.97E-06 |
| D+CA1 | Ptgds | -1.8119 | 1.15E-07 | -0.87731 | 0.0037891 | -1.3212 | 4.24E-05 |
| D+CA1 | Rtn1 | -0.52915 | 0.0039628 | -0.47397 | 0.0057954 | -0.50729 | 0.0023641 |
| D+CA1 | Ryr2 | -0.67735 | 2.54E-06 | -0.48199 | 0.00041183 | -0.56312 | 8.09E-06 |
| D+CA1 | Scg2 | -0.47744 | 0.00032973 | -0.61852 | 8.57E-06 | -0.55502 | 1.20E-05 |
| D+CA1 | Sphkap | -0.90131 | 4.61E-05 | -1.2042 | 1.56E-05 | -1.0943 | 7.45E-05 |
| D+CA1 | Syn2 | -0.73247 | 2.38E-09 | -0.68868 | 1.19E-08 | -0.70628 | 8.15E-09 |
| D+CA1 | Ube2e2 | -1.2079 | 1.70E-07 | -0.85999 | 0.0020597 | -1.0251 | 0.00033346 |
| D+CA1 | Zeb2 | -0.7623 | 9.81E-09 | -0.88404 | 1.79E-10 | -0.83762 | 1.29E-09 |
| D+O | Malat1 | -0.44627 | 3.78E-07 | -0.42111 | 6.68E-07 | -0.42969 | 2.63E-07 |
| D+O | Pcdh9 | -1.5094 | 1.21E-13 | -0.98066 | 1.29E-08 | -1.3484 | 4.13E-11 |
| N+CA3 | Atp1b1 | -0.62451 | 0.0074442 | -0.78999 | 0.0018427 | -0.72355 | 0.0064454 |
| N+Ng | Nrg3 | -0.49794 | 1.19E-21 | -0.55269 | 0.0039019 | -0.52077 | 3.06E-289 |
| O+Ca | Nkain2 | -0.52773 | 2.55E-110 | -0.66 | 0.00072476 | -0.60164 | 0 |

* The full names of heterotypic pairs can be found in Table S5.
* The logFC was calculated based on the natural logarithm.
* In the case that fdr values were too small to be calucated, they were represented as zeros.

#### Table S2A. Commonly detected genes between CellNeighborEX and NicheNet for the the mouse embryo seqFISH data

| Centered cell type/neighboring cell type | Commonly detected |
|---|---|
| Gut-tube/Endothelium | Hoxc6 |
| Cardiomyocytes/Splanchnic-mesoderm | Cbfa2t3 |
| Gut-tube/Cranial-mesoderm | Sp5 |
| Forebrain-Midbrain-Hindbrain/Cranial-mesoderm | Kitl |
| Forebrain-Midbrain-Hindbrain/Definitive-endoderm | Kitl |
| Splanchnic-mesoderm/Cardiomyocytes | Hand2 |
| Surface-ectoderm/Mixed-mesenchymal-mesoderm | Hand2 |
| Gut-tube/Cranial-mesoderm | Snai1 |
| Surface-ectoderm/Neural-crest | Snai1 |
| Forebrain-Midbrain-Hindbrain/Definitive-endoderm | Nrp1 |
| Gut-tube/Splanchnic-mesoderm | Gata4 |
| Splanchnic-mesoderm/Cardiomyocytes | Aplnr |
| Cardiomyocytes/Splanchnic-mesoderm | Meis1 |
| Gut-tube/Splanchnic-mesoderm | Meis1 |
| Surface-ectoderm/Neural-crest | Tfap2b |
| Surface-ectoderm/Mixed-mesenchymal-mesoderm | Tgm1 |
| Cardiomyocytes/Splanchnic-mesoderm | Sfrp2 |
| Forebrain-Midbrain-Hindbrain/Spinal-cord | Sfrp2 |
| Lateral-plate-mesoderm/Intermediate-mesoderm | Sfrp2 |
| Spinal-cord/Forebrain-Midbrain-Hindbrain | Sfrp2 |
| Splanchnic-mesoderm/Gut-tube | Ets1 |
| Forebrain-Midbrain-Hindbrain/Cranial-mesoderm | Itga3 |
| Forebrain-Midbrain-Hindbrain/Definitive-endoderm | Itga3 |
| Splanchnic-mesoderm/Gut-tube | Itga3 |
| Gut-tube/Neural-crest | Postn |
| Forebrain-Midbrain-Hindbrain/Cranial-mesoderm | Shh |
| Forebrain-Midbrain-Hindbrain/Definitive-endoderm | Shh |
| Forebrain-Midbrain-Hindbrain/Neural-crest | Shh |
| Splanchnic-mesoderm/Gut-tube | Shh |
| Forebrain-Midbrain-Hindbrain/Cranial-mesoderm | Bmp7 |
| Forebrain-Midbrain-Hindbrain/Definitive-endoderm | Bmp7 |
| Gut-tube/Cranial-mesoderm | Bmp7 |
| Splanchnic-mesoderm/Cardiomyocytes | Bmp7 |
| Cardiomyocytes/Splanchnic-mesoderm | Meis2 |
| Gut-tube/Neural-crest | Meis2 |
| Gut-tube/Splanchnic-mesoderm | Meis2 |
| Spinal-cord/Forebrain-Midbrain-Hindbrain | Meis2 |
| Gut-tube/Splanchnic-mesoderm | Msx2 |
| Cardiomyocytes/Splanchnic-mesoderm | Lin28a |
| Cranial-mesoderm/Gut-tube | Lin28a |
| Gut-tube/Neural-crest | Lin28a |
| Gut-tube/Splanchnic-mesoderm | Lin28a |
| Gut-tube/Cranial-mesoderm | Fzd2 |
| Gut-tube/Neural-crest | Fzd2 |
| Gut-tube/Splanchnic-mesoderm | Fzd2 |
| Spinal-cord/Forebrain-Midbrain-Hindbrain | Fzd2 |
| Spinal-cord/Forebrain-Midbrain-Hindbrain | Sfrp1 |
| Gut-tube/Cranial-mesoderm | Mnt |
| Gut-tube/Neural-crest | Fli1 |
| Lateral-plate-mesoderm/Intermediate-mesoderm | Nr1d1 |
| Neural-crest/Surface-ectoderm | Cdh1 |
| Forebrain-Midbrain-Hindbrain/Definitive-endoderm | Gpc4 |
| Gut-tube/Neural-crest | Gpc4 |

| | |
|---|---|
| Gut-tube/Splanchnic-mesoderm | Osr1 |
| Cardiomyocytes/Endothelium | Pdgfra |
| Cardiomyocytes/Splanchnic-mesoderm | Pdgfra |
| Splanchnic-mesoderm/Cardiomyocytes | Pdgfra |
| Gut-tube/Neural-crest | Hoxa11 |
| Forebrain-Midbrain-Hindbrain/Definitive-endoderm | Etv4 |
| Splanchnic-mesoderm/Gut-tube | Etv4 |
| Forebrain-Midbrain-Hindbrain/Neural-crest | Hoxb5 |
| Gut-tube/Endothelium | Hoxb5 |
| Spinal-cord/Forebrain-Midbrain-Hindbrain | Hoxb5 |
| Gut-tube/Endothelium | Bmp4 |
| Splanchnic-mesoderm/Cardiomyocytes | Bmp4 |
| Cardiomyocytes/Endothelium | Acvr1 |
| Splanchnic-mesoderm/Gut-tube | Wnt3 |
| Forebrain-Midbrain-Hindbrain/Neural-crest | Tagln |
| Splanchnic-mesoderm/Cardiomyocytes | Tagln |
| Lateral-plate-mesoderm/Intermediate-mesoderm | Fgf3 |
| Gut-tube/Cranial-mesoderm | Chrd |
| Gut-tube/Splanchnic-mesoderm | Jarid2 |
| Lateral-plate-mesoderm/Intermediate-mesoderm | Wnt2 |
| Gut-tube/Splanchnic-mesoderm | Aldh1a2 |
| Presomitic-mesoderm/Dermomyotome | Aldh1a2 |
| Splanchnic-mesoderm/Gut-tube | Aldh1a2 |
| Gut-tube/Cranial-mesoderm | Col1a1 |
| Surface-ectoderm/Mixed-mesenchymal-mesoderm | Col1a1 |
| Gut-tube/Neural-crest | Efna5 |
| Spinal-cord/Forebrain-Midbrain-Hindbrain | Efna5 |
| Lateral-plate-mesoderm/Intermediate-mesoderm | Msx1 |
| Cardiomyocytes/Mixed-mesenchymal-mesoderm | Bmp2 |
| Dermomyotome/Presomitic-mesoderm | Hoxa7 |
| Lateral-plate-mesoderm/Intermediate-mesoderm | Icam2 |
| Lateral-plate-mesoderm/Intermediate-mesoderm | Wnt11 |
| Lateral-plate-mesoderm/Intermediate-mesoderm | Hoxb9 |
| Gut-tube/Neural-crest | Tfap2a |
| Surface-ectoderm/Mixed-mesenchymal-mesoderm | Tfap2a |
| Cardiomyocytes/Endothelium | Fst |
| Cardiomyocytes/Endothelium | Pdgfa |
| Cardiomyocytes/Splanchnic-mesoderm | Pdgfa |
| Gut-tube/Neural-crest | Pdgfa |
| Forebrain-Midbrain-Hindbrain/Cranial-mesoderm | En1 |
| Forebrain-Midbrain-Hindbrain/Definitive-endoderm | En1 |
| Lateral-plate-mesoderm/Intermediate-mesoderm | En1 |
| Forebrain-Midbrain-Hindbrain/Cranial-mesoderm | Foxa1 |
| Forebrain-Midbrain-Hindbrain/Definitive-endoderm | Foxa1 |
| Gut-tube/Cranial-mesoderm | Foxa1 |
| Gut-tube/Splanchnic-mesoderm | Foxa1 |
| Splanchnic-mesoderm/Gut-tube | Foxa1 |
| Neural-crest/Surface-ectoderm | Cldn4 |
| Splanchnic-mesoderm/Gut-tube | Cldn4 |
| Forebrain-Midbrain-Hindbrain/Cranial-mesoderm | Dnmt3a |
| Cardiomyocytes/Endothelium | Tbx5 |
| Cardiomyocytes/Splanchnic-mesoderm | Tbx5 |
| Gut-tube/Splanchnic-mesoderm | Tbx5 |
| Splanchnic-mesoderm/Gut-tube | Tbx5 |

| | |
|---|---|
| Lateral-plate-mesoderm/Intermediate-mesoderm | Col1a2 |
| Forebrain-Midbrain-Hindbrain/Cranial-mesoderm | Myh9 |
| Forebrain-Midbrain-Hindbrain/Definitive-endoderm | Myh9 |
| Splanchnic-mesoderm/Gut-tube | Myh9 |
| Dermomyotome/Presomitic-mesoderm | Lfng |
| Spinal-cord/Forebrain-Midbrain-Hindbrain | Lfng |
| Dermomyotome/Presomitic-mesoderm | Dll3 |
| Forebrain-Midbrain-Hindbrain/Spinal-cord | Dll3 |
| Spinal-cord/Forebrain-Midbrain-Hindbrain | Dll3 |
| Gut-tube/Endothelium | Hoxa9 |
| Forebrain-Midbrain-Hindbrain/Cranial-mesoderm | Gsn |
| Lateral-plate-mesoderm/Intermediate-mesoderm | Irx5 |
| Splanchnic-mesoderm/Cardiomyocytes | Irx5 |
| Splanchnic-mesoderm/Gut-tube | Irx5 |
| Dermomyotome/Presomitic-mesoderm | Lef1 |
| Forebrain-Midbrain-Hindbrain/Neural-crest | Lef1 |
| Gut-tube/Neural-crest | Lef1 |
| Dermomyotome/Presomitic-mesoderm | Cxcl12 |
| Splanchnic-mesoderm/Cardiomyocytes | Cxcl12 |
| Splanchnic-mesoderm/Gut-tube | Cxcl12 |
| Gut-tube/Splanchnic-mesoderm | Axin2 |
| Spinal-cord/Forebrain-Midbrain-Hindbrain | Axin2 |
| Cardiomyocytes/Endothelium | Cdh5 |
| Gut-tube/Endothelium | Cdh5 |
| Gut-tube/Cranial-mesoderm | Cdh2 |
| Cardiomyocytes/Splanchnic-mesoderm | Dusp6 |
| Forebrain-Midbrain-Hindbrain/Cranial-mesoderm | Dusp6 |
| Forebrain-Midbrain-Hindbrain/Definitive-endoderm | Dusp6 |
| Gut-tube/Neural-crest | Dusp6 |
| Splanchnic-mesoderm/Cardiomyocytes | Dusp6 |
| Forebrain-Midbrain-Hindbrain/Cranial-mesoderm | Hoxb3 |
| Forebrain-Midbrain-Hindbrain/Spinal-cord | Hoxb3 |
| Forebrain-Midbrain-Hindbrain/Definitive-endoderm | Apln |
| Forebrain-Midbrain-Hindbrain/Cranial-mesoderm | Igfbp3 |
| Forebrain-Midbrain-Hindbrain/Definitive-endoderm | Igfbp3 |
| Cardiomyocytes/Endothelium | Igf1 |
| Gut-tube/Cranial-mesoderm | Igf1 |
| Surface-ectoderm/Mixed-mesenchymal-mesoderm | Ahnak |
| Gut-tube/Neural-crest | Dlk1 |
| Lateral-plate-mesoderm/Intermediate-mesoderm | Dlk1 |
| Forebrain-Midbrain-Hindbrain/Spinal-cord | Fgfr3 |
| Spinal-cord/Forebrain-Midbrain-Hindbrain | Fgfr3 |
| Gut-tube/Endothelium | Hoxa13 |
| Lateral-plate-mesoderm/Intermediate-mesoderm | Apob |
| Gut-tube/Neural-crest | Bid |
| Splanchnic-mesoderm/Gut-tube | Bid |
| Forebrain-Midbrain-Hindbrain/Definitive-endoderm | Wnt5a |
| Gut-tube/Endothelium | Wnt5a |
| Splanchnic-mesoderm/Cardiomyocytes | Wnt5a |
| Splanchnic-mesoderm/Gut-tube | Wnt5a |
| Forebrain-Midbrain-Hindbrain/Cranial-mesoderm | Foxa2 |
| Forebrain-Midbrain-Hindbrain/Definitive-endoderm | Foxa2 |
| Splanchnic-mesoderm/Gut-tube | Foxa2 |
| Gut-tube/Endothelium | Tbx3 |

| | |
|---|---|
| Gut-tube/Cranial-mesoderm | Nr2f1 |
| Spinal-cord/Forebrain-Midbrain-Hindbrain | Nr2f1 |
| Surface-ectoderm/Neural-crest | Nr2f1 |
| Gut-tube/Splanchnic-mesoderm | Hoxb1 |
| Cardiomyocytes/Splanchnic-mesoderm | Fgfr2 |
| Cranial-mesoderm/Gut-tube | Fgfr2 |
| Gut-tube/Cranial-mesoderm | Fgfr2 |
| Gut-tube/Splanchnic-mesoderm | Fgfr2 |
| Lateral-plate-mesoderm/Intermediate-mesoderm | Fgfr2 |
| Forebrain-Midbrain-Hindbrain/Cranial-mesoderm | Sox2 |
| Gut-tube/Cranial-mesoderm | Sox2 |
| Gut-tube/Neural-crest | Sox2 |
| Splanchnic-mesoderm/Gut-tube | Sox2 |

**Table S2B. Commonly detected genes between CellNeighborEX and NicheNet for the homotypic spots of the mouse embryo Slide-seq data**

| Heterotypic pair | Commonly detected |
|---|---|
| En+L | Sfrp1 |

**Table S2C. Commonly detected genes between CellNeighborEX and NicheNet for the homotypic spots of the mouse liver cancer Slide-seq data**

| Heterotypic pair | Commonly detected |
|---|---|
| HepatocyteI+HepatocyteII | Arg1 |
| VSMC+TumorIII | Col1a2 |
| VSMC+TumorIII | Col3a1 |
| TumorIII+Monocyte | F13a1 |
| HepatocyteI+HepatocyteII | Igf1 |
| HepatocyteI+VSMC | Lcn2 |
| HepatocyteI+Monocyte | Marco |
| HepatocyteI+HepatocyteII | Rnase4 |
| HepatocyteI+Monocyte | Slc40a1 |
| HepatocyteII+Monocyte | Slc40a1 |
| TumorII+HepatocyteI | Spp1 |
| TumorIII+HepatocyteI | Spp1 |

**Table S2D. Commonly detected genes between CellNeighborEX and NicheNet for the homotypic spots of the mouse hippocampus Slide-seq data**

| Heterotypic pair | Commonly detected |
|---|---|
| Ep+Ca | Ptgds |
| In+Ca | Ptgds |
| In+D | Cck |
| In+Ento | Cck |
| In+Ento | Gabbr1 |
| In+Ento | Gabbr2 |
| In+Ento | Grm1 |
| In+Ento | Rgs4 |
| In+Ento | Ntng1 |
| In+Ento | Nrxn1 |
| In+Ep | Gabbr1 |

**Table S3A. Commonly detected genes between CellNeighborEX and NicheNet for the heterotypic spots of the mouse embryo Slide-seq data**

| Sender | Receiver | Up-regulated gene | Type |
|---|---|---|---|
| En+L | En+L | Ednrb | target gene |
| En+L | En+L | Slc2a1 | target gene |

**Table S3B. Commonly detected genes between CellNeighborEX and NicheNet for the heterotypic spots of the mouse liver cancer Slide-seq data**

| Sender | Receiver | Up-regulated gene | Type |
|---|---|---|---|
| HepatocyteI+HepatocyteII | HepatocyteI+HepatocyteII | Igf1 | target |
| HepatocyteI+Monocyte | HepatocyteI+Monocyte | Cd5l | target |
| HepatocyteI+Monocyte | HepatocyteI+Monocyte | Marco | target |
| HepatocyteI+VSMC | HepatocyteI+VSMC | Lcn2 | target |
| HepatocyteI+HepatocyteII | HepatocyteI+HepatocyteII | Arg1 | target |
| HepatocyteI+HepatocyteII | HepatocyteI+HepatocyteII | Rnase4 | target |
| HepatocyteII+Monocyte | HepatocyteII+Monocyte | Alcam | ligand |
| VSMC+TumorIII | VSMC+TumorIII | Col1a2 | target |
| HepatocyteI+Monocyte | HepatocyteI+Monocyte | Slc40a1 | receptor/target gene |
| HepatocyteII+Monocyte | HepatocyteII+Monocyte | Slc40a1 | receptor/target gene |
| TumorIII+Monocyte | TumorIII+Monocyte | F13a1 | target |
| VSMC+TumorIII | VSMC+TumorIII | Col3a1 | target |
| TumorIII+HepatocyteII | TumorIII+HepatocyteII | Tgfbr1 | receptor/target gene |
| TumorII+HepatocyteI | TumorII+HepatocyteI | Spp1 | ligand/target gene |
| TumorIII+HepatocyteI | TumorIII+HepatocyteI | Spp1 | ligand/target gene |

**Table S3C. Commonly detected genes between CellNeighborEX and NicheNet for the heterotypic spots of the mouse hippocampus Slide-seq data**

| Sender | Receiver | Up-regulated gene | Type |
|---|---|---|---|
| CA3+D | CA3+D | Igfbp4 | target gene |
| EnT+In | EnT+In | Agt | ligand/target gene |
| EnT+In | EnT+In | Mbp | target gene |
| EnT+In | EnT+In | Plp1 | target gene |
| EnT+In | EnT+In | Tcf7l2 | target gene |
| Ep+O | Ep+O | Zbtb20 | target gene |
| In+Ca | In+Ca | Ptgds | target gene |
| In+D | In+D | Cck | ligand |
| In+Ento | In+Ento | Cck | ligand |
| In+Ep | In+Ep | Cpne9 | target gene |
| In+Ep | In+Ep | Zic4 | target gene |

**Table S4A. Cell types and their marker genes in the mouse embryo scRNA-seq data**

| Cluster | Cell type | Abbrev | Marker1 | Marker2 |
|---|---|---|---|---|
| 1 | Connective tissue progenitors | CTP | Tgfb2 | Postn |
| 2 | Chondrocytes & osteoblasts | CO | Twist2 | Prrx1 |
| 3 | Intermediate Mesoderm | IM | Mylk | Ednra |
| 4 | Jaw and tooth progenitors | JTP | Sox9 | Col2a1 |
| 5 | Excitatory neurons | EN | Car10 | Ntng1 |
| 6 | Epithelial cells | Ept | Epcam | Trp63 |
| 7 | Radial glia | Rg | Fabp7 | Pax3 |
| 8 | Early mesenchyme | Em | Gpc5 | Smoc1 |
| 9 | Neural progenitor cells | NP | Prmt8 | Gadd45g |
| 10 | Postmitotic premature neurons | PPN | Nkx6-3 | Nrn1 |
| 11 | Oligodendrocyte Progenitors | OP | Slc17a6 | Sox1 |
| 12 | Isthmic organizer cells | IO | Fgf15 | - |
| 13 | Myocytes | My | Myh3 | Neb |
| 14 | Neural Tube | NT | Foxb1 | Scube2 |
| 15 | Inhibitory neurons | In | Pax2 | Slc6a5 |
| 16 | Stromal cells | Sc | Bmpr1a | Rab5a |
| 17 | Osteoblasts | O | Col1a1 | Camk1d |
| 18 | Inhibitory neuron progenitors | Inp | Slc6a5 | Pax2 |
| 19 | Premature oligodendrocyte | PO | Pcdh19 | - |
| 20 | Endothelial cells | En | Pecam1 | Egfl7 |
| 21 | Chondrocyte progenitors | CP | Itga11 | Atp1a2 |
| 22 | Definitive erythroid lineage | DEL | Slc4a1 | Kel |
| 23 | Schwann cell precursor | SCP | Plp1 | Cdh19 |
| 24 | Sensory neurons | SN | Syt13 | Shox2 |
| 25 | Limb mesenchyme | LM | Msx1 | Lmx1b |
| 26 | Primitive erythroid lineage | PEL | Hba-a1 | Hbb-y |
| 27 | Inhibitory interneurons | II | Dlx1 | - |
| 28 | Granule neurons | Gn | NeuroD2 | Tiam2 |
| 29 | Hepatocytes | H | Afp | Alb |
| 30 | Notochord cells | Nc | Shh | Slit1 |
| 31 | White blood cells | WB | Apoe | Ctss |
| 32 | Ependymal cell | Epn | Sostdc1 | Htr2c |
| 33 | Cholinergic neurons | Cn | Slit2 | Slit3 |
| 34 | Cardiac muscle lineages | Cml | Myl2 | Myocd |
| 35 | Megakaryocytes | Mg | Pf4 | Itgb3 |
| 36 | Melanocytes | Ml | Trpm1 | Pmel |
| 37 | Lens | L | Cryba1 | Cryaa |

**Table S4B. Cell types and their marker genes in the mouse liver cancer snRNA-seq data**

| Cluster | Cell type | Marker |
|---|---|---|
| 0 | Tumor I | Hmga2 |
| 1 | Hepatocyte II | Reln |
| 2 | Monocyte | Trbc1 |
| 3 | LSEC | Ccr2 |
| 4 | Tumor II | Hmga2 |
| 5 | Hepatocyte I | Fmo5 |
| 6 | Tumor III | Aqp5 |
| 7 | Interferon | Ifit3 |
| 8 | T | Ly6d |
| 9 | VSMC | Egfl7 |
| 10 | Kupffer/Monocyte | Ctss |
| 11 | B | Ptprc(CD45) |
| 12 | Ribosomal+ | Clec4f |
| 13 | Hepatic Stellate Cell | Adamts2 |

**Table S4C. Cell types and their marker genes in the mouse hippocampus scRNA-seq data**

| Cluster | Cell type | Abbrev | Marker1 | Marker2 |
|---|---|---|---|---|
| 1 | Interneuron | In | Sst | Gad1 |
| 2 | Neuron.Slc17a6 | N | Nefm | Nell1 |
| 3 | Entorhinal | Ento | Slc17a7 | Ajap1 |
| 4 | Dentate | D | C1ql2 | Penk |
| 5 | CA1 | CA1 | Grm5 | Lypd1 |
| 6 | CA3 | CA3 | Slc17a7 | Pvrl3 |
| 7 | Astrocyte | A | Gja1 | Fabp7 |
| 8 | Oligodendrocyte | O | Trf | Il33 |
| 9 | Polydendrocyte | P | Tnr | Ptprz1 |
| 10 | Microglia_Macrophages | MM | C1qb | Tmem119 |
| 11 | Ependymal | Ep | Ccdc153 | - |
| 12 | Choroid | Ch | Ttr | - |
| 13 | Neurogenesis | Ng | Sox4 | Efhd2 |
| 14 | Cajal_Retzius | Ca | Slc17a6 | - |
| 15 | Endothelial_Stalk | EnS | Flt1 | Cdkn2b |
| 16 | Mural | M | Acta2 | Kcnj8 |
| 17 | Endothelial_Tip | EnT | Dcn | Mgp |

#### Table S5A. Comparison of expression origin between regression and single cell data in the heterotypic spots of the mouse embryo Slide-seq data

| Heterotypic pair | Up-regulated_gene | Expression in scRNA-seq | Expression in regression | Match | p-value in regression |
|---|---|---|---|---|---|
| DEL+PEL | Hbb-bt | PEL | PEL | 1 | 1.43E-09 |
| DEL+PEL | Hbb-bs | PEL | PEL | 1 | 2.48E-07 |
| DEL+PEL | Hba-a1 | PEL | PEL | 1 | 0.00031519 |
| LM+PEL | Cdkn1c | PEL | PEL | 1 | 0.0042853 |
| En+L | Cd24a | En | En | 1 | 2.53E-02 |
| En+L | Tmem40 | En | L | 0 | 0.017969 |
| En+L | Etl4 | L | En | 0 | 0.048494 |

#### Table S5B. Comparison of expression origin between regression and single nucleus data in the heterotypic spots of the mouse liver cancer Slide-seq data

| Heterotypic pair | Up-regulated_gene | Expression in snRNA-seq | Expression in regression | Match | p-value in regression |
|---|---|---|---|---|---|
| HepatocyteI+Monocyte | Slc40a1 | Monocyte | Monocyte | 1 | 5.57E-14 |
| HepatocyteI+HepatocyteII | Serpina1c | HepatocyteI | HepatocyteI | 1 | 3.17E-13 |
| HepatocyteI+Monocyte | Clec4f | Monocyte | Monocyte | 1 | 2.66E-11 |
| HepatocyteI+Monocyte | Cd163 | Monocyte | Monocyte | 1 | 1.29E-10 |
| HepatocyteI+Monocyte | Marco | Monocyte | Monocyte | 1 | 6.44E-09 |
| HepatocyteI+Monocyte | Cd5l | Monocyte | Monocyte | 1 | 2.40E-07 |
| HepatocyteI+HepatocyteII | Hsd17b13 | HepatocyteII | HepatocyteII | 1 | 2.55E-06 |
| TumorIII+Monocyte | F13a1 | Monocyte | Monocyte | 1 | 3.93E-06 |
| HepatocyteI+HepatocyteII | Rnase4 | HepatocyteI | HepatocyteI | 1 | 4.62E-06 |
| HepatocyteII+Monocyte | Clec4f | Monocyte | Monocyte | 1 | 7.43E-05 |
| HepatocyteI+HepatocyteII | Saa4 | HepatocyteII | HepatocyteII | 1 | 0.00054523 |
| HepatocyteII+Monocyte | Slc40a1 | Monocyte | Monocyte | 1 | 0.0017785 |
| HepatocyteI+HepatocyteII | Itih4 | HepatocyteI | HepatocyteI | 1 | 0.023217 |
| VSMC+TumorIII | Col3a1 | VSMC | VSMC | 1 | 0.032209 |
| TumorIII+HepatocyteII | Ext1 | TumorIII | TumorIII | 1 | 0.043759 |
| HepatocyteII+Monocyte | Vti1a | Monocyte | Monocyte | 1 | 0.045122 |
| HepatocyteI+HepatocyteII | Arg1 | HepatocyteII | HepatocyteI | 0 | 5.05E-07 |
| HepatocyteI+HepatocyteII | Tdo2 | HepatocyteII | HepatocyteI | 0 | 0.000255 |
| HepatocyteII+Monocyte | Hsd17b13 | HepatocyteII | Monocyte | 0 | 0.012494 |

**Table S5C. Comparison of expression origin between regression and single cell data in the heterotypic spots of the mouse hippocampus Slide-seq data**

| Heterotypic pair | Up-regulated_gene | Expression in scRNA-seq | Expression in regression | Match | p-value in regression |
|---|---|---|---|---|---|
| In+A | Mbp | A | A | 1 | 1.08E-39 |
| In+A | Plp1 | A | A | 1 | 7.91E-35 |
| Ento+Ch | Meg3 | Ento | Ento | 1 | 1.95E-15 |
| In+Ep | Meg3 | In | In | 1 | 3.00E-11 |
| Ento+Ch | Plp1 | Ento | Ento | 1 | 3.41E-07 |
| A+O | Dbi | A | A | 1 | 1.20E-06 |
| Ento+Ch | Nwd2 | Ento | Ento | 1 | 1.73E-06 |
| In+Ca | Zic1 | Ca | Ca | 1 | 5.04E-06 |
| Ento+Ch | Calb2 | Ento | Ento | 1 | 1.34E-05 |
| Ento+Ch | Zcchc12 | Ento | Ento | 1 | 1.98E-05 |
| In+Ep | Snhg11 | In | In | 1 | 6.83E-05 |
| In+Ep | Ttr | Ep | Ep | 1 | 0.00047689 |
| In+O | Ndrg2 | O | O | 1 | 0.00056087 |
| In+MM | Plp1 | MM | MM | 1 | 0.00079119 |
| Ento+Ch | Nnat | Ento | Ento | 1 | 0.0020071 |
| Ento+Ch | Tac2 | Ento | Ento | 1 | 0.0023071 |
| Ep+N | Calb2 | N | N | 1 | 0.0069948 |
| In+Ep | Kcnma1 | In | In | 1 | 0.0071623 |
| In+Ep | Gabbr1 | In | In | 1 | 0.0076462 |
| In+Ento | Ntng1 | In | In | 1 | 0.0092721 |
| In+Ca | Gucy1a3 | Ca | Ca | 1 | 0.0098233 |
| Ento+Ch | Gm5741 | Ento | Ento | 1 | 0.011658 |
| In+Ento | Ramp3 | Ento | Ento | 1 | 0.017423 |
| In+Ep | Pnmal2 | In | In | 1 | 0.020333 |
| Ento+Ch | Malat1 | Ento | Ento | 1 | 0.020804 |
| In+Ep | Pcp4 | In | In | 1 | 0.022588 |
| EnT+Ento | Necab2 | Ento | Ento | 1 | 0.023669 |
| Ep+Ca | Ptgds | Ca | Ca | 1 | 0.031829 |
| In+Ento | Meg3 | In | In | 1 | 0.037265 |
| In+Ep | Kcnip1 | In | In | 1 | 0.03909 |
| In+Ca | Ptgds | Ca | Ca | 1 | 0.044691 |
| Ento+Ch | Zic1 | Ch | Ento | 0 | 4.11E-06 |
| In+Ca | Cygb | In | Ca | 0 | 5.45E-06 |
| Ento+Ch | Necab2 | Ch | Ento | 0 | 3.79E-05 |
| N+O | Pcp4 | N | O | 0 | 0.00021799 |
| Ento+Ch | Msi2 | Ch | Ento | 0 | 0.0028733 |
| Ch+EnT | Igfbp7 | EnT | Ch | 0 | 0.010201 |
| In+Ento | Gabbr1 | In | Ento | 0 | 0.010334 |
| O+P | Slc44a1 | O | P | 0 | 0.016222 |
| Ch+EnT | Lum | EnT | Ch | 0 | 0.025337 |
| Ento+Ch | Aifm3 | Ch | Ento | 0 | 0.031357 |
| In+Ento | Cck | In | Ento | 0 | 0.038743 |
| Ento+Ch | Apod | Ch | Ento | 0 | 0.046048 |
